# Ectomycorrhizal symbiosis evolved independently and by convergent gene duplication in rosid lineages

**DOI:** 10.1101/2024.10.21.618687

**Authors:** Fabian van Beveren, Yvet Boele, Camille Puginier, Matheus Bianconi, Cyril Libourel, Maxime Bonhomme, Jean Keller, Pierre-Marc Delaux

**Affiliations:** Laboratoire de Recherche en Sciences Végétales (LRSV), Université de Toulouse, CNRS, UPS, INP, Toulouse, 31320, Castanet-Tolosan, France; Laboratory of Cell and Developmental Biology, Wageningen University, Droevendaalsesteeg 1, 6708 PB Wageningen, the Netherlands; Department of Terrestrial Ecology, Netherlands Institute of Ecology (NIOO-KNAW), Droevendaalsesteeg 10, 6708 PB Wageningen, the Netherlands

## Abstract

Many land plants rely on mutualistic symbiotic associations to thrive, starting with their common ancestor associating with arbuscular mycorrhizal (AM) fungi. Similar to AM symbiosis, multiple other intracellular symbiotic interactions have evolved in a variety of land plants. Ectomycorrhizal (ECM) symbiosis contrasts with these relationships as there is no intracellular accommodation of the symbiont inside the plant cell, but only intercellular colonization. This symbiotic relationship is common in a variety of seed plants, mostly trees and shrubs such as pines, willows and oaks.

Although it is known which plant lineages are involved in ECM symbiosis, there has been little investigation into the evolutionary origin of ECM symbiosis in these lineages. Furthermore, the genetic innovations and transcriptomic response related to this symbiosis have been studied largely in a genus or species-specific context, which hinders the study of their evolutionary origins. In this study, we reconstruct the origin of ECM in the rosid clade, showing at least 16 independent origins, resulting in the 17 known extant ECM rosid lineages. Moreover, comparative genomics of these lineages highlight genes involved in cell wall remodeling which underwent duplications in a convergent manner across ectomycorrhizal lineages.

## Introduction

Many land plants rely on mutualistic symbiotic associations to thrive, starting with their common ancestor associating with arbuscular mycorrhizal (AM) fungi (Rich *et al*., 2021). Similar to AM symbiosis, multiple other intracellular symbiotic interactions have evolved in a variety of land plants, such as ericoid symbiosis in Ericaceae and root nodule symbiosis with nitrogen-fixing bacteria in the legumes and relatives (Strullu-Derrien *et al*., 2018; Radhakrishnan *et al*., 2020). Ectomycorrhizal (ECM) symbiosis contrasts with these relationships as there is no intracellular accommodation of the symbiont inside the plant cell, but only intercellular colonization (Brundrett & Tedersoo, 2018). This symbiotic relationship is common in a variety of seed plants, mostly trees and shrubs such as pines, willows and oaks (Cairney, 2000; Tedersoo & Brundrett, 2017).

Effort has been put to uncover the fungal side of this symbiotic relationship (Kohler *et al*., 2015), but the plant side remains more elusive. Although it is known which plant lineages are involved in ECM symbiosis (Tedersoo & Brundrett, 2017), there has been little investigation into the evolutionary origin of ECM symbiosis in these lineages. Furthermore, the genetic innovations and transcriptomic response related to this symbiosis have been studied largely in a genus or species-specific context (Liao *et al*., 2016; Bouffaud *et al*., 2020; Chowdhury *et al*., 2022; Hill *et al*., 2022), which hinders the study of their evolutionary origins. In this study, we reconstruct the origin of ECM in the rosid clade, showing at least 16 independent origins, resulting in the 17 known extant ECM rosid lineages. Moreover, comparative genomics of these lineages highlight genes involved in cell wall remodeling which underwent duplications in a convergent manner across ectomycorrhizal lineages.

## Results and Discussion

### Ancestral state reconstruction supports multiple independent gains of ECM in angiosperms

The many different ECM lineages have been proposed to have originated independently (Brundrett, 2009; Tedersoo & Smith, 2013). In rosids, the main ECM lineages are Fagales, Salicaceae (Malpighiales) and Myrtoideae (Myrtales), but also include less species-diverse clades in the orders Fabales, Rosales, Malpighiales and Malvales (Tedersoo & Brundrett, 2017). We reconstructed a phylogenetic tree of rosids based on protein sequences of *RbcL* and *MatK*, to model the evolution of the ECM character. Using the best-performing model, we found a largely independent origin of ECM lineages, in agreement with the literature (Fig. 1). A clear single origin in the common ancestor of Fagales can be detected, with a loss in the Myricaceae, a group that instead associates with nitrogen-fixing bacteria of the genus *Frankia* (Huguet *et al*., 2005). ECM symbiosis in Myrtoideae has a clear single origin with a putative single loss in the common ancestor of *Pimenta* and *Myrtus*, both members of Myrteae. This single loss agrees with the literature (Tedersoo & Brundrett, 2017), although additional losses have been suggested that can only be recovered with a denser taxon sampling of this clade (Thornhill *et al*., 2015; Tedersoo & Brundrett, 2017).

**Fig. 1:**
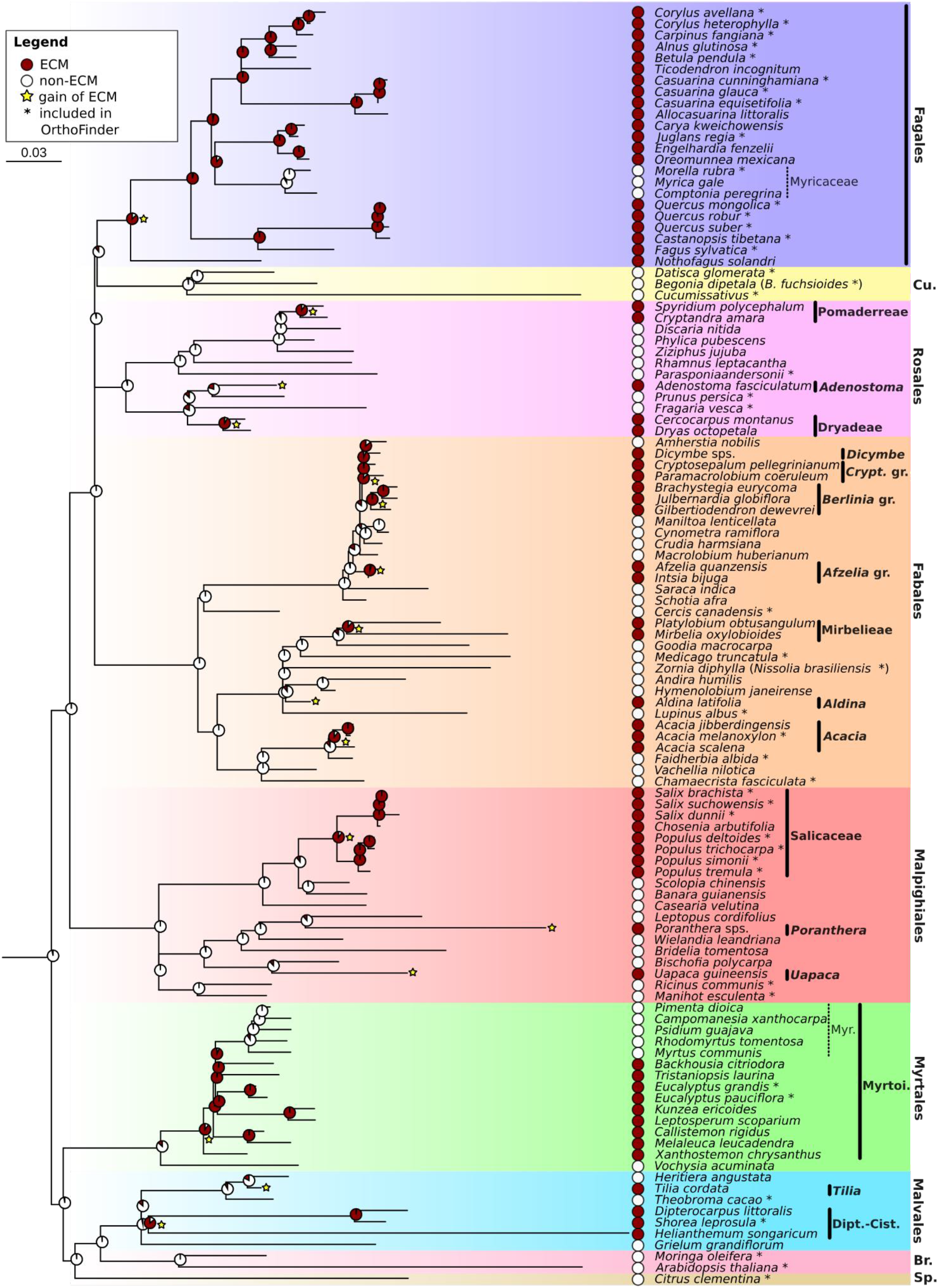
Ancestral reconstruction of ectomycorrhizal symbiosis in plant lineages of rosids. Phylogenetic tree was reconstructed based on plastid protein sequences of RbcL and MatK, reconstructed under the model Q.plant+C60+F0+R5. Ancestral reconstruction is based on the ARD+kappa model. Likelihood of the ECM (red) or non-ECM (white) status is shown proportionally for each node as a pie diagram, whereas status of included rosid species is shown next to the species name. The 17 rosid ECM clades identified by Tedersoo and Brundrett (2017) are shown with thick black lines and their names and the 16 yellow stars represent independent origins of ECM symbiosis. The two clades represented by multiple species that lost the ability for ECM symbiosis are shown with dashed lines. Asterisks denote species also included in the OrthoFinder analysis and the species name is placed between parentheses if a different species was used. Scale bar represents estimated substitutions per site. Cu. = Cucurbitales, Crypt. gr. = Cryptosepalum group, Myr. = Myrteae, Myrtoi. = Myrtoideae, Dipt.-Cist. = Dipterocarpaceae-Cistacea, Br. = Brassicales, Sp. = Sapindales.

Tedersoo and Brundrett (2017) separate four ECM lineages within the Detarioideae (Fabales; Fig. 1), two of these we estimate to have originated independently: the Afzelia group (*Afzelia* and *Intsia*) and the Berlinia group (here represented by *Brachystegia, Gilbertiodendron* and *Julbernardia*). We detect a single origin in the common ancestor of the other two groups (*Dicymbe* and *Cryptosepalum*+*Paramacrolobium*), followed by a loss in *Amherstia* (Fig. 1). Other genera are more closely related to *Dicymbe* than *Amherstia*, which may bias our results (de la Estrella *et al*., 2017). The relationships among lineages of Detarioideae remain uncertain, and more data besides *RbcL* and *MatK* markers might be necessary to resolve their phylogeny. All other rosid ECM lineages are shown to have evolved the symbiosis independently (Fig. 1). Overall, our reconstruction of ancestral states within the rosids shows that the identified ECM lineages largely evolved this symbiotic association independently.

As ancestral state reconstruction is limited by phylogenetic signal, taxon sampling and limited models for character evolution, we decided to also look for genetic signal that may indicate an ancestral origin of ECM in rosids. With a database of 63 genomes, including 25 ECM and 19 non-ECM rosids (Supplementary table S1), we reconstructed orthologous groups associated to the common ancestral node of rosids, and mined for orthogroups being largely absent in the non-ECM rosids (Supplementary Methods S1). These groups would reflect convergent gene losses in these lineages which could be explained by co-elimination following the loss of ECM, a phenomenon observed following the loss of other symbioses (Delaux *et al*., 2014; Bravo *et al*., 2016; Griesmann *et al*., 2018). After manual curation, we identified six genes that share an origin in the common ancestor of rosids, but have been retained mostly by ECM species, although they remain present in several genomes of non-ECM species (Fig. 2, Supplementary Figs. S1-S6). Part of the reason may be that the pressure to lose them is strengthened in non-ECM lineages, but not enough to result in a complete loss in these lineages. It may also be that the loss of ECM is recent in some of these lineages. For example, a specific short-chain dehydrogenase/reductase (SDR) gene was retained in non-ECM Rosales, but Rosales contains several ECM-lineages, thus the presence of this gene could be explained by the recent loss of ECM (Fig. 2). It is however difficult to tell how many genes are expected to follow such a pattern by chance, with retention only in a small subset of lineages. In addition, none of these genes are conserved in all the ECM lineages covered by our analysis.

**Fig. 2:**
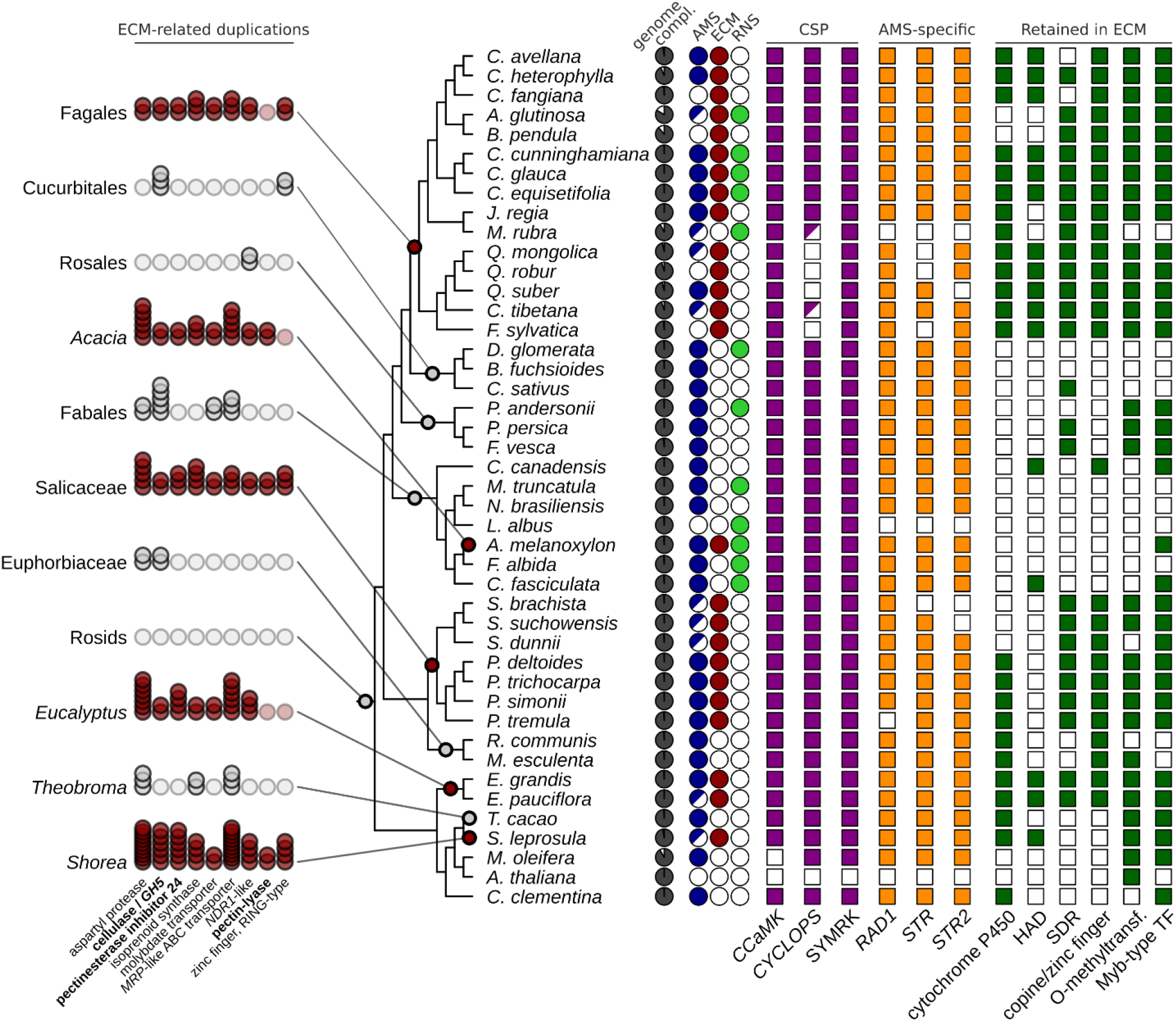
Cladogram of rosid species included in the orthogroup analysis, depicting ECM-specific gene duplications and retainment of genes with a known or putative role in symbiosis. Phylogenetic relationships are based on Figure 1. Duplications are shown for 11 taxonomic groups. A single faint circle means no duplications occurred, whereas two or multiple circles represent one or more duplications, and the circles are colored by the ECM (red) or non-ECM (gray) status of that node. Presence (colored) or absence (white) are shown for genes part of the common signaling pathway (CSP), specific to arbuscular mycorrhizal symbiosis (AMS) and those retained in ECM lineages. When the AMS state of a species is uncertain the circle is half-colored. Genes partially present are shown by a half-colored square. Duplicated genes related to cell wall remodeling are shown in bold. Abbreviations: compl. = completeness, HAD = haloacid dehalogenase-like, SDR = short-chain dehydrogenase/reductase, O-methylfransf. = O-methyltransferase, TF = transcription factor.

Overall, the retention of these genes does not clearly support an ancestral origin of this symbiosis, and in light of the ancestral state reconstruction is still best explained by multiple independent origins of ECM in rosids.

### ECM evolution is linked with convergent gene duplications

The independent emergence of traits may result either from completely different or from convergent genetic pathways. Convergent evolution occurs through the co-option of existing genes or following gene-duplication and may result in similar molecular responses (Satterlee *et al*., 2024).

First, we reasoned that convergence towards the ability to engage in ECM symbiosis should be detected at the transcriptomic level, thus we looked for shared transcriptomic responses to ECM in three host species: *Quercus robur, Populus tremula × P. tremuloides* and *Eucalyptus grandis* (Bouffaud *et al*., 2020; Chowdhury *et al*., 2022; Hill *et al*., 2022). Comparing the overlap in deregulation of genes during ECM, we find a set of 537 orthogroups that are deregulated in all three species (Supplementary Fig. S7). This number of shared differentially expressed orthogroups is higher than would be expected by chance (mean: 359, p-value: 0.00001), which is especially notable considering different symbionts and time points tested in the three studies included in this analysis. This shows that at least from a transcriptomic perspective, there is a similar response to the presence of ECM fungi.

Then, we investigated the potential contribution of gene duplication to the convergent evolution of ECM. To do so, we looked into gene duplication events that have occurred independently in the ancestral nodes of the five ECM rosid lineages that were covered in our genome analysis: Fagales, Salicaceae, Myrtoideae, Dipterocarpaceae-Cistacea and *Acacia*. Using OrthoFinder, we detected 61 orthogroups with duplication events in the groups represented by multiple genera: the common ancestors of the Salicaceae and the one of the Fagales. Among these 61 orthogroups, a number are overall frequently duplicated across the whole phylogeny. To further identify relevant candidates in this list, we selected the ones that matched one of the following three additional criteria: (1) at least one of the duplicated genes is deregulated in the three ECM species with available RNAseq data, (2) a high proportion (> 1/7) of duplication co-occur with the gain of ECM, or (3) more than half of the duplication occurred in any ECM node. This filter resulted in the shortlisting of 27 orthogroups, which after manual curation through single-gene phylogenetic analyses resulted in a final selection of nine genes (Supplementary Figs. S8-S16). These genes are duplicated in a minimum of three ECM origin nodes and not frequently duplicated in a set of non-ECM nodes (Fig. 2; supplementary table S2).

Strikingly, three out of the nine candidate genes have functions associated with cell-wall remodeling: a Pectinesterase inhibitor, a Pectin methylesterase and a Cellulase (Fig. 2). Modification of the plant cell wall is a common feature in intra- and intercellular plant -fungi mutualistic symbioses (Su, 2023). In particular, extensive pectin modifications have been observed during the interaction between ECM fungi and their hosts, and part of these modifications are mediated by the fungal partner (Chowdhury *et al*., 2022). However, comparative phylogenomics of the fungal symbionts have revealed that ECM fungal lineages have a reduced set of plant cell wall degrading enzymes, a genomic reduction that occurred in a convergent manner (Kohler *et al*., 2015). The convergent expansions of gene families encoding for cell-wall modifying enzymes in plant lineages that evolved the ability to engage in ECM symbiosis might reflect a shift in the distribution of this functionality from the symbiont to the host which may represent a common principle for the evolution of ECM symbiosis.

### Convergent co-option of the common symbiosis pathway for ECM may have occurred in rosids

Besides gene duplication, convergent evolution originates from co-option of existing genes and pathways. In the context of ECM, it has been proposed that its evolution in Fagales was facilitated by the co-option of the common signaling pathway (CSP; Li et al. 2024), a module of several proteins essential for the establishment of intracellular symbiosis (Parniske, 2008). This hypothesis, supported by reverse genetic analyses in poplar (Cope *et al*., 2019), is based on the conservation of those genes in plant species forming ECM which have, in parallel, lost the ability to do any form of intracellular symbioses. Indeed, the loss of this ability has been correlated in many plant lineages with the loss of associated genes by relaxed selection (Delaux *et al*., 2014; Bravo *et al*., 2016; Radhakrishnan *et al*., 2020). Thus, the retention of these genes may reflect the evolution of another selection pressure, possibly by ECM (Li *et al*., 2024). Similarly, genes have been found to be specifically dedicated to the ability to engage in arbuscular mycorrhizal symbiosis (AMS). The retention of one of such genes in ECM Fagales, the Glycerol 3-Phosphate Acyltransferase RAM2, an enzyme essential for the functioning of the AMS (Wang *et al*., 2012), also indicates that it might have been co-opted for ECM in that lineage (Li *et al*., 2024).

To determine whether the convergent evolution of ECM relied more broadly on the co-option of these genes, we conducted targeted phylogenies on the CSP genes *SYMRK, CCaMK* and *CYCLOPS*, and on the AMS-specific genes *RAD1, STR* and *STR2* (Supplementary Figs. S17-S22). Only three of the included genomes are lacking *RAD1, STR1* and *STR2*. Two of these are *Arabidopsis thaliana* and *Lupinus albus*, both known to lack mycorrhizal (both AM and ECM) symbioses. The other is *Morella rubra*, which has multiple observations of AM listed on the FungalRoot database (Soudzilovskaia *et al*., 2020, 2024). Moreover, other members of the genus *Myrica*/*Morella* are considered to be both ECM and AMS species (Urgiles *et al*., 2014; Teste *et al*., 2020). It is possible that the loss of AMS is species-specific, although we cannot rule out that the genes were simply not recovered in the genome assembly (BUSCO completeness 92.2%). More interestingly, the AMS-specific genes were mostly retained in the ECM species that lost AMS: *C. fangiana* and *B. pendula* retained all three genes, whereas *Q. robur* and *F. sylvatica* retained two of the tree genes (Fig. 2). Similarly, among the CSP genes, *SYMRK* and *CCaMK* were found in all ECM species, including those that lost AMS (Fig. 2). The same pattern was observed for CYCLOPS with the notable exception of Fagaceae and the genus *Morella*, in which the gene was either not detected or only partially recovered. Although this may be attributable to low quality genomes, the observation that these potential losses are restricted to a single clade might deserve further investigation. Altogether, this suggests there may be co-option of these genes from AMS to independent instances of ECM, or that the losses of AMS in ECM-only lineages are relatively recent.

The combination of ancestral state reconstruction, comparative phylogenomics and transcriptomics, and single-gene phylogeny conducted here do not support an ancestral gain of ECM in rosids, but rather the independent evolution of this symbiotic ability. These independent events, involving different fungal lineages (Kohler *et al*., 2015), occurred in a convergent manner by gene co-option and duplications leading to the emergence of a similar transcriptional response.

## Acknowledgements

The authors thank the Genotoul bioinformatics platform Toulouse Occitanie (Bioinfo Genotoul, https://doi.org/10.15454/1.5572369328961167E12) for providing computing resources. P-M.D was supported by the project Engineering Nitrogen Symbiosis for Africa (ENSA) currently funded through a grant to the University of Cambridge by the Bill and Melinda Gates Foundation (OPP1172165) and the UK Foreign, Commonwealth and Development Office as Engineering Nitrogen Symbiosis for Africa (OPP1172165). This project received funding from the European Research Council (ERC) under the European Union’s Horizon 2020 research and innovation programme (grant agreement no. 101001675 - ORIGINS) to P.-M.D. This project was supported by the Laboratoires d’Excellence (LABEX)’ TULIP (ANR-10-LABX-41) and the ‘École Universitaire de Recherche (EUR)’ TULIP-GS (ANR-18-EURE-0019). MBi has received funding from the European Union’s Horizon 2020 research and innovation programme under the Marie Skłodowska-Curie grant agreement No. 101105838 as SYMBIOLOSS.

## Data availability

Code used for this work is available on gitlab [link] and phylogenies and alignments can be found on figshare [link]. Genome data used is listed in table S1 and sources for the RbcL and MatK sequences in table S3.

