## Supplementary information for "Ectomycorrhizal symbiosis evolved independently and by convergent gene duplication in rosid lineages"

The following Supporting Information is available for this article:

**Table S1 (see Excel file).** Genome statistics and sources of all included species.

**Table S2 (see Excel file).** Number of duplications for 10 different taxonomic groups are shown for the nine genes frequently duplicated in ECM lineages.

**Table S3 (see Excel file).** Sources of the MatK and RbcL protein sequences

**Table S4 (see Excel file).** Results for the tested models for ancestral state reconstruction.

**Methods S1.**

**Fig. S1-6**: Single gene phylogeny of cytochrome P450s (S1), SDR-like proteins (S2), Zinc fingers (S3) and HAD-like proteins (S4) that are largely retained in ECM lineages.

**Fig. S7**: Random sampling approach testing the significance of the number of orthogroups with genes deregulated in three ECM plant species.

**Fig. S8-16**: Single gene phylogeny of MRP-like ABC transporters (S6), aspartyl proteases (S7), NDR-like proteins (S8), isoprenoid synthases (S9), pectinesterase inhibitor 24s (S10), GH5 cellulases (S11), MoO transporters (S12), ring-type zinc fingers (S13), pectin lyases (S14), O-methyltransferases (S15) and Myb-type transcription factors (S16) that are convergently duplicated in ECM lineages.

**Fig. S17-19**: Single gene phylogenies of the CSP genes SYMRK (S15), CCaMK (S16) and CYCLOPS (S17).

**Fig. S20-22**: Single gene phylogenies of AMS-related genes RAD1 (S18), STR1 (S19) and STR2 (S20).

**Fig. S23**: A rooted cladogram showing the constrained tree used for the reconstruction of rosids and other angiosperms based on the plastid protein sequences of RbcL and MatK.

**Fig. S24**: A rooted cladogram showing the constrained tree used for the reconstruction of rosids and other angiosperms based on nuclear data with OrthoFinder.

**Fig. S25**: Venn diagrams showing the overlap in differentially expressed orthogroups (DEOGs) between the ECM plant species *Quercus robur*, *Populus tremula* × *P. tremuloides* and *Eucalyptus grandis.*

**Fig. S26**: Heatmap of hierarchical orthologous groups (HOGs) of the node representing the common ancestor of ECM-rosids.

**Methods S1**

### Ancestral state reconstruction

Protein sequences of RbcL and MatK were acquired for 133 angiosperms from NCBI and UniProt (Supplementary table S3) (Benson *et al.*, 2013; The UniProt Consortium, 2015), except for the species *Cryptosepalum pellegrinianum* and *Paramacrolobium coeruleum* as only the MatK sequences are known. For the genera *Dicymbe* and *Poranthera* RbcL and MatK sequences were derived from two different species. For each gene, protein sequences were aligned using MAFFT L-INS-i (v7.526, default parameters) and the alignment was trimmed using trimAl (v1.4.rev15) with parameter -gt 0.1 (Katoh *et al.*, 2005; Capella-Gutiérrez *et al.*, 2009; Katoh & Standley, 2013). Subsequently, the alignments were concatenated and the phylogeny was reconstructed with IQ-TREE (v2.2.2.3) with parameters -m TESTNEW -mset Q.plant,cpREV,LG -mfreq FU,F,FO -mrate G,I+G,R (Minh *et al.*, 2020). As two plastid genes cannot be expected to resolve deep relationships in the tree of angiosperms, the tree was constrained based on known relationships between the different angiosperms (Fig. S23), following APweb (Stevens & Davis, 2005; Stevens, 2017). The final phylogeny was reconstructed using the best fitting model Q.plant+C60+FO+R5 and the parameters --ufboot 10000 --alrt 10000 (Guindon *et al.*, 2010; Hoang *et al.*, 2018).

To reconstruct ancestral states, we used the RbcL+MatK phylogeny and the known states of different species largely following Tedersoo & Brundrett (2017), with the exception of the state of *Juglans regia* (Li *et al.*, 2024). Multiple models were tested using the *geiger* package v2.0.11 in R v4.4.1 (Pennell *et al.*, 2014; R Core Team, 2024), comparing all possible combinations of models and transforms (Supplementary table S4; code available on gitlab). The best model was ARD (All Rates Different) + kappa (Pagel, 1999), based on the corrected Akaike Information Criterion. To visualize the results, we fit the above model with the same value for kappa with the *ace* package in the R package *ape* v5.8 (Paradis & Schliep, 2019). Model testing results were merged using the R package *tidyverse* v2.0.0 and the phylogeny was plotted using the R package *ggtree* v3.11.2 (Yu *et al.*, 2017; Wickham *et al.*, 2019).

### Gene annotation and grouping of orthologous genes

A total of 44 rosids and 19 outgroup angiosperm genomes were included in the analysis from different sources (Supplementary table S1). We annotated six genomes for which no annotation was available (*Acacia melanoxylon*, *Corylus avellana*, *C. heterophylla*, *Eucalyptus pauciflora*, *Populus simonii* and *Quercus mongolica*). Repeats were soft-masked using Red (Girgis, 2015). Structural annotation was performed using BRAKER2 (version e56d039) in EP-mode on the soft-masked genomes (Stanke *et al.*, 2006, 2008; Hoff *et al.*, 2016, 2019; Brůna *et al.*, 2021). As input for the ProtHint pipeline in BRAKER2, the Viridiplantae OrthoDB (version 10) and the locally created angiosperm OrthoDB (Kriventseva *et al.*, 2019) were used. Genes with support at any of the exons were kept for further analysis. From the proteomes, the longest isoform of all predicted genes was retained, with a minimal length of 20 amino acids. To remove genes overlapping repeated regions, repeats were hard-masked using RepeatModeler (version 2.0.1) and RepeatMasker (version 4.1.2) (Chen, 2004; Flynn *et al.*, 2020) and proteins predicted in repeat-masked regions were filtered out from the proteomes. Genome completion was estimated using BUSCO v5.7.1 with the Viridiplantae OrthoDB with the miniprot gene predictor (Manni *et al.*, 2021; Li, 2023).

To reconstruct orthogroups, OrthoFinder v2.5.5 (Emms & Kelly, 2019) was used for a first round using diamond_ultra_sens as the sequence search program. The species tree was adjusted following known relationships among angiosperms (Fig. S24) based on APweb (Stevens & Davis, 2005; Stevens, 2017); the structure is comparable to that of the RbcL+MatK phylogeny. A second orthofinder analysis was done based on this tree (using -M msa).

**Comparative differential expression**

For *Populus tremula* × *P. tremuloides* RNAseq reads were collected from BioProject PRJEB41173 (Chowdhury *et al.*, 2022), excluding the free-living mycelia control (FLM). For *Eucalyptus grandis*, reads were collected from the SRA (SRR6962691-SRR6962693, SRR8246608-SRR8246625), also excluding reads related to free-living mycelium (Hill *et al.*, 2022). For *Quercus robur*, reads were collected from BioProject PRJNA516042 representing experiments with two different ECM-fungi (*Paxillus involutus* and *Pisolithus microcarpus*) and the control (Bouffaud *et al.*, 2020). For *Q. robur*, data from the experiment with the ECM fungus *Laccaria bicolor* was not included as data for only one replicate was available.

For *E. grandis* and *Q. robur,* reads were mapped to its relevant genome annotation included in the OrthoFinder analysis. For *P. tremula* × *P. tremuloides* for which the reads were mapped to the genome of *P. tremula* (Schiffthaler *et al.*, 2019). For all three species, RNAseq analysis was done using Nextflow (v21.10.6) using the RNAseq pipeline from nf-core (3.9) with the parameters -profile genotoul --aligner star_salmon --remove_ribo_rna --skip_qc (refs), which includes Trimgalore, picard, STAR, Salmon, BEDTools, MultiQC, GffRead, StringTie2 and UCSC tools (Quinlan & Hall, 2010; Kent *et al.*, 2010; Dobin *et al.*, 2013; Krueger, 2015; Ewels *et al.*, 2016, 2022; Di Tommaso *et al.*, 2017; Patro *et al.*, 2017; Kovaka *et al.*, 2019; Pertea & Pertea, 2020).

Calling differentially expressed genes was done using the R package *edgeR* v3.42.4 (Robinson *et al.*, 2010; Chen *et al.*, 2014). For each analysis, genes with 10 or fewer reads mapped to it were excluded. Then, for each comparison between treatment and control, the gene counts were normalized by library size and trimmed mean of M-values (Robinson & Oshlack, 2010). Differential expression was estimated using the glmLRT function in edgeR, using a maximum FDR of 0.05 and a minimum LogFC of 0 (*Q. robur* and *Populus tremula* × *P. tremuloides*) or 1.2 (*E. grandis*) to retain comparable sets of genes. Up- and downregulated genes were kept for every condition and every time point.

To be able to compare the RNAseq between the three different species, deregulation was summarized for each species per orthogroup. To visualize overlaps between up- and down-regulated genes (Fig. S25), the R package VennDiagram v1.7.3 was used (Chen & Boutros, 2011). To estimate whether found overlaps between the three species could be found by chance, we made use of a random-sampling approach. The same number of genes based on the experimental data were randomly sampled 100,000 times for each species using the *srswor* (simple random sampling without replacement) function from the R package *sampling* v2.10 (Tillé, 2006) and the *powerSet* function from the R package *rje* v1.12.1. Overlaps between orthogroups were estimated and a distribution was made based on these samples. To calculate the statistical difference between the observed overlap and estimated overlaps, we estimated the p-value as the proportion of random samples that showed a higher number of overlap than the observed number of overlaps. Code used for the estimation of overlaps can be found on gitlab.

**Detection of ancestrally retained and convergently duplicated genes**

To detect whether any genes were potentially present in the common ancestor of ECM-rosids, but lost in non-ECM lineages, a heatmap was made of all hierarchical orthogroups (HOGs) based ECM and non-ECM lineages encoding genes part of that HOG, and 264 HOGs enriched in ECM-lineages were selected for further analysis (Fig. S26). To select those likely related to the symbiosis, we kept only those deregulated in at least two species (41 HOGs). For these HOGs, seed sequences were taken of five species. representing the five ECM lineages included this analysis (*E. grandis*, *P. tremula*, *Q. robur*, *Shorea leprosula*, and *Acacia melanoxylon*) and aligned against all the protein sequences of the 63 species included in this analysis using BLASTP v2.15.0 (Altschul *et al.*, 1990). Preliminary alignments were done with MAFFT (default parameters) and trimmed with trimAl (-gt 0.3) and phylogenetic reconstruction was done using IQ-TREE with parameters -fast -m LG+G4. Trees were inspected for the presence of potential clades largely only retained by ECM and the clade including an outgroup was aligned again using MAFFT and trimAl (same parameters) followed by phylogenetic reconstruction with IQ-TREE using parameters -m TESTNEW -mset Q.plant,LG,WAG,JTT --ufboot 10000 --alrt 10000. To assure that the pattern found was not limited by our taxon selection, seed sequences were aligned to the RefSeq database using BLASTP (O'Leary *et al.*, 2016). The retained six sequence sets were then aligned, trimmed and reconstructed as above and trees were visualized using iTOL (Letunic & Bork, 2024).

To detect genes duplicated in multiple ECM lineages, we made use of the Gene_Duplication_Events results from OrthoFinder. To only keep highly-confident results, we filtered the results to a minimum support of 50%. For the five ECM clades represented in our analysis, we looked for common duplication events. As Fagales and Salicaceae were represented each by more than one genus, we specifically focused on duplications occurring at least in the common ancestor of each of these two groups, leaving 61 orthogroups. These were filtered further in three ways. All orthogroups were kept where at least one duplicated gene was deregulated in all three species with RNAseq data. Also, orthogroups where many (1 out of 7) of the duplications occurred in nodes where ECM was estimated to originate were kept, as these would be the nodes where duplication occurs related to convergent evolution. Finally, orthogroups where 50% or more of duplications occurred in any ECM-node were kept. This resulted in a set of 27 candidates. Phylogenies were made as above, but instead looking for convergent duplications in the most ancestral nodes of the five represented ECM lineages. After verification 9 candidates were kept as they showed clear signs of convergent duplications and were visualized using iTOL (Letunic & Bork, 2024). Manual inspection of final single gene trees was done to estimate the number of duplications for the 11 different nodes shown in Fig. 2, and these results were visualized in R using ggplot2 v3.5.1 (Wickham, 2016).

**Detection of symbiosis-related genes**

We searched the genomes for sequences of six protein-coding genes known to be associated with the establishment of symbiotic associations in plants. These included genes from the common symbiotic signaling pathway (CSSP; CCaMK/DIM3, CYCLOPS/IPD3 and SYMRK/DMI2), and genes specific to AM symbiosis (RAD1, STR and STR2), which were shown to have been consistently lost in non-symbiotic and non-AM lineages, respectively (Radhakhrishnan et al., 2020). BLASTP searches (e-value = $10^{-9}$) were conducted on proteomes using as queries sequences of *Medicago truncatula*. Phylogenetic trees were then used to identify the orthologous groups containing the target genes. First, protein sequences were aligned using MAFFT in the automatic mode (*--auto*, *--maxiterate 10*), and alignments were trimmed using trimAl (*-gt 0.3*). An approximately-maximum-likelihood phylogenetic tree was then inferred using FastTree v2.1.11 (Price et al., 2010) with default parameters. We conducted an additional search for the target genes on the genome sequences to rule out the possibility of gene absence due to potential genome annotation issues. For that, we used Exonerate v2.2.0 (Slater & Birner, 2005) with protein queries (--model protein2genome), and reporting only the five best matches (--bestn 5). Sequences were then aligned with the nucleotide sequences corresponding to the orthologs retrieved using BLASTP, trimmed, and a phylogenetic tree was inferred on the resulting alignment as described above. After a visual inspection of the resulting tree, orthologous sequences were extracted, realigned with MAFFT L-INS-i and trimmed as above, and a final phylogenetic tree was inferred using IQ-TREE with ultrafast bootstrap and SH-like approximate likelihood ratio tests (-m TESTNEW --ufboot 10000 --alrt 10000).

**Fig. S1**: Single gene phylogeny of cytochrome P450s that are largely retained in ECM lineages. Gene names from our database (Supplementary table S1) are shown with full species name and colored by the taxonomic placement of that species. ECM status is shown for all sequences from the database, and for all sequences in the putative clade of genes retained in ECM lineages. The tree was reconstructed with the model Q.plant+R7. Support is shown based on ultrafast bootstrap. The scale represents the estimated number of substitutions per site.

**Fig. S2**: Single gene phylogeny of SDR-like proteins that are largely retained in ECM lineages. Gene names from our database (Supplementary table S1) are shown with full species name and colored by the taxonomic placement of that species. ECM status is shown for all sequences from the database, and for all sequences in the putative clade of genes retained in ECM lineages. The tree was reconstructed with the model JTT+R6. Support is shown based on ultrafast bootstrap. The scale represents the estimated number of substitutions per site.

**Fig. S3**: Single gene phylogeny of Zinc fingers that are largely retained in ECM lineages. Gene names from our database (Supplementary table S1) are shown with full species name and colored by the taxonomic placement of that species. ECM status is shown for all sequences from the database, and for all sequences in the putative clade of genes retained in ECM lineages. The tree was reconstructed with the model JTT+I+G4. Support is shown based on ultrafast bootstrap. The scale represents the estimated number of substitutions per site.

**Fig. S4**: Single gene phylogeny of HAD-like proteins that are largely retained in ECM lineages. Gene names from our database (Supplementary table S1) are shown with full species name and colored by the taxonomic placement of that species. ECM status is shown for all sequences from the database, and for all sequences in the putative clade of genes retained in ECM lineages. The tree was reconstructed with the model WAG+R6. Support is shown based on ultrafast bootstrap. The scale represents the estimated number of substitutions per site.

**Fig. S5**: Single gene phylogeny of O-methyltransferases that are largely retained in ECM lineages. Gene names from our database (Supplementary table S1) are shown with full species name and colored by the taxonomic placement of that species. ECM status is shown for all sequences from the database, and for all sequences in the putative clade of genes retained in ECM lineages. The tree was reconstructed with the model Q.plant+R5. Support is shown based on ultrafast bootstrap. The scale represents the estimated number of substitutions per site.

**Fig. S6**: Single gene phylogeny of Myb-type transcription factors that are largely retained in ECM lineages. Gene names from our database (Supplementary table S1) are shown with full species name and colored by the taxonomic placement of that species. ECM status is shown for all sequences from the database, and for all sequences in the putative clade of genes retained in ECM lineages. The tree was reconstructed with the model Q.plant+I+R7. Support is shown based on ultrafast bootstrap. The scale represents the estimated number of substitutions per site.

**Fig. S7**: Random sampling approach testing the significance of the number of orthogroups with genes deregulated in three ECM plant species. Red line shows the observed number of orthogroups deregulated during ECM symbiosis in three different plant species: *Quercus robur*, *Populus tremula* × *P. tremuloides* and *Eucalyptus grandis*. The results of randomly sampling the same number of genes for each species and estimating the overlaps are shown in blue, based on 100,000 random samples. The mean is shown as a dashed blue line. Figure is split in showing orthogroups with genes deregulated in all species in any direction (top), genes upregulated in all three species (middle) and genes downregulated in all three species (bottom).

**Fig. S8**: Single gene phylogeny of MRP-like ABC transporters. Nodes representing a duplication in one of the lineages shown in Fig. 2 (main text) are highlighted with a star. The tree was reconstructed with the model Q.plant+F+R8. Support is shown based on ultrafast bootstrap. The scale represents the estimated number of substitutions per site.

**Fig. S9**: Single gene phylogeny of aspartyl proteases. Nodes representing a duplication in one of the lineages shown in Fig. 2 (main text) are highlighted with a star. The tree was reconstructed with the model JTT+R8. Support is shown based on ultrafast bootstrap. The scale represents the estimated number of substitutions per site.

**Fig. S10**: Single gene phylogeny of NDR-like proteins. Nodes representing a duplication in one of the lineages shown in Fig. 2 (main text) are highlighted with a star. The tree was reconstructed with the model Q.plant+R7. Support is shown based on ultrafast bootstrap. The scale represents the estimated number of substitutions per site.

**Fig. S11**: Single gene phylogeny of isoprenoid synthases. Nodes representing a duplication in one of the lineages shown in Fig. 2 (main text) are highlighted with a star. The tree was reconstructed with the model Q.plant+R7. Support is shown based on ultrafast bootstrap. The scale represents the estimated number of substitutions per site.

**Fig. S12**: Single gene phylogeny of pectinesterase inhibitor 24s. Nodes representing a duplication in one of the lineages shown in Fig. 2 (main text) are highlighted with a star. The tree was reconstructed with the model Q.plant+R8. Support is shown based on ultrafast bootstrap. The scale represents the estimated number of substitutions per site.

**Fig. S13**: Single gene phylogeny of GH5 cellulases. Nodes representing a duplication in one of the lineages shown in Fig. 2 (main text) are highlighted with a star. The tree was reconstructed with the model Q.plant+I+R7. Support is shown based on ultrafast bootstrap. The scale represents the estimated number of substitutions per site.

**Fig. S14**: Single gene phylogeny of MoO transporters. Nodes representing a duplication in one of the lineages shown in Fig. 2 (main text) are highlighted with a star. The tree was reconstructed with the model Q.plant+I+R7. Support is shown based on ultrafast bootstrap. The scale represents the estimated number of substitutions per site.

**Fig. S15**: Single gene phylogeny of ring-type zinc fingers. Nodes representing a duplication in one of the lineages shown in Fig. 2 (main text) are highlighted with a star. The tree was reconstructed with the model Q.plant+I+R5. Support is shown based on ultrafast bootstrap. The scale represents the estimated number of substitutions per site.

**Fig. S16**: Single gene phylogeny of pectin lyases. Nodes representing a duplication in one of the lineages shown in Fig. 2 (main text) are highlighted with a star. The tree was reconstructed with the model JTT+R5. Support is shown based on ultrafast bootstrap. The scale represents the estimated number of substitutions per site.

**Fig. S17**: Single gene phylogeny of SYMRK. The tree was reconstructed with the model GTR+F+R4. Support is shown based on ultrafast bootstrap. The scale represents the estimated number of substitutions per site.

**Fig. S18**: Single gene phylogeny of CCaMK. The tree was reconstructed with the model GTR+F+R4. Support is shown based on ultrafast bootstrap. The scale represents the estimated number of substitutions per site.

**Fig. S19**: Single gene phylogeny of CYCLOPS. The tree was reconstructed with the model GTR+F+R4. Support is shown based on ultrafast bootstrap. The scale represents the estimated number of substitutions per site.

**Fig. S20**: Single gene phylogeny of RAD1. The tree was reconstructed with the model TPM2+R4. Support is shown based on ultrafast bootstrap. The scale represents the estimated number of substitutions per site.

**Fig. S21**: Single gene phylogeny of STR1. The tree was reconstructed with the model TVMe+R4. Support is shown based on ultrafast bootstrap. The scale represents the estimated number of substitutions per site.

**Fig. S22**: Single gene phylogeny of STR2. The tree was reconstructed with the model SYM+R4. Support is shown based on ultrafast bootstrap. The scale represents the estimated number of substitutions per site.

**Fig. S23**: A rooted cladogram showing the constrained tree used for the reconstruction of rosids and other angiosperms based on the plastid protein sequences of RbcL and MatK. Relationships with no clear consensus were not constrained, and are thus shown as polytomies.

**Fig. S24**: A rooted cladogram showing the constrained tree used for the reconstruction of rosids and other angiosperms based on nuclear data with OrthoFinder.

**Fig. S25**: Venn diagrams showing the overlap in differentially expressed orthogroups (DEOGs) between the ECM plant species *Quercus robur*, *Populus tremula* × *P. tremuloides* and *Eucalyptus grandis.* Results are orthogroups deregulated in any direction in the tree species (top), only up-regulated genes (middle) and only down-regulated genes (bottom).

**Fig. S26**: Heatmap of hierarchical orthologous groups (HOGs) of the node representing the common ancestor of ECM-rosids. The axes represent the number of ECM species (x-axis) and non-ECM species (y-axis) with a gene in each HOG. Squares with thick lines contain the HOGs further analyzed for gene retention using single gene phylogenies.

Fig. S01

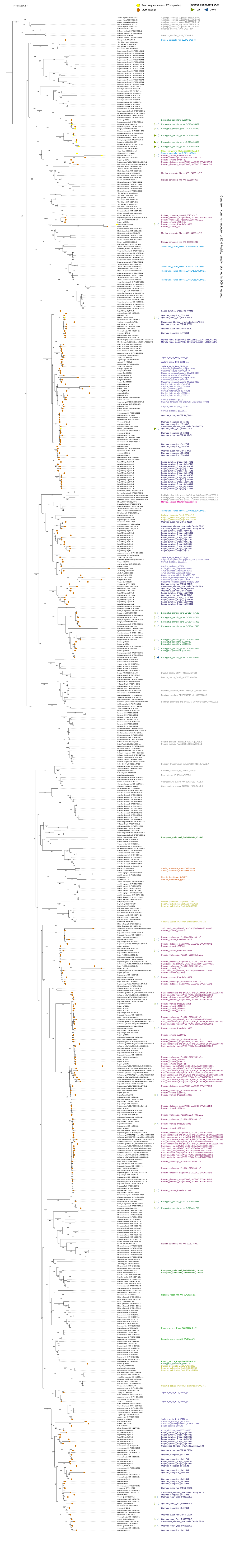
Tree scale: 0.1
Seed sequences (and ECM species)
ECM species
Expression during ECM
Up
Down
Gene from common ancestor of ECM Rosids, largely retained in ECM lineages.

Fig. S02

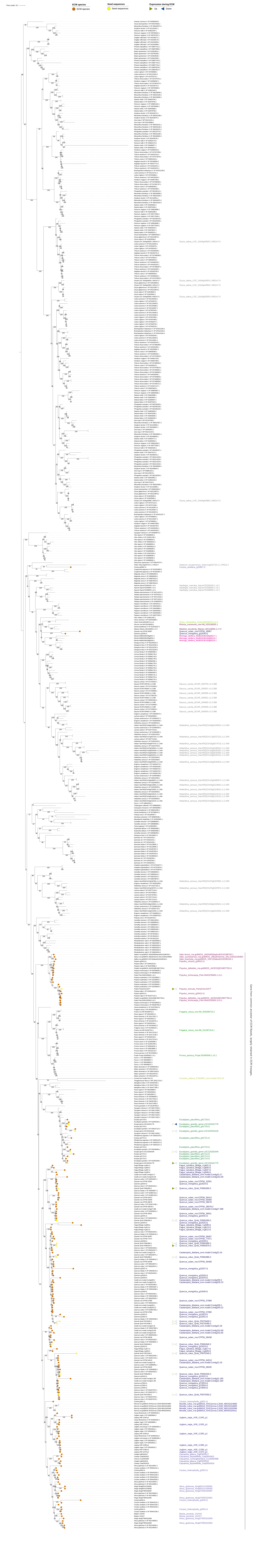

Tree scale: 0.1
ECM species
ECM species
Seed sequences
Seed sequences
Expression during ECM
Up
Down

Fig. S03

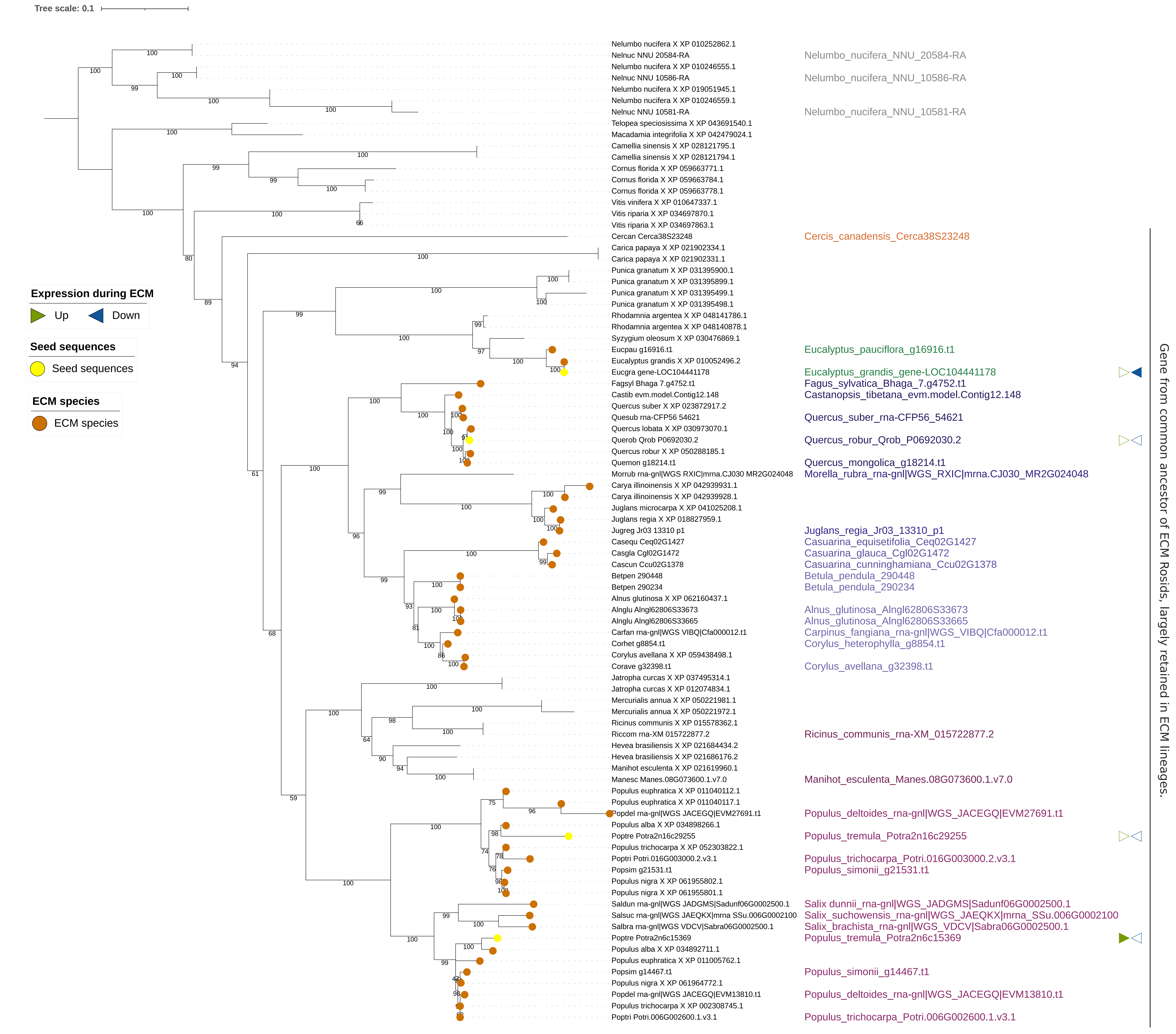
Tree scale: 0.1
Expression during ECM
Up
Down
Seed sequences
Seed sequences
ECM species
ECM species
Nelumbo nucifera X XP 010252862.1
Nelnuc NNU 20584-RA
Nelumbo nucifera X XP 010246555.1
Nelnuc NNU 10586-RA
Nelumbo nucifera X XP 019051945.1
Nelumbo nucifera X XP 010246559.1
Nelnuc NNU 10581-RA
Telopea speciosissima X XP 043691540.1
Macadamia integrifolia X XP 042479024.1
Camellia sinensis X XP 028121795.1
Camellia sinensis X XP 028121794.1
Cornus florida X XP 059663771.1
Cornus florida X XP 059663784.1
Cornus florida X XP 059663778.1
Vitis vinifera X XP 010647337.1
Vitis riparia X XP 034697870.1
Vitis riparia X XP 034697863.1
Cercan Cerca38S23248
Carica papaya X XP 021902334.1
Carica papaya X XP 021902331.1
Punica granatum X XP 031395900.1
Punica granatum X XP 031395899.1
Punica granatum X XP 031395499.1
Punica granatum X XP 031395498.1
Rhodamnia argentea X XP 048141786.1
Rhodamnia argentea X XP 048140878.1
Syzygium oleosum X XP 030476869.1
Eucpau g16916.t1
Eucalyptus grandis X XP 010052496.2
Eucgra gene-LOC104441178
Fagsyl Bhaga 7.g4752.t1
Castib evm.model.Contig12.148
Quercus suber X XP 023872917.2
Quesub rna-CFP56 54621
Quercus lobata X XP 030973070.1
Querob Qrob P0692030.2
Quercus robur X XP 050288185.1
Quemon g18214.t1
Morrub rna-gnl|WGS RXIC|mrna.CJ030 MR2G024048
Carya illinoinensis X XP 042939931.1
Carya illinoinensis X XP 042939928.1
Juglans microcarpa X XP 041025208.1
Juglans regia X XP 018827959.1
Jugreg Jr03 13310 p1
Casequ Ceq02G1427
Casgla Cgl02G1472
Cascun Ccu02G1378
Betpen 290448
Betpen 290234
Alnus glutinosa X XP 062160437.1
Alnglu Alngl62806S33673
Alnglu Alngl62806S33665
Carfan rna-gnl|WGS VIBQ|Cfa000012.t1
Corhet g8854.t1
Corylus avellana X XP 059438498.1
Corave g32398.t1
Jatropha curcas X XP 037495314.1
Jatropha curcas X XP 012074834.1
Mercurialis annua X XP 050221981.1
Mercurialis annua X XP 050221972.1
Ricinus communis X XP 015578362.1
Riccom rna-XM 015722877.2
Hevea brasiliensis X XP 021684434.2
Hevea brasiliensis X XP 021686176.2
Manihot esculenta X XP 021619960.1
Manesc Manes.08G073600.1.v7.0
Populus euphratica X XP 011040112.1
Populus euphratica X XP 011040117.1
Popdel rna-gnl|WGS JACEGQ|EVM27691.t1
Populus alba X XP 034898266.1
Poptre Potra2n16c29255
Populus trichocarpa X XP 052303822.1
Poptri Potri.016G003000.2.v3.1
Popsim g21531.t1
Populus nigra X XP 061955802.1
Populus nigra X XP 061955801.1
Saldun rna-gnl|WGS JADGMS|Sadunf06G0002500.1
Salsuc rna-gnl|WGS JAEQKX|mrna SSu.006G0002100
Salbra rna-gnl|WGS VDCV|Sabra06G0002500.1
Poptre Potra2n6c15369
Populus alba X XP 034892711.1
Populus euphratica X XP 011005762.1
Popsim g14467.t1
Populus nigra X XP 061964772.1
Popdel rna-gnl|WGS JACEGQ|EVM13810.t1
Populus trichocarpa X XP 002308745.1
Poptri Potri.006G002600.1.v3.1
Nelumbo_nucifera_NNU_20584-RA
Nelumbo_nucifera_NNU_10586-RA
Nelumbo_nucifera_NNU_10581-RA
Cercis_canadensis_Cerca38S23248
Eucalyptus_pauciflora_g16916.t1
Eucalyptus_grandis_gene-LOC104441178
Fagus_sylvatica_Bhaga_7.g4752.t1
Castanopsis_tibetana_evm.model.Contig12.148
Quercus_suber_rna-CFP56_54621
Quercus_robur_Qrob_P0692030.2
Quercus_mongolica_g18214.t1
Morella_rubra_rna-gnl|WGS_RXIC|mrna.CJ030_MR2G024048
Juglans_regia_Jr03_13310_p1
Casuarina_equisetifolia_Ceq02G1427
Casuarina_glauca_Cgl02G1472
Casuarina_cunninghamiana_Ccu02G1378
Betula_pendula_290448
Betula_pendula_290234
Alnus_glutinosa_Alngl62806S33673
Alnus_glutinosa_Alngl62806S33665
Carpinus_fangiana_rna-gnl|WGS_VIBQ|Cfa000012.t1
Corylus_heterophylla_g8854.t1
Corylus_avellana_g32398.t1
Ricinus_communis_rna-XM_015722877.2
Manihot_esculenta_Manes.08G073600.1.v7.0
Populus_deltoides_rna-gnl|WGS_JACEGQ|EVM27691.t1
Populus_tremula_Potra2n16c29255
Populus_trichocarpa_Potri.016G003000.2.v3.1
Populus_simonii_g21531.t1
Salix dunnii_rna-gnl|WGS_JADGMS|Sadunf06G0002500.1
Salix_suchowensis_rna-gnl|WGS_JAEQKX|mrna_SSu.006G0002100
Salix_brachista_rna-gnl|WGS_VDCV|Sabra06G0002500.1
Populus_tremula_Potra2n6c15369
Populus_simonii_g14467.t1
Populus_deltoides_rna-gnl|WGS_JACEGQ|EVM13810.t1
Populus_trichocarpa_Potri.006G002600.1.v3.1
Gene from common ancestor of ECM Rosids, largely retained in ECM lineages.

Fig. S04

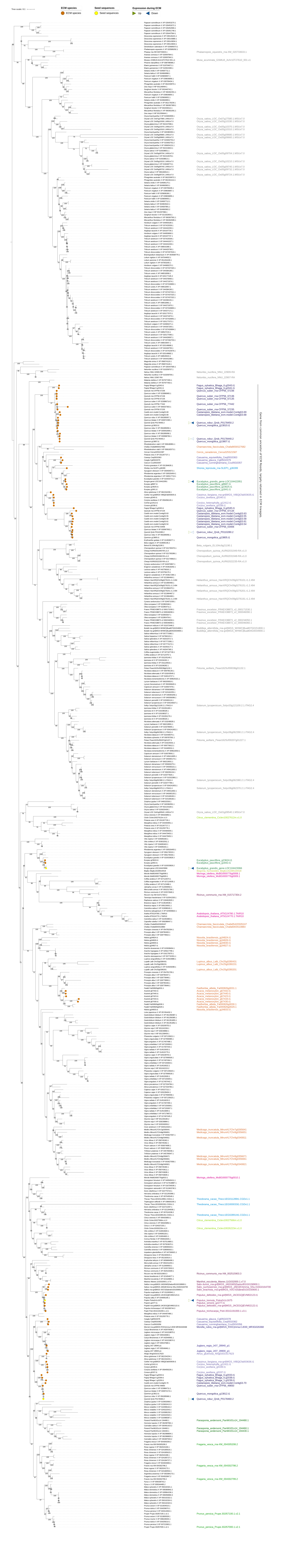

Tree scale: 0.1
ECM species
ECM species
Seed sequences
Seed sequences
Expression during ECM
Up
Down
Papaver somniferum X XP 026453270.1
Papaver somniferum X XP 026453272.1
Papaver somniferum X XP 026452906.1
Papaver somniferum X XP 026451748.1
Papaver somniferum X XP 026447564.1
Dioscorea cayenensis X XP 039126221.1
Dioscorea cayenensis X XP 039143880.1
Dioscorea cayenensis X XP 039143834.1
Dioscorea cayenensis X XP 039143833.1
Dendrobium catenatum X XP 020685073.1
Phalaenopsis equestris X XP 020589292.1
Phaequ rna-XM 020733633.1
Ananas comosus X XP 020097844.1
Ananas comosus X XP 020097842.1
Musacu GSMUA Achr10T27610 001.v1
Phoenix dactylifera X XP 008798588.2
Elaeis guineensis X XP 019704671.1
Elaeis guineensis X XP 010915365.1
Setaria viridis X XP 034587711.1
Setaria italica X XP 004960898.1
Panicum hallii X XP 025806957.1
Panicum virgatum X XP 039836856.1
Panicum virgatum X XP 039799434.1
Phragmites australis X XP 062230970.1
Zea mays X NP 001148494.1
Sorghum bicolor X XP 002440742.1
Miscanthus floridulus X XP 066362351.1
Panicum virgatum X XP 039836854.1
Panicum hallii X XP 025806952.1
Setaria viridis X XP 004960899.1
Phragmites australis X XP 062179239.1
Gene from common ancestor of ECM Rosids, largely retained in ECM lineages.

Fig. S05

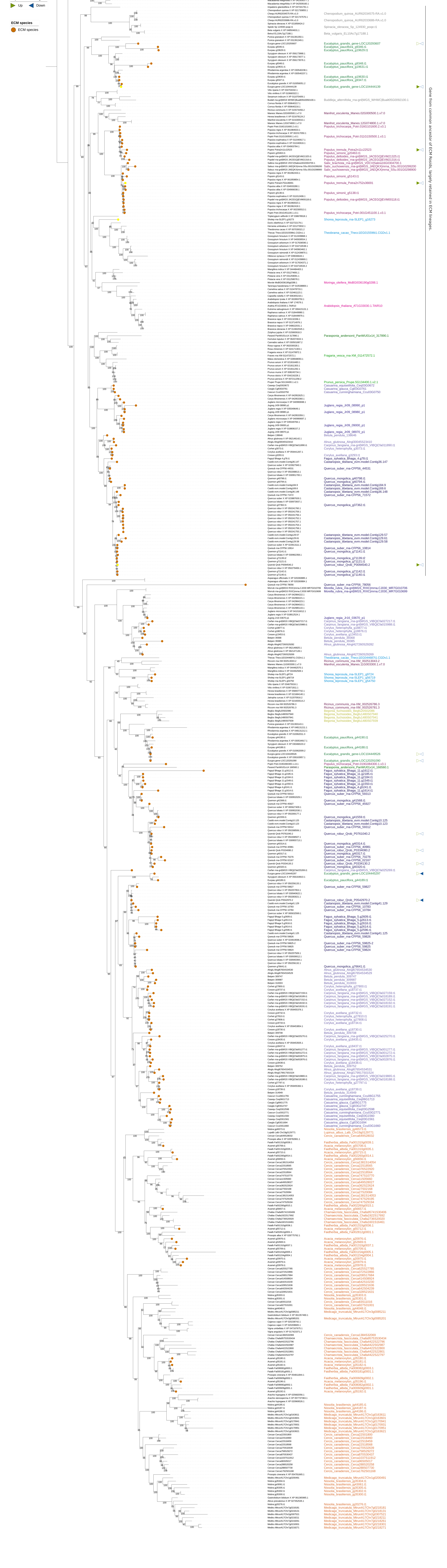
Tree scale: 0.1
Seed sequences
Seed sequences
Expression during ECM
Up
Down
ECM species
ECM species
Gene from common ancestor of ECM Rosids, largely retained in ECM lineages.

Fig. S06

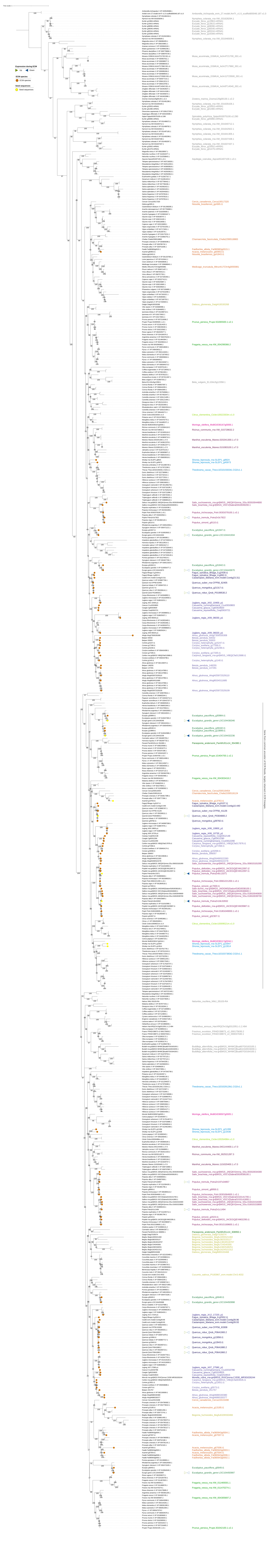
Tree scale: 1
Expression during ECM
Up
Down
ECM species
ECM species
Seed sequences
Seed sequences

Fig. S07

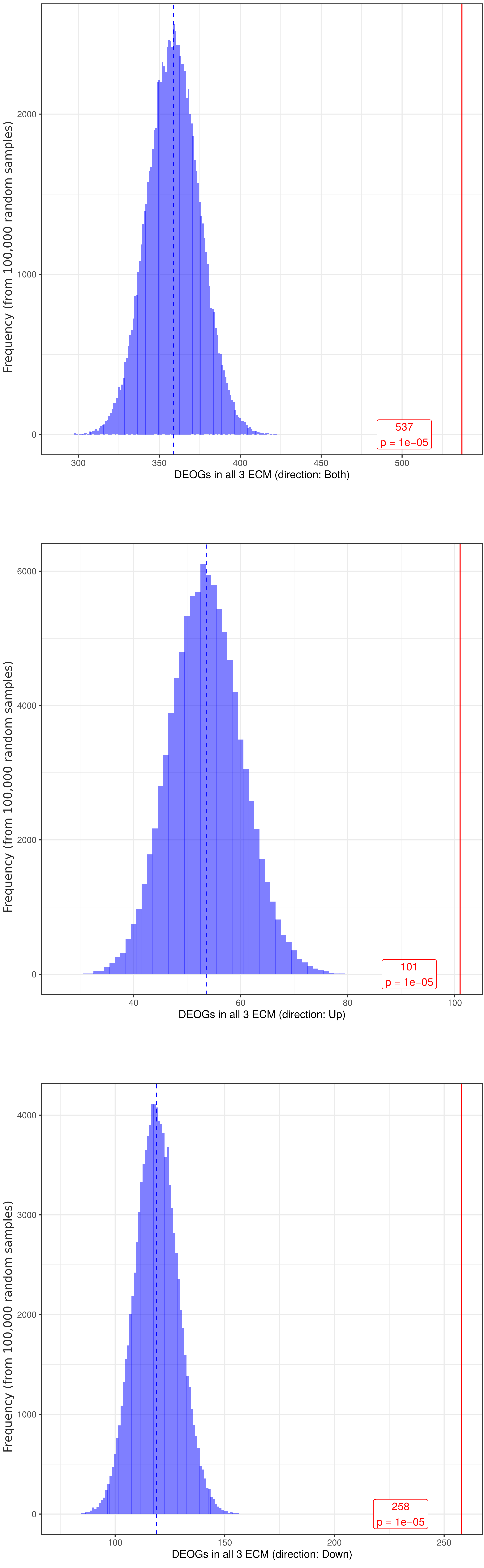

Frequency (from 100,000 random samples)
2000
1000
0
537
p = 1e−05
300
350
400
450
500
DEOGs in all 3 ECM (direction: Both)
Frequency (from 100,000 random samples)
6000
4000
2000
0
101
p = 1e−05
40
60
80
100
DEOGs in all 3 ECM (direction: Up)
Frequency (from 100,000 random samples)
4000
3000
2000
1000
0
258
p = 1e−05
100
150
200
250
DEOGs in all 3 ECM (direction: Down)

Fig. S08

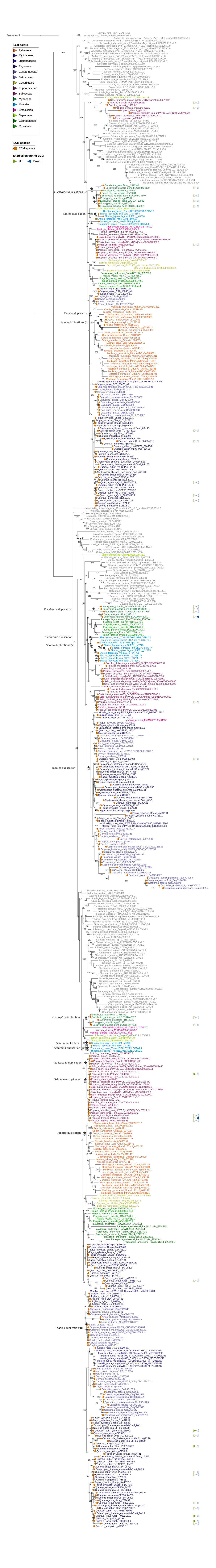

Tree scale: 1
Leaf colors
Fabaceae
Myricaceae
Juglandaceae
Fagaceae
Casuarinaceae
Betulaceae
Cucurbitales
Euphorbiaceae
Salicaceae
Myrtaceae
Malvales
Brassicales
Sapindales
Cannabaceae
Rosaceae
ECM species
ECM species
Expression during ECM
Up
Down

Fig. S09

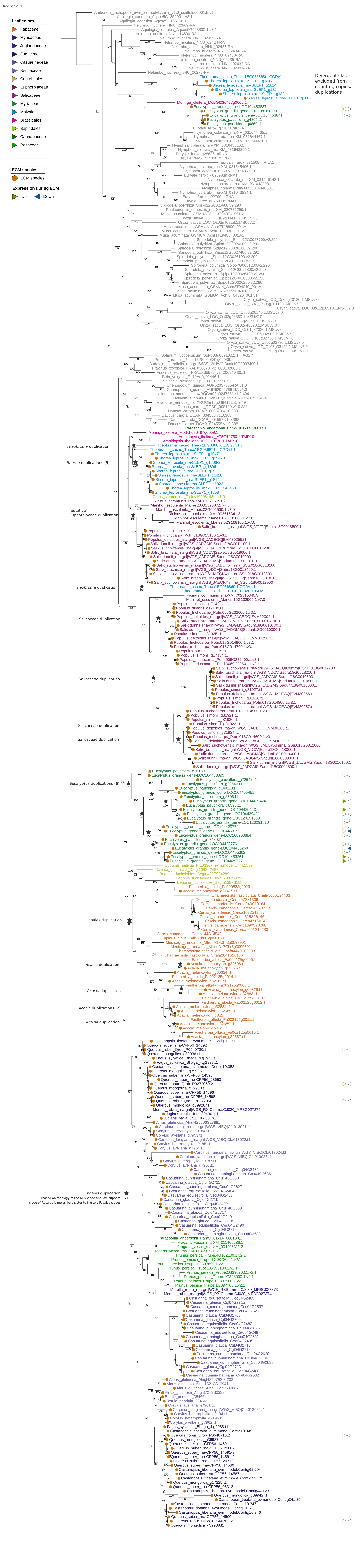
Tree scale: 1
Leaf colors
Fabaceae
Myricaceae
Juglandaceae
Fagaceae
Casuarinaceae
Betulaceae
Cucurbitales
Euphorbiaceae
Salicaceae
Myrtaceae
Malvales
Brassicales
Sapindales
Cannabaceae
Rosaceae
ECM species
ECM species
Expression during ECM
Up
Down
Divergent clade excluded from counting copies/ duplications
Theobroma duplication
Shorea duplications (9)
(putative) Euphorbiaceae duplication
Theobroma duplication
Salicaceae duplication
Salicaceae duplication
Salicaceae duplication
Salicaceae duplication
Eucalyptus duplications (6)
Fabales duplication
Acacia duplication
Acacia duplication
Acacia duplications (2)
Acacia duplication
Fagales duplication
(based on topology of the NFN-clade and low-support, clade of Rosales is more likely sister to the two Fagales clades)

Fig. S10

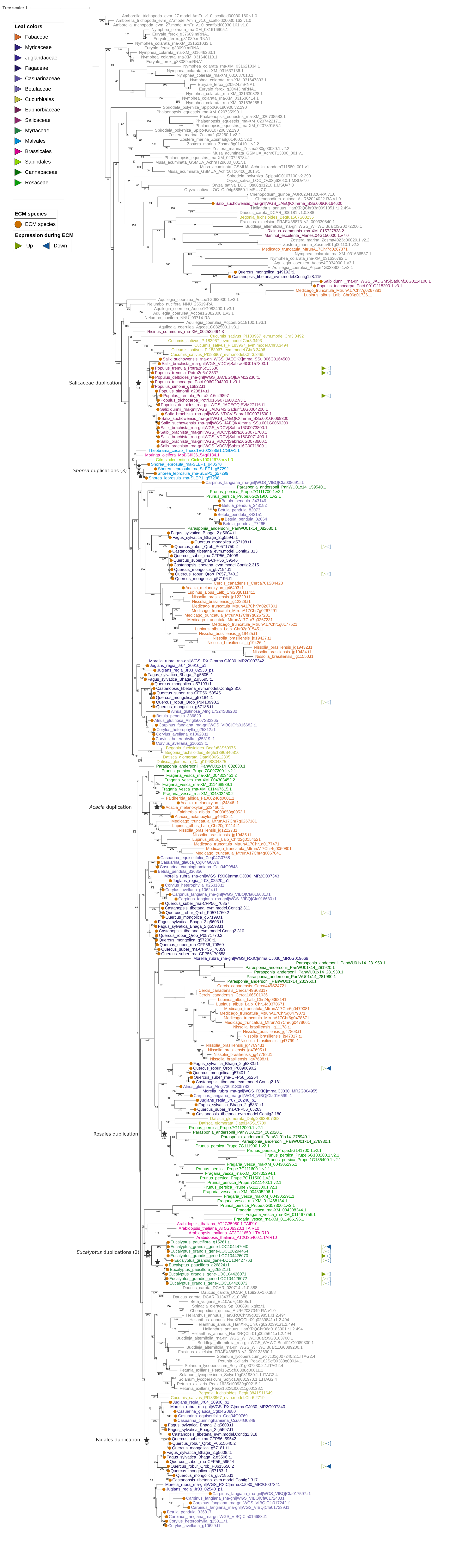

Tree scale: 1
Leaf colors
Fabaceae
Myricaceae
Juglandaceae
Fagaceae
Casuarinaceae
Betulaceae
Cucurbitales
Euphorbiaceae
Salicaceae
Myrtaceae
Malvales
Brassicales
Sapindales
Cannabaceae
Rosaceae
ECM species
ECM species
Expression during ECM
Up
Down
Salicaceae duplication
Shorea duplications (3)
Acacia duplication
Rosales duplication
Eucalyptus duplications (2)
Fagales duplication

Fig. S11

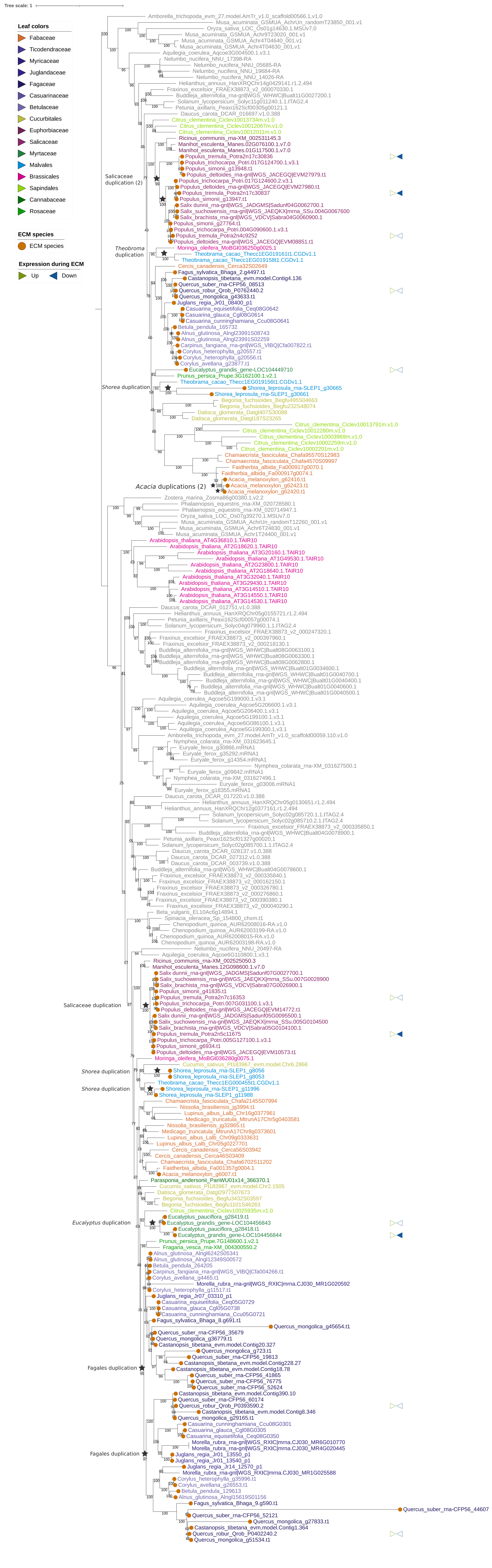

Tree scale: 1
Leaf colors
Fabaceae
Ticodendraceae
Myricaceae
Juglandaceae
Fagaceae
Casuarinaceae
Betulaceae
Cucurbitales
Euphorbiaceae
Salicaceae
Myrtaceae
Malvales
Brassicales
Sapindales
Cannabaceae
Rosaceae
ECM species
ECM species
Expression during ECM
Up
Down
Salicaceae duplication (2)
Theobroma duplication
Shorea duplication
Acacia duplications (2)
Salicaceae duplication
Shorea duplication
Shorea duplication
Eucalyptus duplication
Fagales duplication
Fagales duplication
Amborella_trichopoda_evm_27.model.AmTr_v1.0_scaffold00566.1.v1.0
Musa_acuminata_GSMUA_AchrUn_randomT23850_001.v1
Oryza_sativa_LOC_Os01g14630.1.MSUv7.0
Musa_acuminata_GSMUA_Achr9T23020_001.v1
Musa_acuminata_GSMUA_Achr4T04640_001.v1
Musa_acuminata_GSMUA_Achr4T04630_001.v1
Aquilegia_coerulea_Aqcoe3G004500.1.v3.1
Nelumbo_nucifera_NNU_17398-RA
Nelumbo_nucifera_NNU_05685-RA
Nelumbo_nucifera_NNU_19684-RA
Nelumbo_nucifera_NNU_14026-RA
Helianthus_annuus_HanXRQChr14g0429141.r1.2.494
Fraxinus_excelsior_FRAEX38873_v2_000070330.1
Buddleja_alternifolia_rna-gnl|WGS_WHWC|Bualt01G0027200.1
Solanum_lycopersicum_Solyc11g011240.1.1.ITAG2.4
Petunia_axillaris_Peaxi162Scf00305g00121.1
Daucus_carota_DCAR_016697.v1.0.388
Citrus_clementina_Ciclev10013734m.v1.0
Citrus_clementina_Ciclev10012067m.v1.0
Citrus_clementina_Ciclev10012011m.v1.0
Ricinus_communis_rna-XM_002531145.3
Manihot_esculenta_Manes.02G076100.1.v7.0
Manihot_esculenta_Manes.01G117500.1.v7.0
Populus_tremula_Potra2n17c30836
Populus_trichocarpa_Potri.017G124700.1.v3.1
Populus_simonii_g13948.t1
Populus_deltoides_rna-gnl|WGS_JACEGQ|EVM27979.t1
Populus_trichocarpa_Potri.017G124600.2.v3.1
Populus_deltoides_rna-gnl|WGS_JACEGQ|EVM27980.t1
Populus_tremula_Potra2n17c30837
Populus_simonii_g13947.t1
Salix_dunnii_rna-gnl|WGS_JADGMS|Sadunf04G0062700.1
Salix_suchowensis_rna-gnl|WGS_JAEQKX|mrna_SSu.004G0067600
Salix_brachista_rna-gnl|WGS_VDCV|Sabra04G0060900.1
Populus_simonii_g27764.t1
Populus_trichocarpa_Potri.004G090600.1.v3.1
Populus_tremula_Potra2n4c9252
Populus_deltoides_rna-gnl|WGS_JACEGQ|EVM08851.t1
Moringa_oleifera_MoBGI036250g0025.1
Theobroma_cacao_Thecc1EG019161t1.CGDv1.1
Theobroma_cacao_Thecc1EG019158t1.CGDv1.1
Cercis_canadensis_Cerca32S02649
Fagus_sylvatica_Bhaga_2.g4497.t1
Castanopsis_tibetana_evm.model.Contig4.136
Quercus_suber_rna-CFP56_08513
Quercus_robur_Qrob_P0762440.2
Quercus_mongolica_g43633.t1
Juglans_regia_Jr01_08400_p1
Casuarina_equisetifolia_Ceq08G0642
Casuarina_glauca_Cgl08G0614
Casuarina_cunninghamiana_Ccu08G0641
Betula_pendula_165732
Alnus_glutinosa_Alngl23991S08743
Alnus_glutinosa_Alngl23991S02259
Carpinus_fangiana_rna-gnl|WGS_VIBQ|Cfa007822.t1
Corylus_heterophylla_g20557.t1
Corylus_heterophylla_g20556.t1
Corylus_avellana_g23877.t1
Eucalyptus_grandis_gene-LOC104449710
Prunus_persica_Prupe.3G162100.1.v2.1
Theobroma_cacao_Thecc1EG019156t1.CGDv1.1
Shorea_leprosula_rna-SLEP1_g30665
Shorea_leprosula_rna-SLEP1_g30661
Begonia_fuchsioides_Begfu495S04663
Begonia_fuchsioides_Begfu232S48074
Datisca_glomerata_Datgl407S30088
Datisca_glomerata_Datgl187S23265
Citrus_clementina_Ciclev10013791m.v1.0
Citrus_clementina_Ciclev10012280m.v1.0
Citrus_clementina_Ciclev10003969m.v1.0
Citrus_clementina_Ciclev10002259m.v1.0
Citrus_clementina_Ciclev10002201m.v1.0
Chamaecrista_fasciculata_Chafa95570S12983
Chamaecrista_fasciculata_Chafa4570S09997
Faidherbia_albida_Fa000917g0070.1
Faidherbia_albida_Fa000917g0074.1
Acacia_melanoxylon_g62416.t1
Acacia_melanoxylon_g62423.t1
Acacia_melanoxylon_g62420.t1
Zostera_marina_Zosma86g00380.1.v2.2
Phalaenopsis_equestris_rna-XM_020728580.1
Phalaenopsis_equestris_rna-XM_020714947.1
Oryza_sativa_LOC_Os07g39270.1.MSUv7.0
Musa_acuminata_GSMUA_AchrUn_randomT12260_001.v1
Musa_acuminata_GSMUA_Achr6T24830_001.v1
Musa_acuminata_GSMUA_Achr12T24400_001.v1
Arabidopsis_thaliana_AT4G36810.1.TAIR10
Arabidopsis_thaliana_AT2G18620.1.TAIR10
Arabidopsis_thaliana_AT3G20160.1.TAIR10
Arabidopsis_thaliana_AT1G49530.1.TAIR10
Arabidopsis_thaliana_AT2G23800.1.TAIR10
Arabidopsis_thaliana_AT2G18640.1.TAIR10
Arabidopsis_thaliana_AT3G32040.1.TAIR10
Arabidopsis_thaliana_AT3G29430.1.TAIR10
Arabidopsis_thaliana_AT3G14510.1.TAIR10
Arabidopsis_thaliana_AT3G14550.1.TAIR10
Arabidopsis_thaliana_AT3G14530.1.TAIR10
Daucus_carota_DCAR_012751.v1.0.388
Helianthus_annuus_HanXRQChr05g0155721.r1.2.494
Petunia_axillaris_Peaxi162Scf00057g00074.1
Solanum_lycopersicum_Solyc04g079960.1.1.ITAG2.4
Fraxinus_excelsior_FRAEX38873_v2_000247320.1
Fraxinus_excelsior_FRAEX38873_v2_000397960.1
Fraxinus_excelsior_FRAEX38873_v2_000218130.1
Buddleja_alternifolia_rna-gnl|WGS_WHWC|Bualt08G0063100.1
Buddleja_alternifolia_rna-gnl|WGS_WHWC|Bualt08G0063300.1
Buddleja_alternifolia_rna-gnl|WGS_WHWC|Bualt08G0062800.1
Buddleja_alternifolia_rna-gnl|WGS_WHWC|Bualt01G0034600.1
Buddleja_alternifolia_rna-gnl|WGS_WHWC|Bualt01G0040700.1
Buddleja_alternifolia_rna-gnl|WGS_WHWC|Bualt01G0040400.1
Buddleja_alternifolia_rna-gnl|WGS_WHWC|Bualt01G0040600.1
Buddleja_alternifolia_rna-gnl|WGS_WHWC|Bualt01G0040500.1
Aquilegia_coerulea_Aqcoe5G199000.1.v3.1
Aquilegia_coerulea_Aqcoe5G206600.1.v3.1
Aquilegia_coerulea_Aqcoe5G206400.1.v3.1
Aquilegia_coerulea_Aqcoe5G199100.1.v3.1
Aquilegia_coerulea_Aqcoe6G086100.1.v3.1
Aquilegia_coerulea_Aqcoe5G199300.1.v3.1
Amborella_trichopoda_evm_27.model.AmTr_v1.0_scaffold00059.110.v1.0
Nymphea_colarata_rna-XM_031623645.1
Euryale_ferox_g30866.mRNA1
Euryale_ferox_g35292.mRNA1
Euryale_ferox_g14354.mRNA1
Nymphea_colarata_rna-XM_031627500.1
Euryale_ferox_g09842.mRNA1
Nymphea_colarata_rna-XM_031627496.1
Euryale_ferox_g03006.mRNA1
Euryale_ferox_g18355.mRNA1
Daucus_carota_DCAR_017220.v1.0.388
Helianthus_annuus_HanXRQChr05g0130651.r1.2.494
Helianthus_annuus_HanXRQChr12g0377161.r1.2.494
Solanum_lycopersicum_Solyc02g085720.1.1.ITAG2.4
Solanum_lycopersicum_Solyc02g085710.2.1.ITAG2.4
Fraxinus_excelsior_FRAEX38873_v2_000335850.1
Buddleja_alternifolia_rna-gnl|WGS_WHWC|Bualt04G0078900.1
Petunia_axillaris_Peaxi162Scf01327g00020.1
Solanum_lycopersicum_Solyc02g085700.1.1.ITAG2.4
Daucus_carota_DCAR_028137.v1.0.388
Daucus_carota_DCAR_027312.v1.0.388
Daucus_carota_DCAR_003739.v1.0.388
Buddleja_alternifolia_rna-gnl|WGS_WHWC|Bualt04G0078600.1
Fraxinus_excelsior_FRAEX38873_v2_000335840.1
Fraxinus_excelsior_FRAEX38873_v2_000162150.1
Fraxinus_excelsior_FRAEX38873_v2_000326780.1
Fraxinus_excelsior_FRAEX38873_v2_000276860.1
Fraxinus_excelsior_FRAEX38873_v2_000390380.1
Fraxinus_excelsior_FRAEX38873_v2_000040290.1
Beta_vulgaris_EL10Ac6g14894.1
Spinacia_oleracea_Sp_154800_chxm.t1
Chenopodium_quinoa_AUR62008016-RA.v1.0
Chenopodium_quinoa_AUR62003199-RA.v1.0
Chenopodium_quinoa_AUR62008015-RA.v1.0
Chenopodium_quinoa_AUR62003198-RA.v1.0
Nelumbo_nucifera_NNU_20497-RA
Aquilegia_coerulea_Aqcoe6G110800.1.v3.1
Ricinus_communis_rna-XM_002525050.3
Manihot_esculenta_Manes.12G098600.1.v7.0
Salix_dunnii_rna-gnl|WGS_JADGMS|Sadunf07G0027700.1
Salix_suchowensis_rna-gnl|WGS_JAEQKX|mrna_SSu.007G0028900
Salix_brachista_rna-gnl|WGS_VDCV|Sabra07G0026900.1
Populus_simonii_g41835.t1
Populus_tremula_Potra2n7c16353
Populus_trichocarpa_Potri.007G031100.1.v3.1
Populus_deltoides_rna-gnl|WGS_JACEGQ|EVM14772.t1
Salix_dunnii_rna-gnl|WGS_JADGMS|Sadunf05G0095500.1
Salix_suchowensis_rna-gnl|WGS_JAEQKX|mrna_SSu.005G0104500
Salix_brachista_rna-gnl|WGS_VDCV|Sabra05G0104100.1
Populus_tremula_Potra2n5c11675
Populus_trichocarpa_Potri.005G127100.1.v3.1
Populus_simonii_g6934.t1
Populus_deltoides_rna-gnl|WGS_JACEGQ|EVM10573.t1
Moringa_oleifera_MoBGI036280g0075.1
Cucumis_sativus_PI183967_evm.model.Chr6.2866
Shorea_leprosula_rna-SLEP1_g8056
Shorea_leprosula_rna-SLEP1_g8053
Theobroma_cacao_Thecc1EG000455t1.CGDv1.1
Shorea_leprosula_rna-SLEP1_g11996
Shorea_leprosula_rna-SLEP1_g11988
Chamaecrista_fasciculata_Chafa2145S07994
Nissolia_brasiliensis_jg3994.t1
Lupinus_albus_Lalb_Chr16g0377961
Medicago_truncatula_MtrunA17Chr5g0403581
Nissolia_brasiliensis_jg32865.t1
Medicago_truncatula_MtrunA17Chr8g0373601
Lupinus_albus_Lalb_Chr09g0333631
Lupinus_albus_Lalb_Chr05g0227701
Cercis_canadensis_Cerca56S03942
Cercis_canadensis_Cerca46S03409
Chamaecrista_fasciculata_Chafa6702S11202
Faidherbia_albida_Fa001357g0004.1
Acacia_melanoxylon_g6007.t1
Parasponia_andersonii_PanWU01x14_366370.1
Cucumis_sativus_PI183967_evm.model.Chr2.1505
Datisca_glomerata_Datgl2977S07673
Begonia_fuchsioides_Begfu3432S03597
Begonia_fuchsioides_Begfu1101S46263
Citrus_clementina_Ciclev10025935m.v1.0
Eucalyptus_pauciflora_g28419.t1
Eucalyptus_grandis_gene-LOC104456843
Eucalyptus_pauciflora_g28418.t1
Eucalyptus_grandis_gene-LOC104456844
Prunus_persica_Prupe.7G148600.1.v2.1
Fragaria_vesca_rna-XM_004300550.2
Alnus_glutinosa_Alngl6242S05341
Alnus_glutinosa_Alngl12349S00572
Betula_pendula_264205
Carpinus_fangiana_rna-gnl|WGS_VIBQ|Cfa004266.t1
Corylus_avellana_g4465.t1
Morella_rubra_rna-gnl|WGS_RXIC|mrna.CJ030_MR1G020592
Corylus_heterophylla_g11517.t1
Juglans_regia_Jr07_03310_p1
Casuarina_equisetifolia_Ceq05G0729
Casuarina_glauca_Cgl05G0738
Casuarina_cunninghamiana_Ccu05G0721
Fagus_sylvatica_Bhaga_8.g691.t1
Quercus_mongolica_g45654.t1
Quercus_suber_rna-CFP56_35679
Quercus_mongolica_g36779.t1
Castanopsis_tibetana_evm.model.Contig20.327
Quercus_mongolica_g723.t1
Quercus_suber_rna-CFP56_19813
Castanopsis_tibetana_evm.model.Contig228.27
Castanopsis_tibetana_evm.model.Contig18.78
Quercus_suber_rna-CFP56_41865
Quercus_suber_rna-CFP56_76775
Quercus_suber_rna-CFP56_52624
Castanopsis_tibetana_evm.model.Contig390.10
Quercus_suber_rna-CFP56_60174
Quercus_robur_Qrob_P0393590.2
Castanopsis_tibetana_evm.model.Contig8.346
Quercus_mongolica_g29165.t1
Casuarina_cunninghamiana_Ccu08G0301
Casuarina_glauca_Cgl08G0305
Casuarina_equisetifolia_Ceq08G0350
Morella_rubra_rna-gnl|WGS_RXIC|mrna.CJ030_MR6G010770
Morella_rubra_rna-gnl|WGS_RXIC|mrna.CJ030_MR4G020445
Juglans_regia_Jr01_13550_p1
Juglans_regia_Jr01_13540_p1
Juglans_regia_Jr14_12570_p1
Morella_rubra_rna-gnl|WGS_RXIC|mrna.CJ030_MR1G025588
Corylus_heterophylla_g35996.t1
Corylus_avellana_g26553.t1
Betula_pendula_129613
Alnus_glutinosa_Alngl15619S01156
Fagus_sylvatica_Bhaga_9.g590.t1
Quercus_suber_rna-CFP56_44607
Quercus_suber_rna-CFP56_52121
Quercus_mongolica_g27833.t1
Castanopsis_tibetana_evm.model.Contig1.364
Quercus_robur_Qrob_P0402240.2
Quercus_mongolica_g51534.t1

Fig. S12

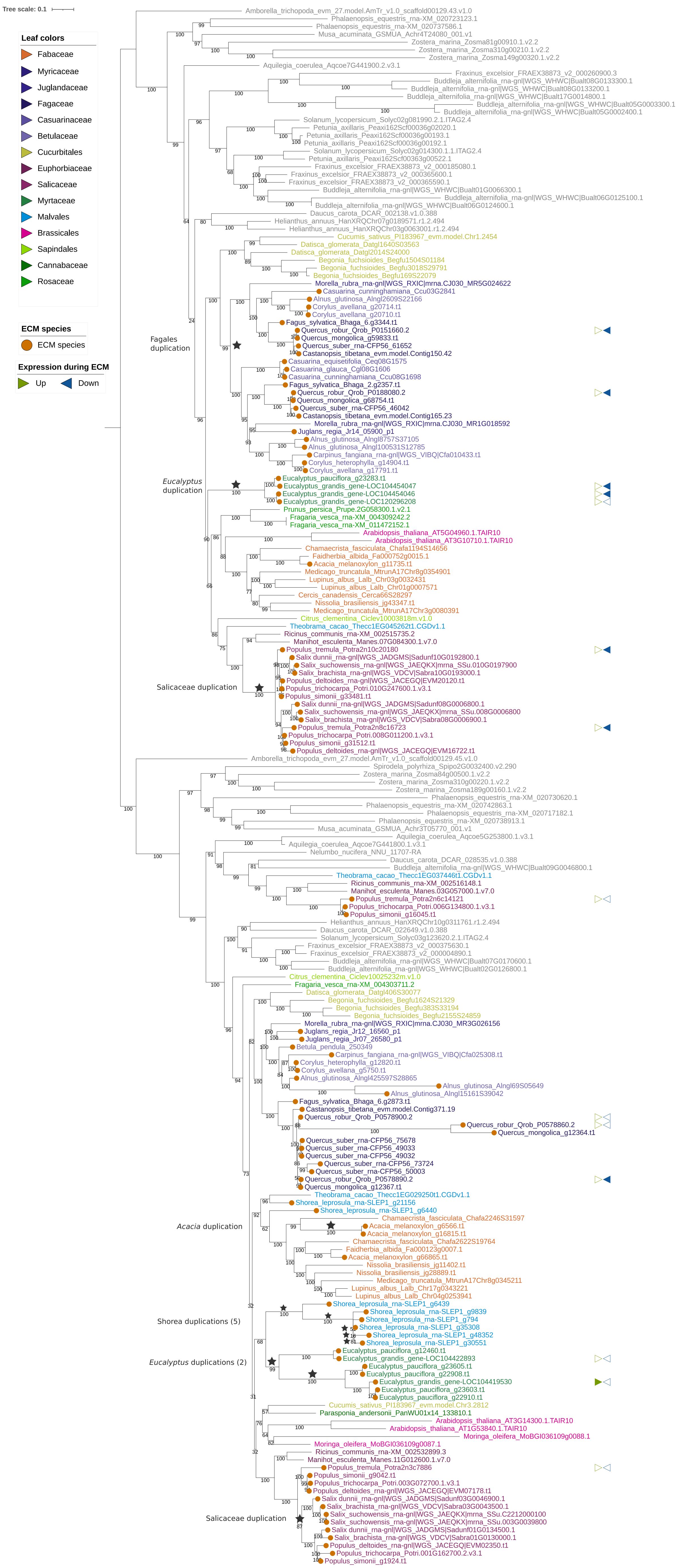

Fig. S13

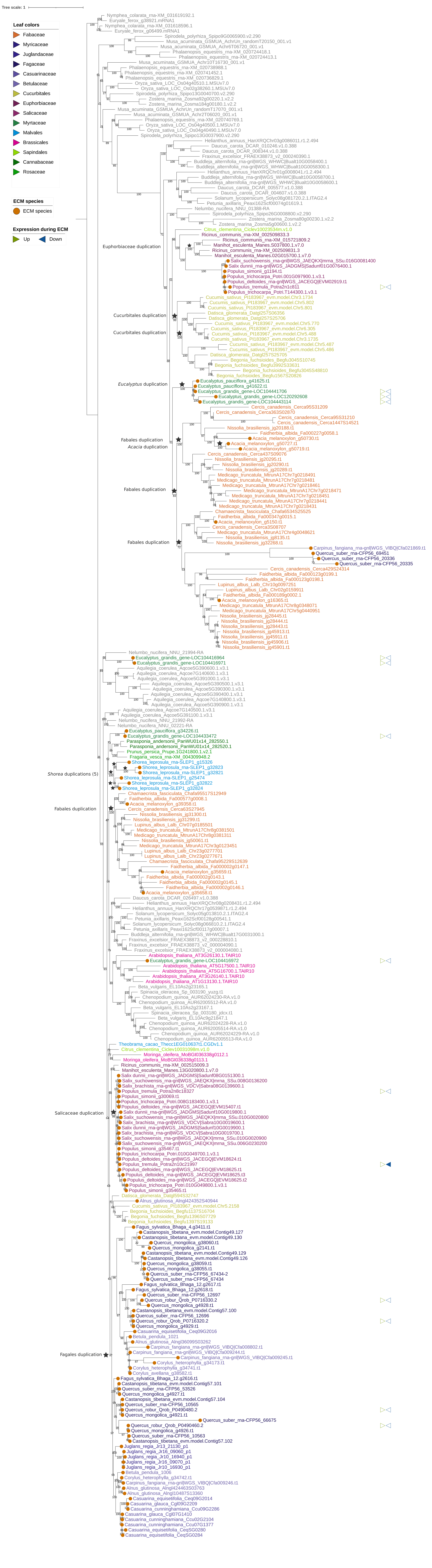

Tree scale: 1
Leaf colors
Fabaceae
Myricaceae
Juglandaceae
Fagaceae
Casuarinaceae
Betulaceae
Cucurbitales
Euphorbiaceae
Salicaceae
Myrtaceae
Malvales
Brassicales
Sapindales
Cannabaceae
Rosaceae
ECM species
ECM species
Expression during ECM
Up
Down
Euphorbiaceae duplication
Cucurbitales duplication
Cucurbitales duplication
Eucalyptus duplication
Fabales duplication
Acacia duplication
Fabales duplication
Fabales duplication
Shorea duplications (5)
Fabales duplication
Salicaceae duplication
Fagales duplication

Fig. S14

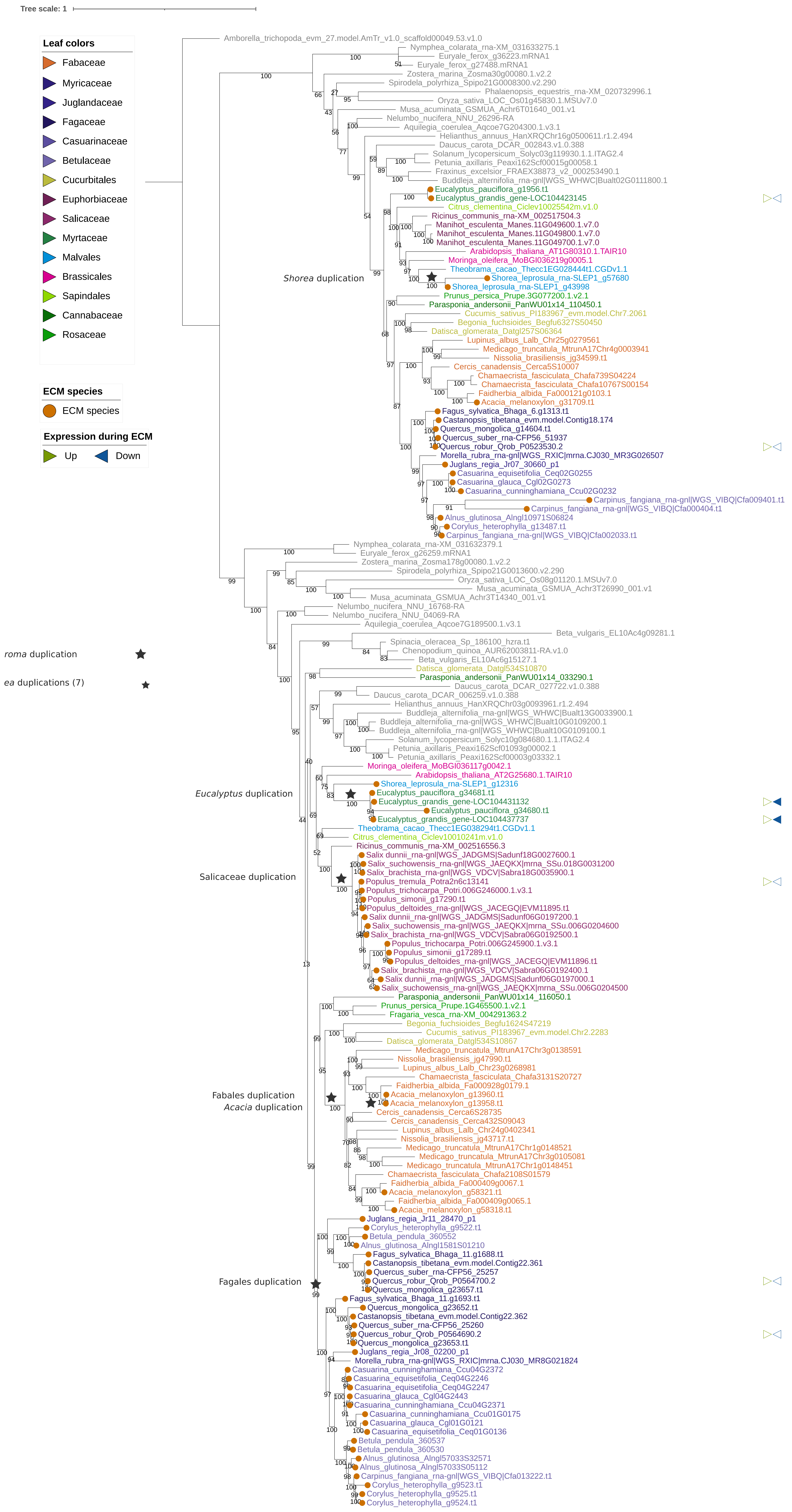

Tree scale: 1
Leaf colors
Fabaceae
Myricaceae
Juglandaceae
Fagaceae
Casuarinaceae
Betulaceae
Cucurbitales
Euphorbiaceae
Salicaceae
Myrtaceae
Malvales
Brassicales
Sapindales
Cannabaceae
Rosaceae
ECM species
ECM species
Expression during ECM
Up
Down
roma duplication
ea duplications (7)
Shorea duplication
Eucalyptus duplication
Salicaceae duplication
Fabales duplication
Acacia duplication
Fagales duplication
Amborella_trichopoda_evm_27.model.AmTr_v1.0_scaffold00049.53.v1.0
Nymphea_colarata_rna-XM_031633275.1
Euryale_ferox_g36223.mRNA1
Euryale_ferox_g27488.mRNA1
Zostera_marina_Zosma30g00080.1.v2.2
Spirodela_polyrhiza_Spipo21G0008300.v2.290
Phalaenopsis_equestris_rna-XM_020732996.1
Oryza_sativa_LOC_Os01g45830.1.MSUv7.0
Musa_acuminata_GSMUA_Achr6T01640_001.v1
Nelumbo_nucifera_NNU_26296-RA
Aquilegia_coerulea_Aqcoe7G204300.1.v3.1
Helianthus_annuus_HanXRQChr16g0500611.r1.2.494
Daucus_carota_DCAR_002843.v1.0.388
Solanum_lycopersicum_Solyc03g119930.1.1.ITAG2.4
Petunia_axillaris_Peaxi162Scf00015g00058.1
Fraxinus_excelsior_FRAEX38873_v2_000253490.1
Buddleja_alternifolia_rna-gnl|WGS_WHWC|Bualt02G0111800.1
Eucalyptus_pauciflora_g1956.t1
Eucalyptus_grandis_gene-LOC104423145
Citrus_clementina_Ciclev10025542m.v1.0
Ricinus_communis_rna-XM_002517504.3
Manihot_esculenta_Manes.11G049600.1.v7.0
Manihot_esculenta_Manes.11G049800.1.v7.0
Manihot_esculenta_Manes.11G049700.1.v7.0
Arabidopsis_thaliana_AT1G80310.1.TAIR10
Moringa_oleifera_MoBGI036219g0005.1
Theobroma_cacao_Thecc1EG028444t1.CGDv1.1
Shorea_leprosula_rna-SLEP1_g57680
Shorea_leprosula_rna-SLEP1_g43998
Prunus_persica_Prupe.3G077200.1.v2.1
Parasponia_andersonii_PanWU01x14_110450.1
Cucumis_sativus_PI183967_evm.model.Chr7.2061
Begonia_fuchsioides_Begfu6327S50450
Datisca_glomerata_Datgl257S06364
Lupinus_albus_Lalb_Chr25g0279561
Medicago_truncatula_MtrunA17Chr4g0003941
Nissolia_brasiliensis_jg34599.t1
Cercis_canadensis_Cerca5S10007
Chamaecrista_fasciculata_Chafa739S04224
Chamaecrista_fasciculata_Chafa10767S00154
Faidherbia_albida_Fa000121g0103.1
Acacia_melanoxylon_g31709.t1
Fagus_sylvatica_Bhaga_6.g1313.t1
Castanopsis_tibetana_evm.model.Contig18.174
Quercus_mongolica_g14604.t1
Quercus_suber_rna-CFP56_51937
Quercus_robur_Qrob_P0523530.2
Morella_rubra_rna-gnl|WGS_RXIC|mrna.CJ030_MR3G026507
Juglans_regia_Jr07_30660_p1
Casuarina_equisetifolia_Ceq02G0255
Casuarina_glauca_Cgl02G0273
Casuarina_cunninghamiana_Ccu02G0232
Carpinus_fangiana_rna-gnl|WGS_VIBQ|Cfa009401.t1
Carpinus_fangiana_rna-gnl|WGS_VIBQ|Cfa000404.t1
Alnus_glutinosa_Alngl10971S06824
Corylus_heterophylla_g13487.t1
Carpinus_fangiana_rna-gnl|WGS_VIBQ|Cfa002033.t1
Nymphea_colarata_rna-XM_031632379.1
Euryale_ferox_g26259.mRNA1
Zostera_marina_Zosma178g00080.1.v2.2
Spirodela_polyrhiza_Spipo21G0013600.v2.290
Oryza_sativa_LOC_Os08g01120.1.MSUv7.0
Musa_acuminata_GSMUA_Achr3T26990_001.v1
Musa_acuminata_GSMUA_Achr3T14340_001.v1
Nelumbo_nucifera_NNU_16768-RA
Nelumbo_nucifera_NNU_04069-RA
Aquilegia_coerulea_Aqcoe7G189500.1.v3.1
Beta_vulgaris_EL10Ac4g09281.1
Spinacia_oleracea_Sp_186100_hzra.t1
Chenopodium_quinoa_AUR62003811-RA.v1.0
Beta_vulgaris_EL10Ac6g15127.1
Datisca_glomerata_Datgl534S10870
Parasponia_andersonii_PanWU01x14_033290.1
Daucus_carota_DCAR_027722.v1.0.388
Daucus_carota_DCAR_006259.v1.0.388
Helianthus_annuus_HanXRQChr03g0093961.r1.2.494
Buddleja_alternifolia_rna-gnl|WGS_WHWC|Bualt13G0033900.1
Buddleja_alternifolia_rna-gnl|WGS_WHWC|Bualt10G0109200.1
Buddleja_alternifolia_rna-gnl|WGS_WHWC|Bualt10G0109100.1
Solanum_lycopersicum_Solyc10g084680.1.1.ITAG2.4
Petunia_axillaris_Peaxi162Scf01093g00002.1
Petunia_axillaris_Peaxi162Scf00003g03332.1
Moringa_oleifera_MoBGI036117g0042.1
Arabidopsis_thaliana_AT2G25680.1.TAIR10
Shorea_leprosula_rna-SLEP1_g12316
Eucalyptus_pauciflora_g34681.t1
Eucalyptus_grandis_gene-LOC104431132
Eucalyptus_pauciflora_g34680.t1
Eucalyptus_grandis_gene-LOC104437737
Theobroma_cacao_Thecc1EG038294t1.CGDv1.1
Citrus_clementina_Ciclev10010241m.v1.0
Ricinus_communis_rna-XM_002516556.3
Salix_dunnii_rna-gnl|WGS_JADGMS|Sadunf18G0027600.1
Salix_suchowensis_rna-gnl|WGS_JAEQKX|mrna_SSu.018G0031200
Salix_brachista_rna-gnl|WGS_VDCV|Sabra18G0035900.1
Populus_tremula_Potra2n6c13141
Populus_trichocarpa_Potri.006G246000.1.v3.1
Populus_simonii_g17290.t1
Populus_deltoides_rna-gnl|WGS_JACEGQ|EVM11895.t1
Salix_dunnii_rna-gnl|WGS_JADGMS|Sadunf06G0197200.1
Salix_suchowensis_rna-gnl|WGS_JAEQKX|mrna_SSu.006G0204600
Salix_brachista_rna-gnl|WGS_VDCV|Sabra06G0192500.1
Populus_trichocarpa_Potri.006G245900.1.v3.1
Populus_simonii_g17289.t1
Populus_deltoides_rna-gnl|WGS_JACEGQ|EVM11896.t1
Salix_brachista_rna-gnl|WGS_VDCV|Sabra06G0192400.1
Salix_dunnii_rna-gnl|WGS_JADGMS|Sadunf06G0197000.1
Salix_suchowensis_rna-gnl|WGS_JAEQKX|mrna_SSu.006G0204500
Parasponia_andersonii_PanWU01x14_116050.1
Prunus_persica_Prupe.1G465500.1.v2.1
Fragaria_vesca_rna-XM_004291363.2
Begonia_fuchsioides_Begfu1624S47219
Cucumis_sativus_PI183967_evm.model.Chr2.2283
Datisca_glomerata_Datgl534S10867
Medicago_truncatula_MtrunA17Chr3g0138591
Nissolia_brasiliensis_jg47990.t1
Lupinus_albus_Lalb_Chr23g0268981
Chamaecrista_fasciculata_Chafa3131S20727
Faidherbia_albida_Fa000928g0179.1
Acacia_melanoxylon_g13960.t1
Acacia_melanoxylon_g13958.t1
Cercis_canadensis_Cerca6S28735
Cercis_canadensis_Cerca432S09043
Lupinus_albus_Lalb_Chr24g0402341
Nissolia_brasiliensis_jg43717.t1
Medicago_truncatula_MtrunA17Chr1g0148521
Medicago_truncatula_MtrunA17Chr3g0105081
Medicago_truncatula_MtrunA17Chr1g0148451
Chamaecrista_fasciculata_Chafa2108S01579
Faidherbia_albida_Fa000409g0067.1
Acacia_melanoxylon_g58321.t1
Faidherbia_albida_Fa000409g0065.1
Acacia_melanoxylon_g58318.t1
Juglans_regia_Jr11_28470_p1
Corylus_heterophylla_g9522.t1
Betula_pendula_360552
Alnus_glutinosa_Alngl1581S01210
Fagus_sylvatica_Bhaga_11.g1688.t1
Castanopsis_tibetana_evm.model.Contig22.361
Quercus_suber_rna-CFP56_25257
Quercus_robur_Qrob_P0564700.2
Quercus_mongolica_g23657.t1
Fagus_sylvatica_Bhaga_11.g1693.t1
Quercus_mongolica_g23652.t1
Castanopsis_tibetana_evm.model.Contig22.362
Quercus_suber_rna-CFP56_25260
Quercus_robur_Qrob_P0564690.2
Quercus_mongolica_g23653.t1
Juglans_regia_Jr08_02200_p1
Morella_rubra_rna-gnl|WGS_RXIC|mrna.CJ030_MR8G021824
Casuarina_cunninghamiana_Ccu04G2372
Casuarina_equisetifolia_Ceq04G2246
Casuarina_equisetifolia_Ceq04G2247
Casuarina_glauca_Cgl04G2443
Casuarina_cunninghamiana_Ccu04G2371
Casuarina_cunninghamiana_Ccu01G0175
Casuarina_glauca_Cgl01G0121
Casuarina_equisetifolia_Ceq01G0136
Betula_pendula_360537
Betula_pendula_360530
Alnus_glutinosa_Alngl57033S32571
Alnus_glutinosa_Alngl57033S05112
Carpinus_fangiana_rna-gnl|WGS_VIBQ|Cfa013222.t1
Corylus_heterophylla_g9523.t1
Corylus_heterophylla_g9525.t1
Corylus_heterophylla_g9524.t1

Fig. S15

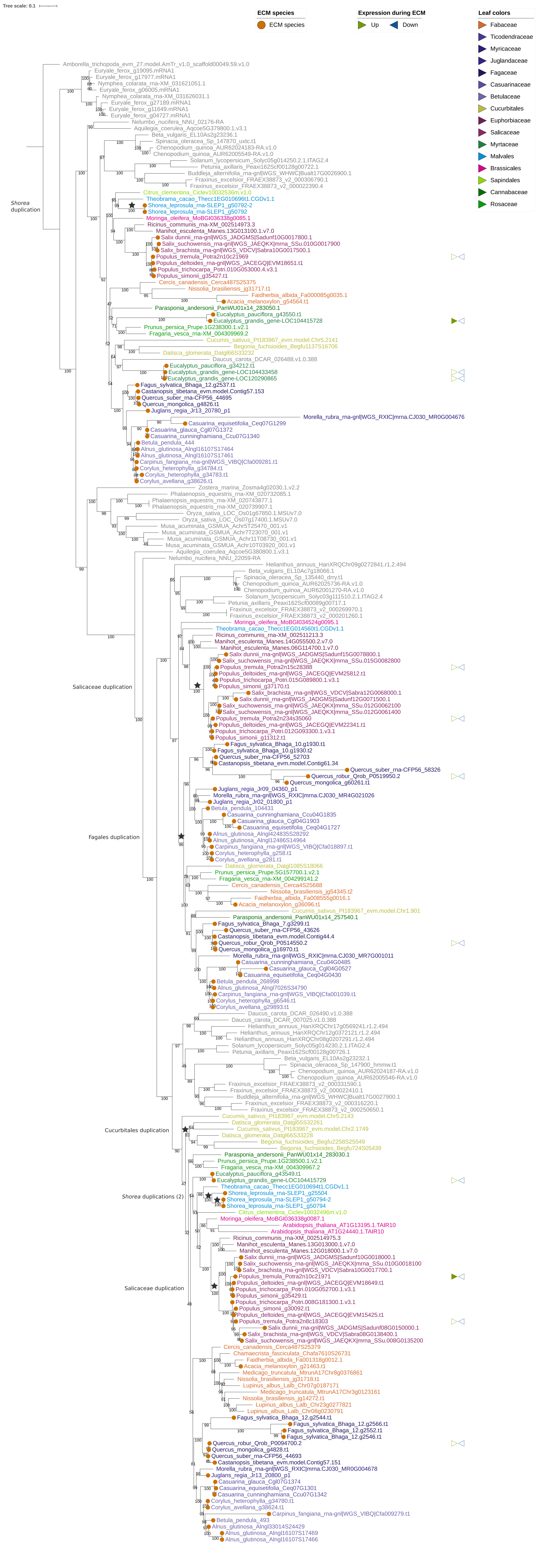

Tree scale: 0.1
ECM species
ECM species
Expression during ECM
Up
Down
Leaf colors
Fabaceae
Ticodendraceae
Myricaceae
Juglandaceae
Fagaceae
Casuarinaceae
Betulaceae
Cucurbitales
Euphorbiaceae
Salicaceae
Myrtaceae
Malvales
Brassicales
Sapindales
Cannabaceae
Rosaceae
Shorea duplication
Salicaceae duplication
Fagales duplication
Cucurbitales duplication
Shorea duplications (2)
Salicaceae duplication

Fig. S16

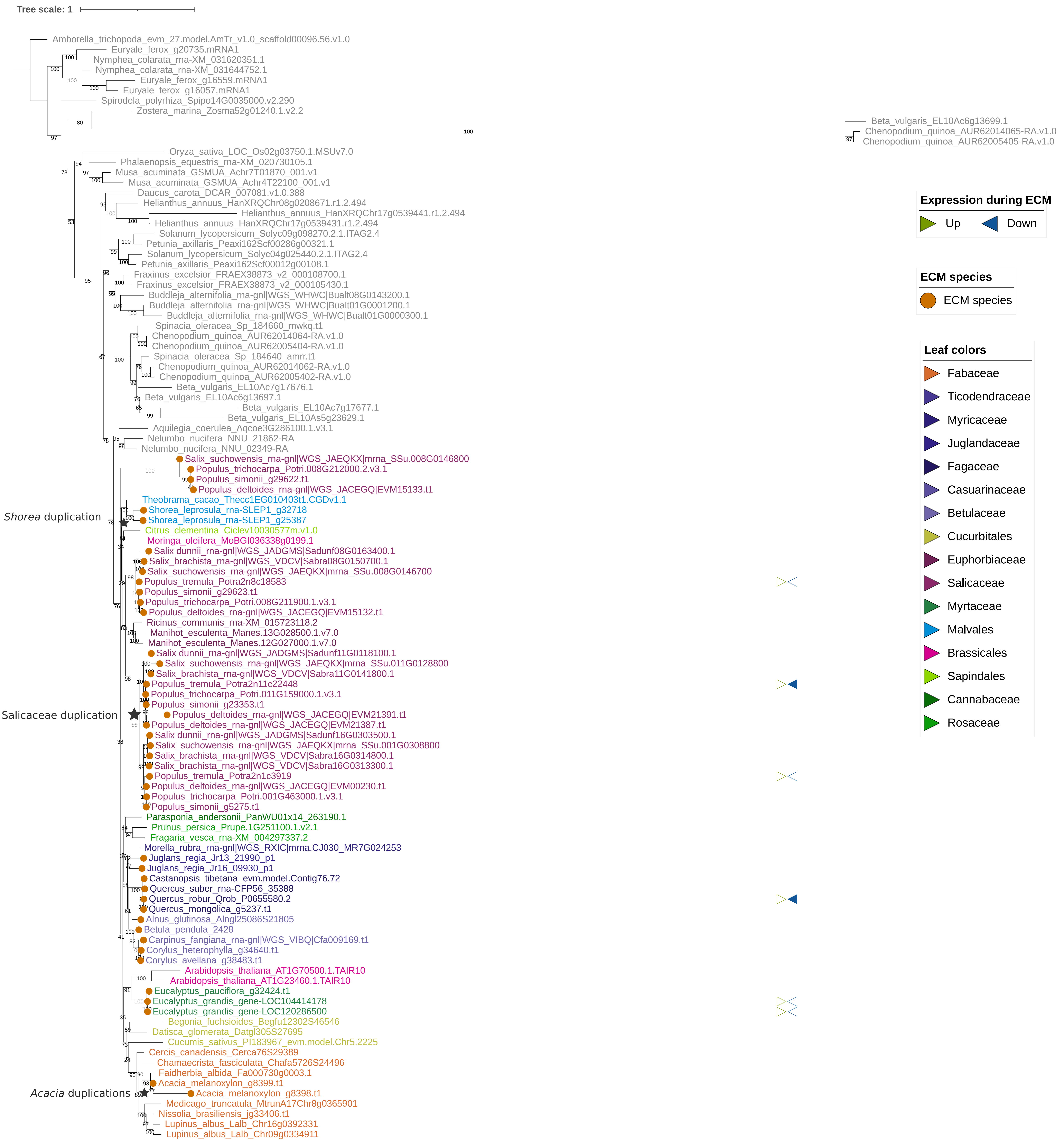
Tree scale: 1
Expression during ECM
Up
Down
ECM species
ECM species
Leaf colors
Fabaceae
Ticodendraceae
Myricaceae
Juglandaceae
Fagaceae
Casuarinaceae
Betulaceae
Cucurbitales
Euphorbiaceae
Salicaceae
Myrtaceae
Malvales
Brassicales
Sapindales
Cannabaceae
Rosaceae
Shorea duplication
Salicaceae duplication
Acacia duplications
Amborella_trichopoda_evm_27.model.AmTr_v1.0_scaffold00096.56.v1.0
Euryale_ferox_g20735.mRNA1
Nymphea_colarata_rna-XM_031620351.1
Nymphea_colarata_rna-XM_031644752.1
Euryale_ferox_g16559.mRNA1
Euryale_ferox_g16057.mRNA1
Spirodela_polyrhiza_Spipo14G0035000.v2.290
Zostera_marina_Zosma52g01240.1.v2.2
Beta_vulgaris_EL10Ac6g13699.1
Chenopodium_quinoa_AUR62014065-RA.v1.0
Chenopodium_quinoa_AUR62005405-RA.v1.0
Oryza_sativa_LOC_Os02g03750.1.MSUv7.0
Phalaenopsis_equestris_rna-XM_020730105.1
Musa_acuminata_GSMUA_Achr7T01870_001.v1
Musa_acuminata_GSMUA_Achr4T22100_001.v1
Daucus_carota_DCAR_007081.v1.0.388
Helianthus_annuus_HanXRQChr08g0208671.r1.2.494
Helianthus_annuus_HanXRQChr17g0539441.r1.2.494
Helianthus_annuus_HanXRQChr17g0539431.r1.2.494
Solanum_lycopersicum_Solyc09g098270.2.1.ITAG2.4
Petunia_axillaris_Peaxi162Scf00286g00321.1
Solanum_lycopersicum_Solyc04g025440.2.1.ITAG2.4
Petunia_axillaris_Peaxi162Scf00012g00108.1
Fraxinus_excelsior_FRAEX38873_v2_000108700.1
Fraxinus_excelsior_FRAEX38873_v2_000105430.1
Buddleja_alternifolia_rna-gnl|WGS_WHWC|Bualt08G0143200.1
Buddleja_alternifolia_rna-gnl|WGS_WHWC|Bualt01G0001200.1
Buddleja_alternifolia_rna-gnl|WGS_WHWC|Bualt01G0000300.1
Spinacia_oleracea_Sp_184660_mwkq.t1
Chenopodium_quinoa_AUR62014064-RA.v1.0
Chenopodium_quinoa_AUR62005404-RA.v1.0
Spinacia_oleracea_Sp_184640_amrr.t1
Chenopodium_quinoa_AUR62014062-RA.v1.0
Chenopodium_quinoa_AUR62005402-RA.v1.0
Beta_vulgaris_EL10Ac7g17676.1
Beta_vulgaris_EL10Ac6g13697.1
Beta_vulgaris_EL10Ac7g17677.1
Beta_vulgaris_EL10As5g23629.1
Aquilegia_coerulea_Aqcoe3G286100.1.v3.1
Nelumbo_nucifera_NNU_21862-RA
Nelumbo_nucifera_NNU_02349-RA
Salix_suchowensis_rna-gnl|WGS_JAEQKX|mrna_SSu.008G0146800
Populus_trichocarpa_Potri.008G212000.2.v3.1
Populus_simonii_g29622.t1
Populus_deltoides_rna-gnl|WGS_JACEGQ|EVM15133.t1
Theobrama_cacao_Thecc1EG010403t1.CGDv1.1
Shorea_leprosula_rna-SLEP1_g32718
Shorea_leprosula_rna-SLEP1_g25387
Citrus_clementina_Ciclev10030577m.v1.0
Moringa_oleifera_MoBGI036338g0199.1
Salix dunnii_rna-gnl|WGS_JADGMS|Sadunf08G0163400.1
Salix_brachista_rna-gnl|WGS_VDCV|Sabra08G0150700.1
Salix_suchowensis_rna-gnl|WGS_JAEQKX|mrna_SSu.008G0146700
Populus_tremula_Potra2n8c18583
Populus_simonii_g29623.t1
Populus_trichocarpa_Potri.008G211900.1.v3.1
Populus_deltoides_rna-gnl|WGS_JACEGQ|EVM15132.t1
Ricinus_communis_rna-XM_015723118.2
Manihot_esculenta_Manes.13G028500.1.v7.0
Manihot_esculenta_Manes.12G027000.1.v7.0
Salix dunnii_rna-gnl|WGS_JADGMS|Sadunf11G0118100.1
Salix_suchowensis_rna-gnl|WGS_JAEQKX|mrna_SSu.011G0128800
Salix_brachista_rna-gnl|WGS_VDCV|Sabra11G0141800.1
Populus_tremula_Potra2n11c22448
Populus_trichocarpa_Potri.011G159000.1.v3.1
Populus_simonii_g23353.t1
Populus_deltoides_rna-gnl|WGS_JACEGQ|EVM21391.t1
Populus_deltoides_rna-gnl|WGS_JACEGQ|EVM21387.t1
Salix dunnii_rna-gnl|WGS_JADGMS|Sadunf16G0303500.1
Salix_suchowensis_rna-gnl|WGS_JAEQKX|mrna_SSu.001G0308800
Salix_brachista_rna-gnl|WGS_VDCV|Sabra16G0314800.1
Salix_brachista_rna-gnl|WGS_VDCV|Sabra16G0313300.1
Populus_tremula_Potra2n1c3919
Populus_deltoides_rna-gnl|WGS_JACEGQ|EVM00230.t1
Populus_trichocarpa_Potri.001G463000.1.v3.1
Populus_simonii_g5275.t1
Parasponia_andersonii_PanWU01x14_263190.1
Prunus_persica_Prupe.1G251100.1.v2.1
Fragaria_vesca_rna-XM_004297337.2
Morella_rubra_rna-gnl|WGS_RXIC|mrna.CJ030_MR7G024253
Juglans_regia_Jr13_21990_p1
Juglans_regia_Jr16_09930_p1
Castanopsis_tibetana_evm.model.Contig76.72
Quercus_suber_rna-CFP56_35388
Quercus_robur_Qrob_P0655580.2
Quercus_mongolica_g5237.t1
Alnus_glutinosa_Alngl25086S21805
Betula_pendula_2428
Carpinus_fangiana_rna-gnl|WGS_VIBQ|Cfa009169.t1
Corylus_heterophylla_g34640.t1
Corylus_avellana_g38483.t1
Arabidopsis_thaliana_AT1G70500.1.TAIR10
Arabidopsis_thaliana_AT1G23460.1.TAIR10
Eucalyptus_pauciflora_g32424.t1
Eucalyptus_grandis_gene-LOC104414178
Eucalyptus_grandis_gene-LOC120286500
Begonia_fuchsioides_Begfu12302S46546
Datisca_glomerata_Datgl305S27695
Cucumis_sativus_PI183967_evm.model.Chr5.2225
Cercis_canadensis_Cerca76S29389
Chamaecrista_fasciculata_Chafa5726S24496
Faidherbia_albida_Fa000730g0003.1
Acacia_melanoxylon_g8399.t1
Acacia_melanoxylon_g8398.t1
Medicago_truncatula_MtrunA17Chr8g0365901
Nissolia_brasiliensis_jg33406.t1
Lupinus_albus_Lalb_Chr16g0392331
Lupinus_albus_Lalb_Chr09g0334911

Fig. S17

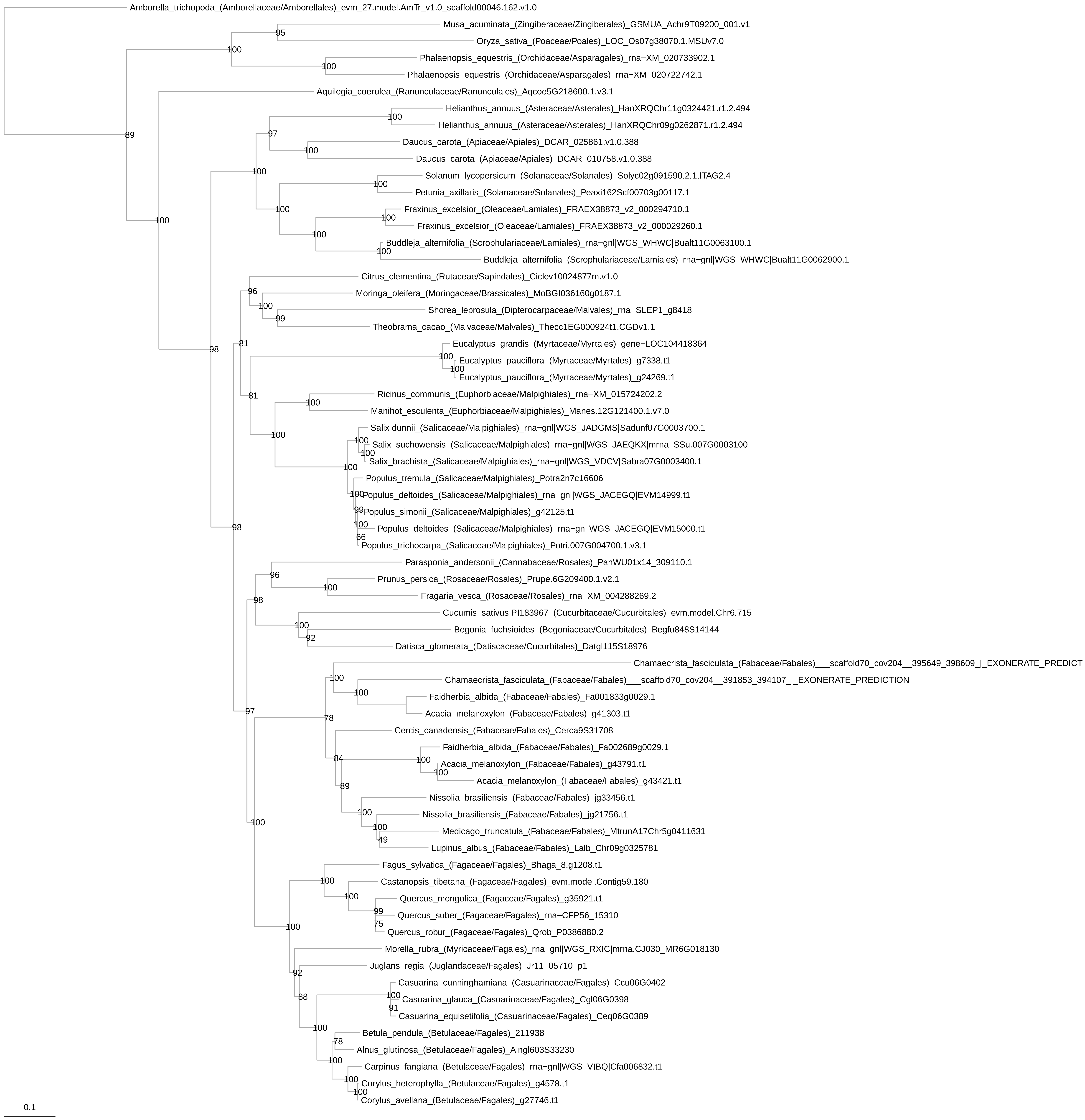
Amborella_trichopoda_(Amborellaceae/Amborellales)_evm_27.model.AmTr_v1.0_scaffold00046.162.v1.0
Musa_acuminata_(Zingiberaceae/Zingiberales)_GSMUA_Achr9T09200_001.v1
Oryza_sativa_(Poaceae/Poales)_LOC_Os07g38070.1.MSUv7.0
Phalaenopsis_equestris_(Orchidaceae/Asparagales)_rna−XM_020733902.1
Phalaenopsis_equestris_(Orchidaceae/Asparagales)_rna−XM_020722742.1
Aquilegia_coerulea_(Ranunculaceae/Ranunculales)_Aqcoe5G218600.1.v3.1
Helianthus_annuus_(Asteraceae/Asterales)_HanXRQChr11g0324421.r1.2.494
Helianthus_annuus_(Asteraceae/Asterales)_HanXRQChr09g0262871.r1.2.494
Daucus_carota_(Apiaceae/Apiales)_DCAR_025861.v1.0.388
Daucus_carota_(Apiaceae/Apiales)_DCAR_010758.v1.0.388
Solanum_lycopersicum_(Solanaceae/Solanales)_Solyc02g091590.2.1.ITAG2.4
Petunia_axillaris_(Solanaceae/Solanales)_Peaxi162Scf00703g00117.1
Fraxinus_excelsior_(Oleaceae/Lamiales)_FRAEX38873_v2_000294710.1
Fraxinus_excelsior_(Oleaceae/Lamiales)_FRAEX38873_v2_000029260.1
Buddleja_alternifolia_(Scrophulariaceae/Lamiales)_rna−gnl|WGS_WHWC|Bualt11G0063100.1
Buddleja_alternifolia_(Scrophulariaceae/Lamiales)_rna−gnl|WGS_WHWC|Bualt11G0062900.1
Citrus_clementina_(Rutaceae/Sapindales)_Ciclev10024877m.v1.0
Moringa_oleifera_(Moringaceae/Brassicales)_MoBGI036160g0187.1
Shorea_leprosula_(Dipterocarpaceae/Malvales)_rna−SLEP1_g8418
Theobrama_cacao_(Malvaceae/Malvales)_Thecc1EG000924t1.CGDv1.1
Eucalyptus_grandis_(Myrtaceae/Myrtales)_gene−LOC104418364
Eucalyptus_pauciflora_(Myrtaceae/Myrtales)_g7338.t1
Eucalyptus_pauciflora_(Myrtaceae/Myrtales)_g24269.t1
Ricinus_communis_(Euphorbiaceae/Malpighiales)_rna−XM_015724202.2
Manihot_esculenta_(Euphorbiaceae/Malpighiales)_Manes.12G121400.1.v7.0
Salix dunnii_(Salicaceae/Malpighiales)_rna−gnl|WGS_JADGMS|Sadunf07G0003700.1
Salix_suchowensis_(Salicaceae/Malpighiales)_rna−gnl|WGS_JAEQKX|mrna_SSu.007G0003100
Salix_brachista_(Salicaceae/Malpighiales)_rna−gnl|WGS_VDCV|Sabra07G0003400.1
Populus_tremula_(Salicaceae/Malpighiales)_Potra2n7c16606
Populus_deltoides_(Salicaceae/Malpighiales)_rna−gnl|WGS_JACEGQ|EVM14999.t1
Populus_simonii_(Salicaceae/Malpighiales)_g42125.t1
Populus_deltoides_(Salicaceae/Malpighiales)_rna−gnl|WGS_JACEGQ|EVM15000.t1
Populus_trichocarpa_(Salicaceae/Malpighiales)_Potri.007G004700.1.v3.1
Parasponia_andersonii_(Cannabaceae/Rosales)_PanWU01x14_309110.1
Prunus_persica_(Rosaceae/Rosales)_Prupe.6G209400.1.v2.1
Fragaria_vesca_(Rosaceae/Rosales)_rna−XM_004288269.2
Cucumis_sativus PI183967_(Cucurbitaceae/Cucurbitales)_evm.model.Chr6.715
Begonia_fuchsioides_(Begoniaceae/Cucurbitales)_Begfu848S14144
Datisca_glomerata_(Datiscaceae/Cucurbitales)_Datgl115S18976
Chamaecrista_fasciculata_(Fabaceae/Fabales)___scaffold70_cov204__395649_398609_|_EXONERATE_PREDICT
Chamaecrista_fasciculata_(Fabaceae/Fabales)___scaffold70_cov204__391853_394107_|_EXONERATE_PREDICTION
Faidherbia_albida_(Fabaceae/Fabales)_Fa001833g0029.1
Acacia_melanoxylon_(Fabaceae/Fabales)_g41303.t1
Cercis_canadensis_(Fabaceae/Fabales)_Cerca9S31708
Faidherbia_albida_(Fabaceae/Fabales)_Fa002689g0029.1
Acacia_melanoxylon_(Fabaceae/Fabales)_g43791.t1
Acacia_melanoxylon_(Fabaceae/Fabales)_g43421.t1
Nissolia_brasiliensis_(Fabaceae/Fabales)_jg33456.t1
Nissolia_brasiliensis_(Fabaceae/Fabales)_jg21756.t1
Medicago_truncatula_(Fabaceae/Fabales)_MtrunA17Chr5g0411631
Lupinus_albus_(Fabaceae/Fabales)_Lalb_Chr09g0325781
Fagus_sylvatica_(Fagaceae/Fagales)_Bhaga_8.g1208.t1
Castanopsis_tibetana_(Fagaceae/Fagales)_evm.model.Contig59.180
Quercus_mongolica_(Fagaceae/Fagales)_g35921.t1
Quercus_suber_(Fagaceae/Fagales)_rna−CFP56_15310
Quercus_robur_(Fagaceae/Fagales)_Qrob_P0386880.2
Morella_rubra_(Myricaceae/Fagales)_rna−gnl|WGS_RXIC|mrna.CJ030_MR6G018130
Juglans_regia_(Juglandaceae/Fagales)_Jr11_05710_p1
Casuarina_cunninghamiana_(Casuarinaceae/Fagales)_Ccu06G0402
Casuarina_glauca_(Casuarinaceae/Fagales)_Cgl06G0398
Casuarina_equisetifolia_(Casuarinaceae/Fagales)_Ceq06G0389
Betula_pendula_(Betulaceae/Fagales)_211938
Alnus_glutinosa_(Betulaceae/Fagales)_Alngl603S33230
Carpinus_fangiana_(Betulaceae/Fagales)_rna−gnl|WGS_VIBQ|Cfa006832.t1
Corylus_heterophylla_(Betulaceae/Fagales)_g4578.t1
Corylus_avellana_(Betulaceae/Fagales)_g27746.t1
0.1

Fig. S18

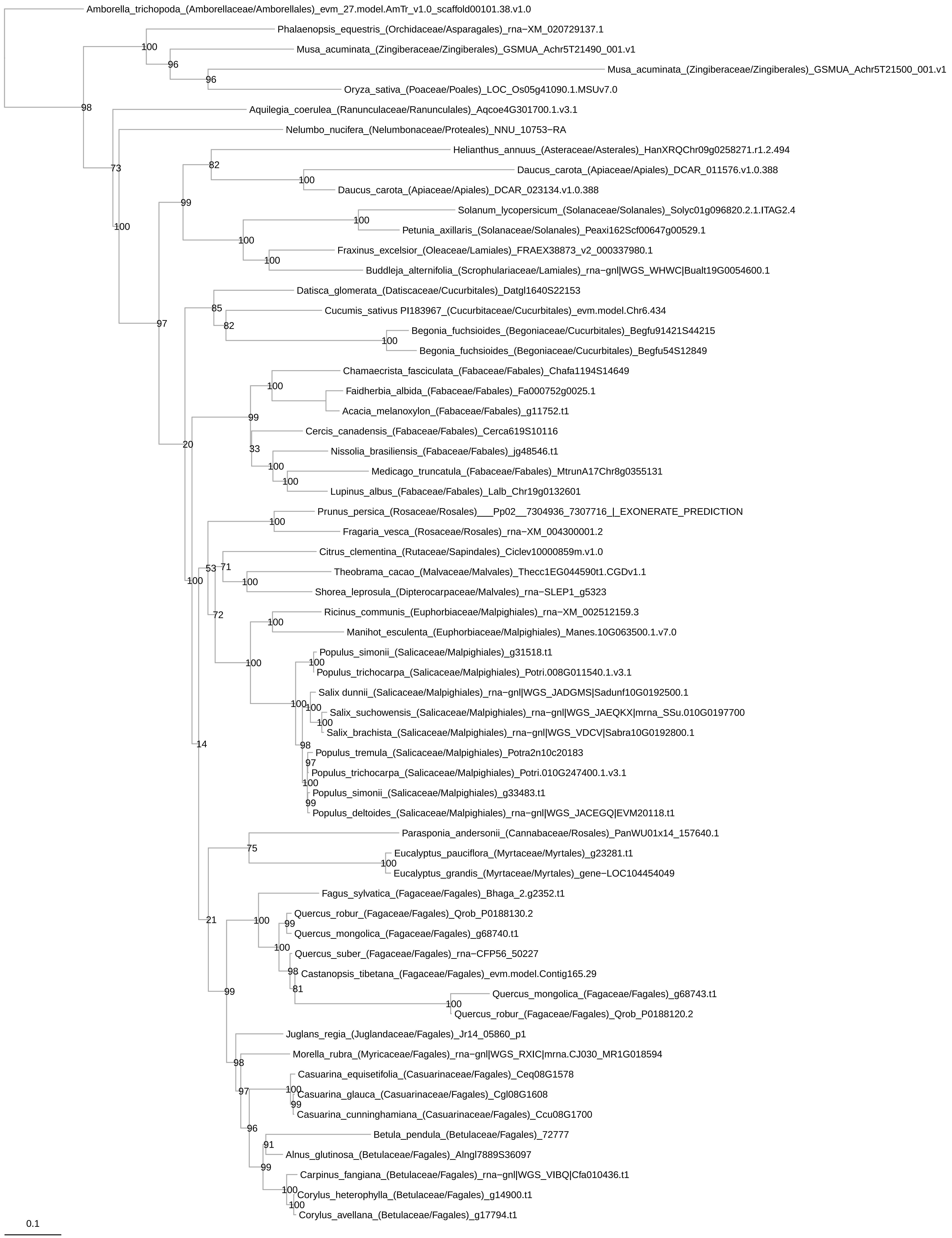

Amborella_trichopoda_(Amborellaceae/Amborellales)_evm_27.model.AmTr_v1.0_scaffold00101.38.v1.0
Phalaenopsis_equestris_(Orchidaceae/Asparagales)_rna−XM_020729137.1
Musa_acuminata_(Zingiberaceae/Zingiberales)_GSMUA_Achr5T21490_001.v1
Musa_acuminata_(Zingiberaceae/Zingiberales)_GSMUA_Achr5T21500_001.v1
Oryza_sativa_(Poaceae/Poales)_LOC_Os05g41090.1.MSUv7.0
Aquilegia_coerulea_(Ranunculaceae/Ranunculales)_Aqcoe4G301700.1.v3.1
Nelumbo_nucifera_(Nelumbonaceae/Proteales)_NNU_10753−RA
Helianthus_annuus_(Asteraceae/Asterales)_HanXRQChr09g0258271.r1.2.494
Daucus_carota_(Apiaceae/Apiales)_DCAR_011576.v1.0.388
Daucus_carota_(Apiaceae/Apiales)_DCAR_023134.v1.0.388
Solanum_lycopersicum_(Solanaceae/Solanales)_Solyc01g096820.2.1.ITAG2.4
Petunia_axillaris_(Solanaceae/Solanales)_Peaxi162Scf00647g00529.1
Fraxinus_excelsior_(Oleaceae/Lamiales)_FRAEX38873_v2_000337980.1
Buddleja_alternifolia_(Scrophulariaceae/Lamiales)_rna−gnl|WGS_WHWC|Bualt19G0054600.1
Datisca_glomerata_(Datiscaceae/Cucurbitales)_Datgl1640S22153
Cucumis_sativus PI183967_(Cucurbitaceae/Cucurbitales)_evm.model.Chr6.434
Begonia_fuchsioides_(Begoniaceae/Cucurbitales)_Begfu91421S44215
Begonia_fuchsioides_(Begoniaceae/Cucurbitales)_Begfu54S12849
Chamaecrista_fasciculata_(Fabaceae/Fabales)_Chafa1194S14649
Faidherbia_albida_(Fabaceae/Fabales)_Fa000752g0025.1
Acacia_melanoxylon_(Fabaceae/Fabales)_g11752.t1
Cercis_canadensis_(Fabaceae/Fabales)_Cerca619S10116
Nissolia_brasiliensis_(Fabaceae/Fabales)_jg48546.t1
Medicago_truncatula_(Fabaceae/Fabales)_MtrunA17Chr8g0355131
Lupinus_albus_(Fabaceae/Fabales)_Lalb_Chr19g0132601
Prunus_persica_(Rosaceae/Rosales)___Pp02__7304936_7307716_|_EXONERATE_PREDICTION
Fragaria_vesca_(Rosaceae/Rosales)_rna−XM_004300001.2
Citrus_clementina_(Rutaceae/Sapindales)_Ciclev10000859m.v1.0
Theobrama_cacao_(Malvaceae/Malvales)_Thecc1EG044590t1.CGDv1.1
Shorea_leprosula_(Dipterocarpaceae/Malvales)_rna−SLEP1_g5323
Ricinus_communis_(Euphorbiaceae/Malpighiales)_rna−XM_002512159.3
Manihot_esculenta_(Euphorbiaceae/Malpighiales)_Manes.10G063500.1.v7.0
Populus_simonii_(Salicaceae/Malpighiales)_g31518.t1
Populus_trichocarpa_(Salicaceae/Malpighiales)_Potri.008G011540.1.v3.1
Salix dunnii_(Salicaceae/Malpighiales)_rna−gnl|WGS_JADGMS|Sadunf10G0192500.1
Salix_suchowensis_(Salicaceae/Malpighiales)_rna−gnl|WGS_JAEQKX|mrna_SSu.010G0197700
Salix_brachista_(Salicaceae/Malpighiales)_rna−gnl|WGS_VDCV|Sabra10G0192800.1
Populus_tremula_(Salicaceae/Malpighiales)_Potra2n10c20183
Populus_trichocarpa_(Salicaceae/Malpighiales)_Potri.010G247400.1.v3.1
Populus_simonii_(Salicaceae/Malpighiales)_g33483.t1
Populus_deltoides_(Salicaceae/Malpighiales)_rna−gnl|WGS_JACEGQ|EVM20118.t1
Parasponia_andersonii_(Cannabaceae/Rosales)_PanWU01x14_157640.1
Eucalyptus_pauciflora_(Myrtaceae/Myrtales)_g23281.t1
Eucalyptus_grandis_(Myrtaceae/Myrtales)_gene−LOC104454049
Fagus_sylvatica_(Fagaceae/Fagales)_Bhaga_2.g2352.t1
Quercus_robur_(Fagaceae/Fagales)_Qrob_P0188130.2
Quercus_mongolica_(Fagaceae/Fagales)_g68740.t1
Quercus_suber_(Fagaceae/Fagales)_rna−CFP56_50227
Castanopsis_tibetana_(Fagaceae/Fagales)_evm.model.Contig165.29
Quercus_mongolica_(Fagaceae/Fagales)_g68743.t1
Quercus_robur_(Fagaceae/Fagales)_Qrob_P0188120.2
Juglans_regia_(Juglandaceae/Fagales)_Jr14_05860_p1
Morella_rubra_(Myricaceae/Fagales)_rna−gnl|WGS_RXIC|mrna.CJ030_MR1G018594
Casuarina_equisetifolia_(Casuarinaceae/Fagales)_Ceq08G1578
Casuarina_glauca_(Casuarinaceae/Fagales)_Cgl08G1608
Casuarina_cunninghamiana_(Casuarinaceae/Fagales)_Ccu08G1700
Betula_pendula_(Betulaceae/Fagales)_72777
Alnus_glutinosa_(Betulaceae/Fagales)_Alngl7889S36097
Carpinus_fangiana_(Betulaceae/Fagales)_rna−gnl|WGS_VIBQ|Cfa010436.t1
Corylus_heterophylla_(Betulaceae/Fagales)_g14900.t1
Corylus_avellana_(Betulaceae/Fagales)_g17794.t1
0.1

Fig. S19

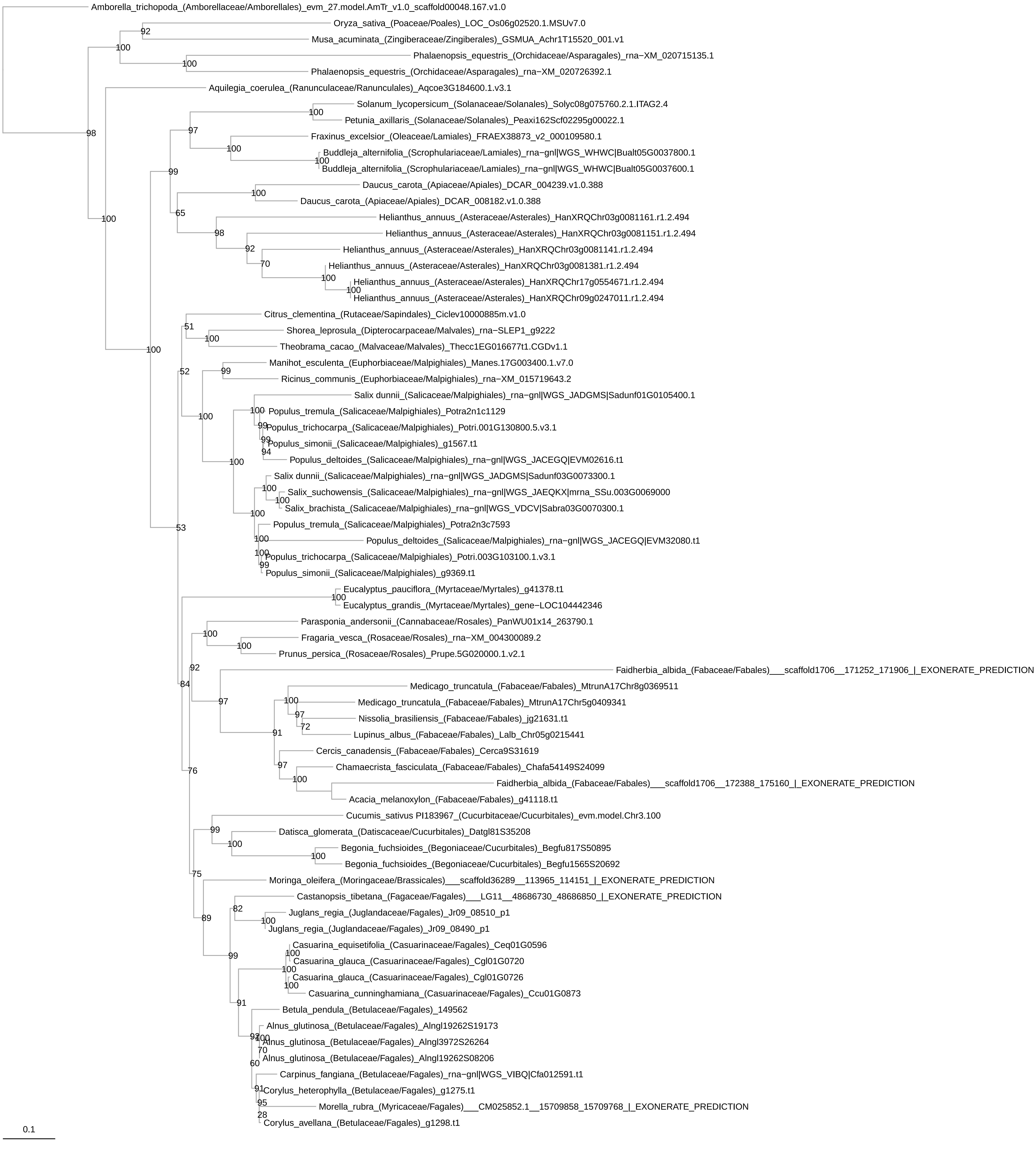

Amborella_trichopoda_(Amborellaceae/Amborellales)_evm_27.model.AmTr_v1.0_scaffold00048.167.v1.0
Oryza_sativa_(Poaceae/Poales)_LOC_Os06g02520.1.MSUv7.0
Musa_acuminata_(Zingiberaceae/Zingiberales)_GSMUA_Achr1T15520_001.v1
Phalaenopsis_equestris_(Orchidaceae/Asparagales)_rna−XM_020715135.1
Phalaenopsis_equestris_(Orchidaceae/Asparagales)_rna−XM_020726392.1
Aquilegia_coerulea_(Ranunculaceae/Ranunculales)_Aqcoe3G184600.1.v3.1
Solanum_lycopersicum_(Solanaceae/Solanales)_Solyc08g075760.2.1.ITAG2.4
Petunia_axillaris_(Solanaceae/Solanales)_Peaxi162Scf02295g00022.1
Fraxinus_excelsior_(Oleaceae/Lamiales)_FRAEX38873_v2_000109580.1
Buddleja_alternifolia_(Scrophulariaceae/Lamiales)_rna−gnl|WGS_WHWC|Bualt05G0037800.1
Buddleja_alternifolia_(Scrophulariaceae/Lamiales)_rna−gnl|WGS_WHWC|Bualt05G0037600.1
Daucus_carota_(Apiaceae/Apiales)_DCAR_004239.v1.0.388
Daucus_carota_(Apiaceae/Apiales)_DCAR_008182.v1.0.388
Helianthus_annuus_(Asteraceae/Asterales)_HanXRQChr03g0081161.r1.2.494
Helianthus_annuus_(Asteraceae/Asterales)_HanXRQChr03g0081151.r1.2.494
Helianthus_annuus_(Asteraceae/Asterales)_HanXRQChr03g0081141.r1.2.494
Helianthus_annuus_(Asteraceae/Asterales)_HanXRQChr03g0081381.r1.2.494
Helianthus_annuus_(Asteraceae/Asterales)_HanXRQChr17g0554671.r1.2.494
Helianthus_annuus_(Asteraceae/Asterales)_HanXRQChr09g0247011.r1.2.494
Citrus_clementina_(Rutaceae/Sapindales)_Ciclev10000885m.v1.0
Shorea_leprosula_(Dipterocarpaceae/Malvales)_rna−SLEP1_g9222
Theobrama_cacao_(Malvaceae/Malvales)_Thecc1EG016677t1.CGDv1.1
Manihot_esculenta_(Euphorbiaceae/Malpighiales)_Manes.17G003400.1.v7.0
Ricinus_communis_(Euphorbiaceae/Malpighiales)_rna−XM_015719643.2
Salix dunnii_(Salicaceae/Malpighiales)_rna−gnl|WGS_JADGMS|Sadunf01G0105400.1
Populus_tremula_(Salicaceae/Malpighiales)_Potra2n1c1129
Populus_trichocarpa_(Salicaceae/Malpighiales)_Potri.001G130800.5.v3.1
Populus_simonii_(Salicaceae/Malpighiales)_g1567.t1
Populus_deltoides_(Salicaceae/Malpighiales)_rna−gnl|WGS_JACEGQ|EVM02616.t1
Salix dunnii_(Salicaceae/Malpighiales)_rna−gnl|WGS_JADGMS|Sadunf03G0073300.1
Salix_suchowensis_(Salicaceae/Malpighiales)_rna−gnl|WGS_JAEQKX|mrna_SSu.003G0069000
Salix_brachista_(Salicaceae/Malpighiales)_rna−gnl|WGS_VDCV|Sabra03G0070300.1
Populus_tremula_(Salicaceae/Malpighiales)_Potra2n3c7593
Populus_deltoides_(Salicaceae/Malpighiales)_rna−gnl|WGS_JACEGQ|EVM32080.t1
Populus_trichocarpa_(Salicaceae/Malpighiales)_Potri.003G103100.1.v3.1
Populus_simonii_(Salicaceae/Malpighiales)_g9369.t1
Eucalyptus_pauciflora_(Myrtaceae/Myrtales)_g41378.t1
Eucalyptus_grandis_(Myrtaceae/Myrtales)_gene−LOC104442346
Parasponia_andersonii_(Cannabaceae/Rosales)_PanWU01x14_263790.1
Fragaria_vesca_(Rosaceae/Rosales)_rna−XM_004300089.2
Prunus_persica_(Rosaceae/Rosales)_Prupe.5G020000.1.v2.1
Faidherbia_albida_(Fabaceae/Fabales)___scaffold1706__171252_171906_|_EXONERATE_PREDICTION
Medicago_truncatula_(Fabaceae/Fabales)_MtrunA17Chr8g0369511
Medicago_truncatula_(Fabaceae/Fabales)_MtrunA17Chr5g0409341
Nissolia_brasiliensis_(Fabaceae/Fabales)_jg21631.t1
Lupinus_albus_(Fabaceae/Fabales)_Lalb_Chr05g0215441
Cercis_canadensis_(Fabaceae/Fabales)_Cerca9S31619
Chamaecrista_fasciculata_(Fabaceae/Fabales)_Chafa54149S24099
Faidherbia_albida_(Fabaceae/Fabales)___scaffold1706__172388_175160_|_EXONERATE_PREDICTION
Acacia_melanoxylon_(Fabaceae/Fabales)_g41118.t1
Cucumis_sativus PI183967_(Cucurbitaceae/Cucurbitales)_evm.model.Chr3.100
Datisca_glomerata_(Datiscaceae/Cucurbitales)_Datgl81S35208
Begonia_fuchsioides_(Begoniaceae/Cucurbitales)_Begfu817S50895
Begonia_fuchsioides_(Begoniaceae/Cucurbitales)_Begfu1565S20692
Moringa_oleifera_(Moringaceae/Brassicales)___scaffold36289__113965_114151_|_EXONERATE_PREDICTION
Castanopsis_tibetana_(Fagaceae/Fagales)___LG11__48686730_48686850_|_EXONERATE_PREDICTION
Juglans_regia_(Juglandaceae/Fagales)_Jr09_08510_p1
Juglans_regia_(Juglandaceae/Fagales)_Jr09_08490_p1
Casuarina_equisetifolia_(Casuarinaceae/Fagales)_Ceq01G0596
Casuarina_glauca_(Casuarinaceae/Fagales)_Cgl01G0720
Casuarina_glauca_(Casuarinaceae/Fagales)_Cgl01G0726
Casuarina_cunninghamiana_(Casuarinaceae/Fagales)_Ccu01G0873
Betula_pendula_(Betulaceae/Fagales)_149562
Alnus_glutinosa_(Betulaceae/Fagales)_Alngl19262S19173
Alnus_glutinosa_(Betulaceae/Fagales)_Alngl3972S26264
Alnus_glutinosa_(Betulaceae/Fagales)_Alngl19262S08206
Carpinus_fangiana_(Betulaceae/Fagales)_rna−gnl|WGS_VIBQ|Cfa012591.t1
Corylus_heterophylla_(Betulaceae/Fagales)_g1275.t1
Morella_rubra_(Myricaceae/Fagales)___CM025852.1__15709858_15709768_|_EXONERATE_PREDICTION
Corylus_avellana_(Betulaceae/Fagales)_g1298.t1
0.1

Fig. S20

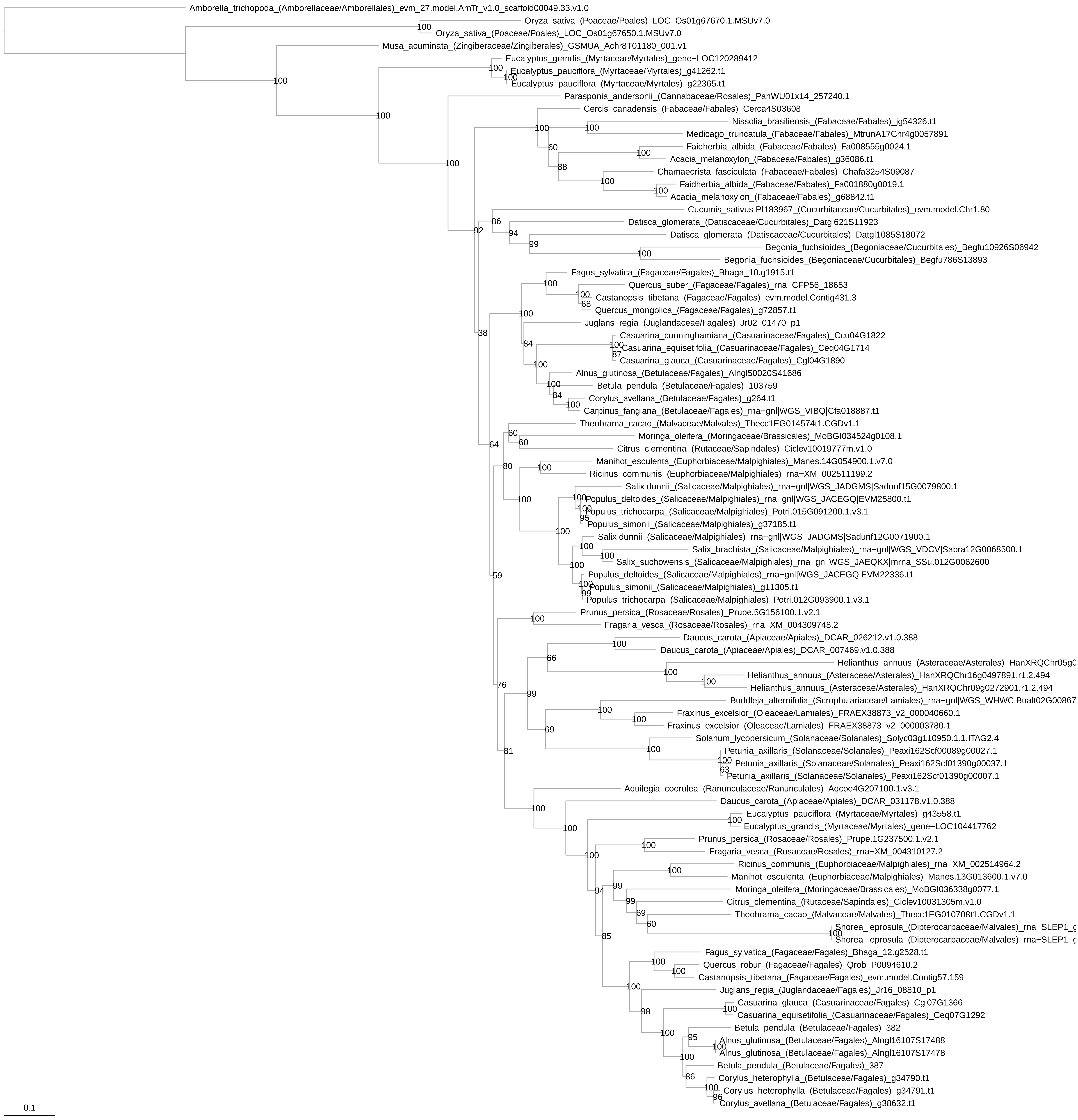

Amborella_trichopoda_(Amborellaceae/Amborellales)_evm_27.model.AmTr_v1.0_scaffold00049.33.v1.0
Oryza_sativa_(Poaceae/Poales)_LOC_Os01g67670.1.MSUv7.0
Oryza_sativa_(Poaceae/Poales)_LOC_Os01g67650.1.MSUv7.0
Musa_acuminata_(Zingiberaceae/Zingiberales)_GSMUA_Achr8T01180_001.v1
Eucalyptus_grandis_(Myrtaceae/Myrtales)_gene−LOC120289412
Eucalyptus_pauciflora_(Myrtaceae/Myrtales)_g41262.t1
Eucalyptus_pauciflora_(Myrtaceae/Myrtales)_g22365.t1
Parasponia_andersonii_(Cannabaceae/Rosales)_PanWU01x14_257240.1
Cercis_canadensis_(Fabaceae/Fabales)_Cerca4S03608
Nissolia_brasiliensis_(Fabaceae/Fabales)_jg54326.t1
Medicago_truncatula_(Fabaceae/Fabales)_MtrunA17Chr4g0057891
Faidherbia_albida_(Fabaceae/Fabales)_Fa008555g0024.1
Acacia_melanoxylon_(Fabaceae/Fabales)_g36086.t1
Chamaecrista_fasciculata_(Fabaceae/Fabales)_Chafa3254S09087
Faidherbia_albida_(Fabaceae/Fabales)_Fa001880g0019.1
Acacia_melanoxylon_(Fabaceae/Fabales)_g68842.t1
Cucumis_sativus PI183967_(Cucurbitaceae/Cucurbitales)_evm.model.Chr1.80
Datisca_glomerata_(Datiscaceae/Cucurbitales)_Datgl621S11923
Datisca_glomerata_(Datiscaceae/Cucurbitales)_Datgl1085S18072
Begonia_fuchsioides_(Begoniaceae/Cucurbitales)_Begfu10926S06942
Begonia_fuchsioides_(Begoniaceae/Cucurbitales)_Begfu786S13893
Fagus_sylvatica_(Fagaceae/Fagales)_Bhaga_10.g1915.t1
Quercus_suber_(Fagaceae/Fagales)_rna−CFP56_18653
Castanopsis_tibetana_(Fagaceae/Fagales)_evm.model.Contig431.3
Quercus_mongolica_(Fagaceae/Fagales)_g72857.t1
Juglans_regia_(Juglandaceae/Fagales)_Jr02_01470_p1
Casuarina_cunninghamiana_(Casuarinaceae/Fagales)_Ccu04G1822
Casuarina_equisetifolia_(Casuarinaceae/Fagales)_Ceq04G1714
Casuarina_glauca_(Casuarinaceae/Fagales)_Cgl04G1890
Alnus_glutinosa_(Betulaceae/Fagales)_Alngl50020S41686
Betula_pendula_(Betulaceae/Fagales)_103759
Corylus_avellana_(Betulaceae/Fagales)_g264.t1
Carpinus_fangiana_(Betulaceae/Fagales)_rna−gnl|WGS_VIBQ|Cfa018887.t1
Theobrama_cacao_(Malvaceae/Malvales)_Thecc1EG014574t1.CGDv1.1
Moringa_oleifera_(Moringaceae/Brassicales)_MoBGI034524g0108.1
Citrus_clementina_(Rutaceae/Sapindales)_Ciclev10019777m.v1.0
Manihot_esculenta_(Euphorbiaceae/Malpighiales)_Manes.14G054900.1.v7.0
Ricinus_communis_(Euphorbiaceae/Malpighiales)_rna−XM_002511199.2
Salix dunnii_(Salicaceae/Malpighiales)_rna−gnl|WGS_JADGMS|Sadunf15G0079800.1
Populus_deltoides_(Salicaceae/Malpighiales)_rna−gnl|WGS_JACEGQ|EVM25800.t1
Populus_trichocarpa_(Salicaceae/Malpighiales)_Potri.015G091200.1.v3.1
Populus_simonii_(Salicaceae/Malpighiales)_g37185.t1
Salix dunnii_(Salicaceae/Malpighiales)_rna−gnl|WGS_JADGMS|Sadunf12G0071900.1
Salix_brachista_(Salicaceae/Malpighiales)_rna−gnl|WGS_VDCV|Sabra12G0068500.1
Salix_suchowensis_(Salicaceae/Malpighiales)_rna−gnl|WGS_JAEQKX|mrna_SSu.012G0062600
Populus_deltoides_(Salicaceae/Malpighiales)_rna−gnl|WGS_JACEGQ|EVM22336.t1
Populus_simonii_(Salicaceae/Malpighiales)_g11305.t1
Populus_trichocarpa_(Salicaceae/Malpighiales)_Potri.012G093900.1.v3.1
Prunus_persica_(Rosaceae/Rosales)_Prupe.5G156100.1.v2.1
Fragaria_vesca_(Rosaceae/Rosales)_rna−XM_004309748.2
Daucus_carota_(Apiaceae/Apiales)_DCAR_026212.v1.0.388
Daucus_carota_(Apiaceae/Apiales)_DCAR_007469.v1.0.388
Helianthus_annuus_(Asteraceae/Asterales)_HanXRQChr05g0
Helianthus_annuus_(Asteraceae/Asterales)_HanXRQChr16g0497891.r1.2.494
Helianthus_annuus_(Asteraceae/Asterales)_HanXRQChr09g0272901.r1.2.494
Buddleja_alternifolia_(Scrophulariaceae/Lamiales)_rna−gnl|WGS_WHWC|Bualt02G00867
Fraxinus_excelsior_(Oleaceae/Lamiales)_FRAEX38873_v2_000040660.1
Fraxinus_excelsior_(Oleaceae/Lamiales)_FRAEX38873_v2_000003780.1
Solanum_lycopersicum_(Solanaceae/Solanales)_Solyc03g110950.1.1.ITAG2.4
Petunia_axillaris_(Solanaceae/Solanales)_Peaxi162Scf00089g00027.1
Petunia_axillaris_(Solanaceae/Solanales)_Peaxi162Scf01390g00037.1
Petunia_axillaris_(Solanaceae/Solanales)_Peaxi162Scf01390g00007.1
Aquilegia_coerulea_(Ranunculaceae/Ranunculales)_Aqcoe4G207100.1.v3.1
Daucus_carota_(Apiaceae/Apiales)_DCAR_031178.v1.0.388
Eucalyptus_pauciflora_(Myrtaceae/Myrtales)_g43558.t1
Eucalyptus_grandis_(Myrtaceae/Myrtales)_gene−LOC104417762
Prunus_persica_(Rosaceae/Rosales)_Prupe.1G237500.1.v2.1
Fragaria_vesca_(Rosaceae/Rosales)_rna−XM_004310127.2
Ricinus_communis_(Euphorbiaceae/Malpighiales)_rna−XM_002514964.2
Manihot_esculenta_(Euphorbiaceae/Malpighiales)_Manes.13G013600.1.v7.0
Moringa_oleifera_(Moringaceae/Brassicales)_MoBGI036338g0077.1
Citrus_clementina_(Rutaceae/Sapindales)_Ciclev10031305m.v1.0
Theobrama_cacao_(Malvaceae/Malvales)_Thecc1EG010708t1.CGDv1.1
Shorea_leprosula_(Dipterocarpaceae/Malvales)_rna−SLEP1_g
Shorea_leprosula_(Dipterocarpaceae/Malvales)_rna−SLEP1_g
Fagus_sylvatica_(Fagaceae/Fagales)_Bhaga_12.g2528.t1
Quercus_robur_(Fagaceae/Fagales)_Qrob_P0094610.2
Castanopsis_tibetana_(Fagaceae/Fagales)_evm.model.Contig57.159
Juglans_regia_(Juglandaceae/Fagales)_Jr16_08810_p1
Casuarina_glauca_(Casuarinaceae/Fagales)_Cgl07G1366
Casuarina_equisetifolia_(Casuarinaceae/Fagales)_Ceq07G1292
Betula_pendula_(Betulaceae/Fagales)_382
Alnus_glutinosa_(Betulaceae/Fagales)_Alngl16107S17488
Alnus_glutinosa_(Betulaceae/Fagales)_Alngl16107S17478
Betula_pendula_(Betulaceae/Fagales)_387
Corylus_heterophylla_(Betulaceae/Fagales)_g34790.t1
Corylus_heterophylla_(Betulaceae/Fagales)_g34791.t1
Corylus_avellana_(Betulaceae/Fagales)_g38632.t1
0.1

Fig. S21

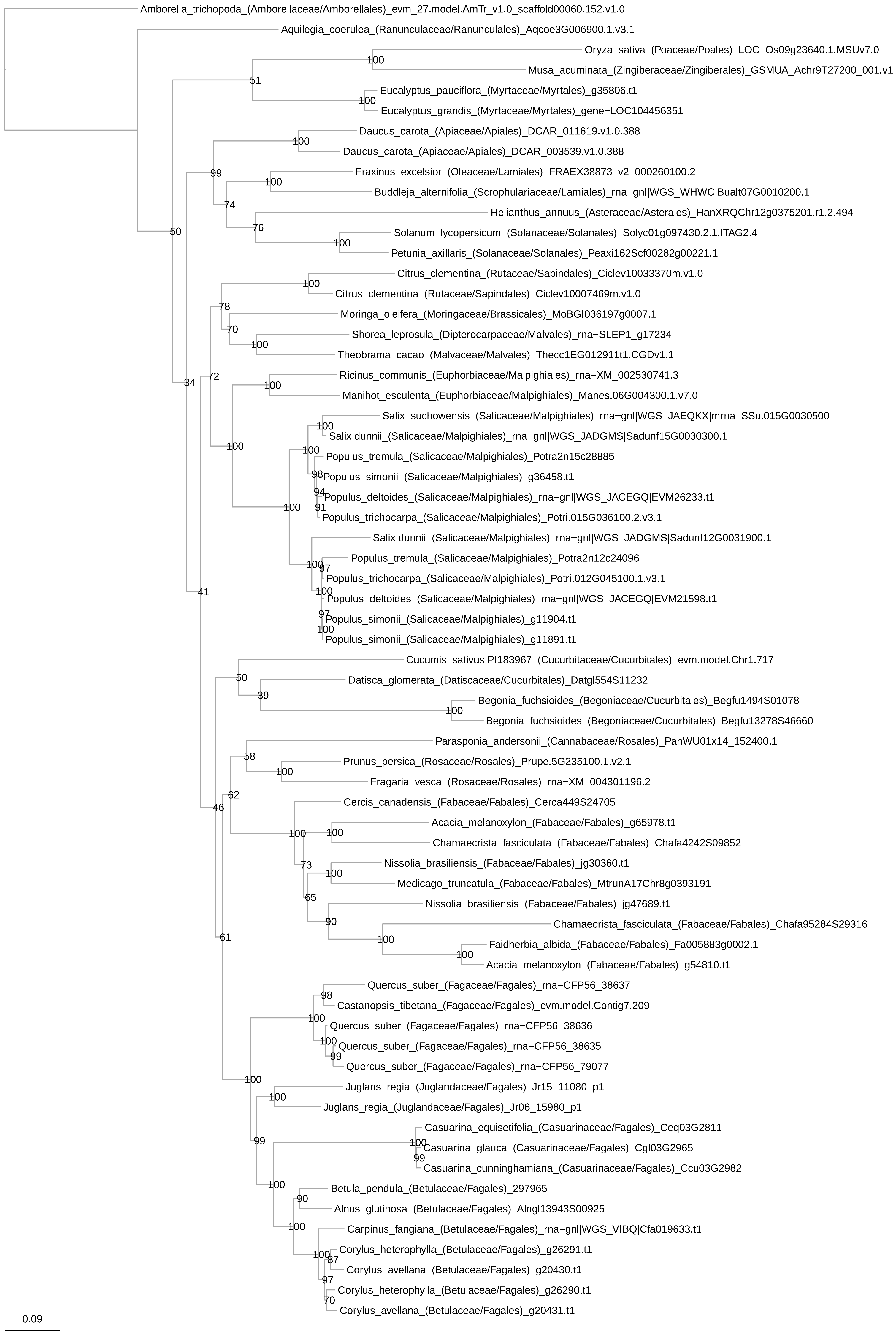

Amborella_trichopoda_(Amborellaceae/Amborellales)_evm_27.model.AmTr_v1.0_scaffold00060.152.v1.0
Aquilegia_coerulea_(Ranunculaceae/Ranunculales)_Aqcoe3G006900.1.v3.1
Oryza_sativa_(Poaceae/Poales)_LOC_Os09g23640.1.MSUv7.0
Musa_acuminata_(Zingiberaceae/Zingiberales)_GSMUA_Achr9T27200_001.v1
Eucalyptus_pauciflora_(Myrtaceae/Myrtales)_g35806.t1
Eucalyptus_grandis_(Myrtaceae/Myrtales)_gene−LOC104456351
Daucus_carota_(Apiaceae/Apiales)_DCAR_011619.v1.0.388
Daucus_carota_(Apiaceae/Apiales)_DCAR_003539.v1.0.388
Fraxinus_excelsior_(Oleaceae/Lamiales)_FRAEX38873_v2_000260100.2
Buddleja_alternifolia_(Scrophulariaceae/Lamiales)_rna−gnl|WGS_WHWC|Bualt07G0010200.1
Helianthus_annuus_(Asteraceae/Asterales)_HanXRQChr12g0375201.r1.2.494
Solanum_lycopersicum_(Solanaceae/Solanales)_Solyc01g097430.2.1.ITAG2.4
Petunia_axillaris_(Solanaceae/Solanales)_Peaxi162Scf00282g00221.1
Citrus_clementina_(Rutaceae/Sapindales)_Ciclev10033370m.v1.0
Citrus_clementina_(Rutaceae/Sapindales)_Ciclev10007469m.v1.0
Moringa_oleifera_(Moringaceae/Brassicales)_MoBGI036197g0007.1
Shorea_leprosula_(Dipterocarpaceae/Malvales)_rna−SLEP1_g17234
Theobrama_cacao_(Malvaceae/Malvales)_Thecc1EG012911t1.CGDv1.1
Ricinus_communis_(Euphorbiaceae/Malpighiales)_rna−XM_002530741.3
Manihot_esculenta_(Euphorbiaceae/Malpighiales)_Manes.06G004300.1.v7.0
Salix_suchowensis_(Salicaceae/Malpighiales)_rna−gnl|WGS_JAEQKX|mrna_SSu.015G0030500
Salix dunnii_(Salicaceae/Malpighiales)_rna−gnl|WGS_JADGMS|Sadunf15G0030300.1
Populus_tremula_(Salicaceae/Malpighiales)_Potra2n15c28885
Populus_simonii_(Salicaceae/Malpighiales)_g36458.t1
Populus_deltoides_(Salicaceae/Malpighiales)_rna−gnl|WGS_JACEGQ|EVM26233.t1
Populus_trichocarpa_(Salicaceae/Malpighiales)_Potri.015G036100.2.v3.1
Salix dunnii_(Salicaceae/Malpighiales)_rna−gnl|WGS_JADGMS|Sadunf12G0031900.1
Populus_tremula_(Salicaceae/Malpighiales)_Potra2n12c24096
Populus_trichocarpa_(Salicaceae/Malpighiales)_Potri.012G045100.1.v3.1
Populus_deltoides_(Salicaceae/Malpighiales)_rna−gnl|WGS_JACEGQ|EVM21598.t1
Populus_simonii_(Salicaceae/Malpighiales)_g11904.t1
Populus_simonii_(Salicaceae/Malpighiales)_g11891.t1
Cucumis_sativus PI183967_(Cucurbitaceae/Cucurbitales)_evm.model.Chr1.717
Datisca_glomerata_(Datiscaceae/Cucurbitales)_Datgl554S11232
Begonia_fuchsioides_(Begoniaceae/Cucurbitales)_Begfu1494S01078
Begonia_fuchsioides_(Begoniaceae/Cucurbitales)_Begfu13278S46660
Parasponia_andersonii_(Cannabaceae/Rosales)_PanWU01x14_152400.1
Prunus_persica_(Rosaceae/Rosales)_Prupe.5G235100.1.v2.1
Fragaria_vesca_(Rosaceae/Rosales)_rna−XM_004301196.2
Cercis_canadensis_(Fabaceae/Fabales)_Cerca449S24705
Acacia_melanoxylon_(Fabaceae/Fabales)_g65978.t1
Chamaecrista_fasciculata_(Fabaceae/Fabales)_Chafa4242S09852
Nissolia_brasiliensis_(Fabaceae/Fabales)_jg30360.t1
Medicago_truncatula_(Fabaceae/Fabales)_MtrunA17Chr8g0393191
Nissolia_brasiliensis_(Fabaceae/Fabales)_jg47689.t1
Chamaecrista_fasciculata_(Fabaceae/Fabales)_Chafa95284S29316
Faidherbia_albida_(Fabaceae/Fabales)_Fa005883g0002.1
Acacia_melanoxylon_(Fabaceae/Fabales)_g54810.t1
Quercus_suber_(Fagaceae/Fagales)_rna−CFP56_38637
Castanopsis_tibetana_(Fagaceae/Fagales)_evm.model.Contig7.209
Quercus_suber_(Fagaceae/Fagales)_rna−CFP56_38636
Quercus_suber_(Fagaceae/Fagales)_rna−CFP56_38635
Quercus_suber_(Fagaceae/Fagales)_rna−CFP56_79077
Juglans_regia_(Juglandaceae/Fagales)_Jr15_11080_p1
Juglans_regia_(Juglandaceae/Fagales)_Jr06_15980_p1
Casuarina_equisetifolia_(Casuarinaceae/Fagales)_Ceq03G2811
Casuarina_glauca_(Casuarinaceae/Fagales)_Cgl03G2965
Casuarina_cunninghamiana_(Casuarinaceae/Fagales)_Ccu03G2982
Betula_pendula_(Betulaceae/Fagales)_297965
Alnus_glutinosa_(Betulaceae/Fagales)_Alngl13943S00925
Carpinus_fangiana_(Betulaceae/Fagales)_rna−gnl|WGS_VIBQ|Cfa019633.t1
Corylus_heterophylla_(Betulaceae/Fagales)_g26291.t1
Corylus_avellana_(Betulaceae/Fagales)_g20430.t1
Corylus_heterophylla_(Betulaceae/Fagales)_g26290.t1
Corylus_avellana_(Betulaceae/Fagales)_g20431.t1
0.09

Fig. S22

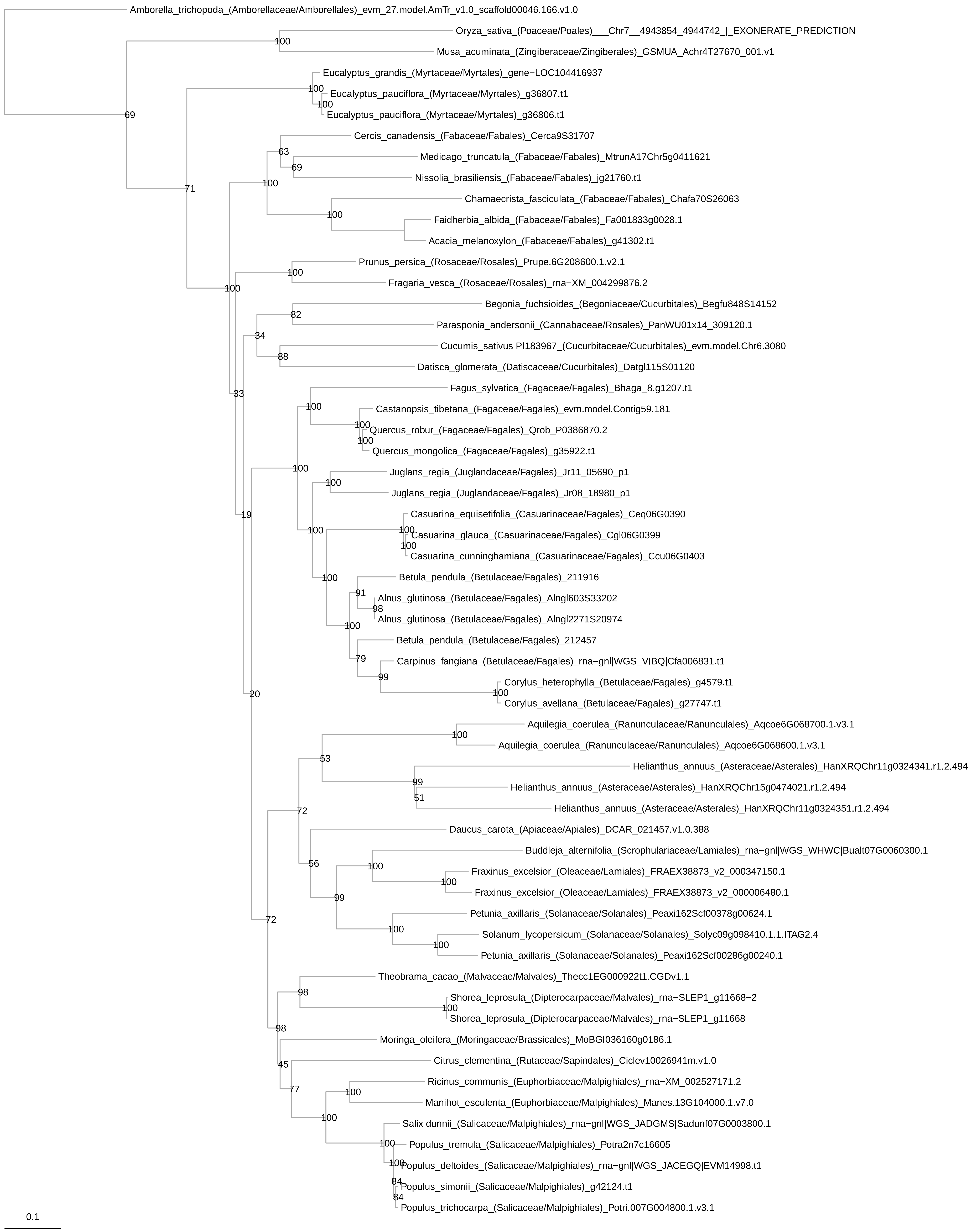
Amborella_trichopoda_(Amborellaceae/Amborellales)_evm_27.model.AmTr_v1.0_scaffold00046.166.v1.0
Oryza_sativa_(Poaceae/Poales)___Chr7__4943854_4944742_|_EXONERATE_PREDICTION
Musa_acuminata_(Zingiberaceae/Zingiberales)_GSMUA_Achr4T27670_001.v1
Eucalyptus_grandis_(Myrtaceae/Myrtales)_gene−LOC104416937
Eucalyptus_pauciflora_(Myrtaceae/Myrtales)_g36807.t1
Eucalyptus_pauciflora_(Myrtaceae/Myrtales)_g36806.t1
Cercis_canadensis_(Fabaceae/Fabales)_Cerca9S31707
Medicago_truncatula_(Fabaceae/Fabales)_MtrunA17Chr5g0411621
Nissolia_brasiliensis_(Fabaceae/Fabales)_jg21760.t1
Chamaecrista_fasciculata_(Fabaceae/Fabales)_Chafa70S26063
Faidherbia_albida_(Fabaceae/Fabales)_Fa001833g0028.1
Acacia_melanoxylon_(Fabaceae/Fabales)_g41302.t1
Prunus_persica_(Rosaceae/Rosales)_Prupe.6G208600.1.v2.1
Fragaria_vesca_(Rosaceae/Rosales)_rna−XM_004299876.2
Begonia_fuchsioides_(Begoniaceae/Cucurbitales)_Begfu848S14152
Parasponia_andersonii_(Cannabaceae/Rosales)_PanWU01x14_309120.1
Cucumis_sativus PI183967_(Cucurbitaceae/Cucurbitales)_evm.model.Chr6.3080
Datisca_glomerata_(Datiscaceae/Cucurbitales)_Datgl115S01120
Fagus_sylvatica_(Fagaceae/Fagales)_Bhaga_8.g1207.t1
Castanopsis_tibetana_(Fagaceae/Fagales)_evm.model.Contig59.181
Quercus_robur_(Fagaceae/Fagales)_Qrob_P0386870.2
Quercus_mongolica_(Fagaceae/Fagales)_g35922.t1
Juglans_regia_(Juglandaceae/Fagales)_Jr11_05690_p1
Juglans_regia_(Juglandaceae/Fagales)_Jr08_18980_p1
Casuarina_equisetifolia_(Casuarinaceae/Fagales)_Ceq06G0390
Casuarina_glauca_(Casuarinaceae/Fagales)_Cgl06G0399
Casuarina_cunninghamiana_(Casuarinaceae/Fagales)_Ccu06G0403
Betula_pendula_(Betulaceae/Fagales)_211916
Alnus_glutinosa_(Betulaceae/Fagales)_Alngl603S33202
Alnus_glutinosa_(Betulaceae/Fagales)_Alngl2271S20974
Betula_pendula_(Betulaceae/Fagales)_212457
Carpinus_fangiana_(Betulaceae/Fagales)_rna−gnl|WGS_VIBQ|Cfa006831.t1
Corylus_heterophylla_(Betulaceae/Fagales)_g4579.t1
Corylus_avellana_(Betulaceae/Fagales)_g27747.t1
Aquilegia_coerulea_(Ranunculaceae/Ranunculales)_Aqcoe6G068700.1.v3.1
Aquilegia_coerulea_(Ranunculaceae/Ranunculales)_Aqcoe6G068600.1.v3.1
Helianthus_annuus_(Asteraceae/Asterales)_HanXRQChr11g0324341.r1.2.494
Helianthus_annuus_(Asteraceae/Asterales)_HanXRQChr15g0474021.r1.2.494
Helianthus_annuus_(Asteraceae/Asterales)_HanXRQChr11g0324351.r1.2.494
Daucus_carota_(Apiaceae/Apiales)_DCAR_021457.v1.0.388
Buddleja_alternifolia_(Scrophulariaceae/Lamiales)_rna−gnl|WGS_WHWC|Bualt07G0060300.1
Fraxinus_excelsior_(Oleaceae/Lamiales)_FRAEX38873_v2_000347150.1
Fraxinus_excelsior_(Oleaceae/Lamiales)_FRAEX38873_v2_000006480.1
Petunia_axillaris_(Solanaceae/Solanales)_Peaxi162Scf00378g00624.1
Solanum_lycopersicum_(Solanaceae/Solanales)_Solyc09g098410.1.1.ITAG2.4
Petunia_axillaris_(Solanaceae/Solanales)_Peaxi162Scf00286g00240.1
Theobrama_cacao_(Malvaceae/Malvales)_Thecc1EG000922t1.CGDv1.1
Shorea_leprosula_(Dipterocarpaceae/Malvales)_rna−SLEP1_g11668−2
Shorea_leprosula_(Dipterocarpaceae/Malvales)_rna−SLEP1_g11668
Moringa_oleifera_(Moringaceae/Brassicales)_MoBGI036160g0186.1
Citrus_clementina_(Rutaceae/Sapindales)_Ciclev10026941m.v1.0
Ricinus_communis_(Euphorbiaceae/Malpighiales)_rna−XM_002527171.2
Manihot_esculenta_(Euphorbiaceae/Malpighiales)_Manes.13G104000.1.v7.0
Salix dunnii_(Salicaceae/Malpighiales)_rna−gnl|WGS_JADGMS|Sadunf07G0003800.1
Populus_tremula_(Salicaceae/Malpighiales)_Potra2n7c16605
Populus_deltoides_(Salicaceae/Malpighiales)_rna−gnl|WGS_JACEGQ|EVM14998.t1
Populus_simonii_(Salicaceae/Malpighiales)_g42124.t1
Populus_trichocarpa_(Salicaceae/Malpighiales)_Potri.007G004800.1.v3.1
0.1

Fig. S23

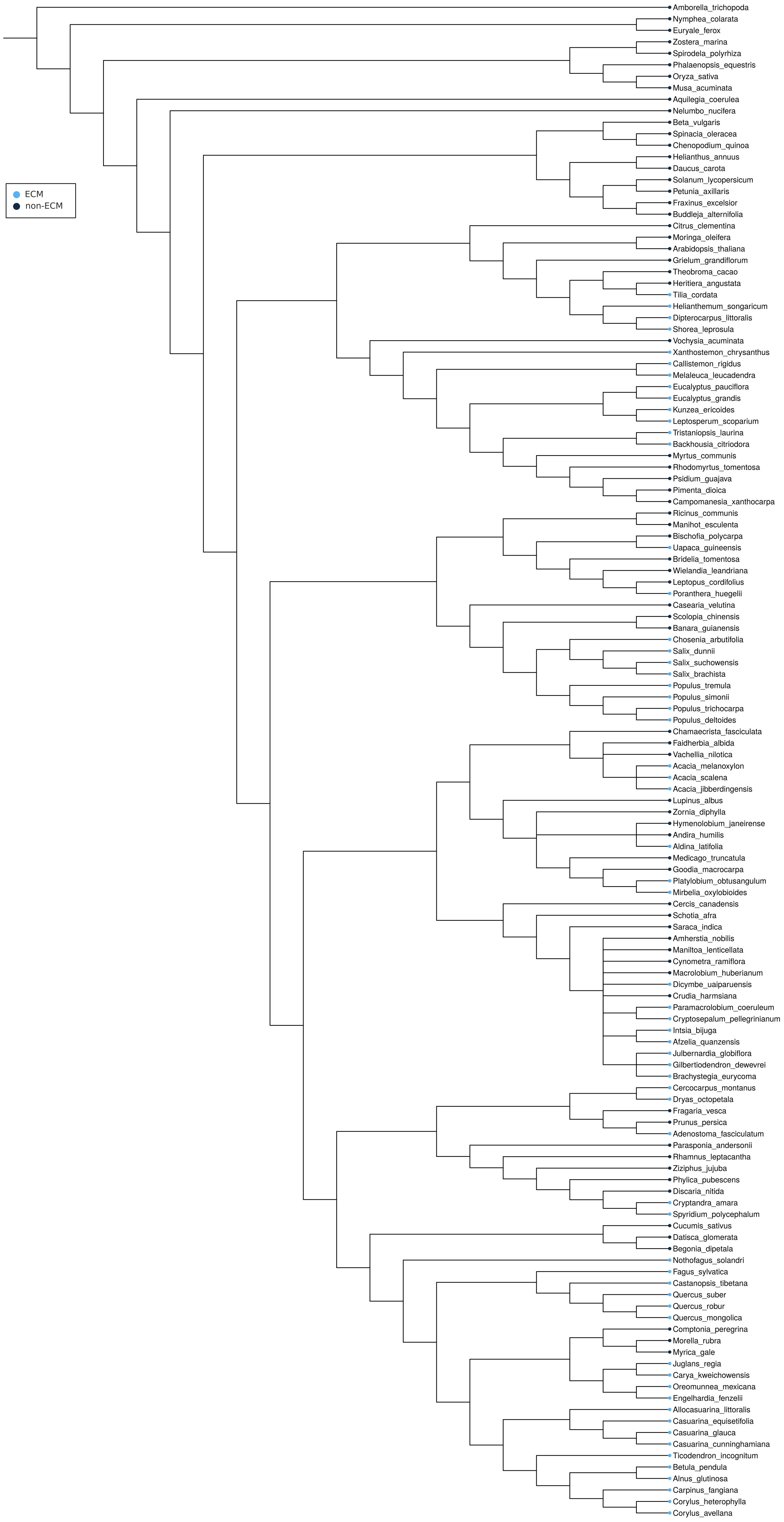
ECM
non-ECM
Amborella_trichopoda
Nymphea_colarata
Euryale_ferox
Zostera_marina
Spirodela_polyrhiza
Phalaenopsis_equestris
Oryza_sativa
Musa_acuminata
Aquilegia_coerulea
Nelumbo_nucifera
Beta_vulgaris
Spinacia_oleracea
Chenopodium_quinoa
Helianthus_annuus
Daucus_carota
Solanum_lycopersicum
Petunia_axillaris
Fraxinus_excelsior
Buddleja_alternifolia
Citrus_clementina
Moringa_oleifera
Arabidopsis_thaliana
Grielum_grandiflorum
Theobroma_cacao
Heritiera_angustata
Tilia_cordata
Helianthemum_songaricum
Dipterocarpus_littoralis
Shorea_leprosula
Vochysia_acuminata
Xanthostemon_chrysanthus
Callistemon_rigidus
Melaleuca_leucadendra
Eucalyptus_pauciflora
Eucalyptus_grandis
Kunzea_ericoides
Leptosperum_scoparium
Tristaniopsis_laurina
Backhousia_citriodora
Myrtus_communis
Rhodomyrtus_tomentosa
Psidium_guajava
Pimenta_dioica
Campomanesia_xanthocarpa
Ricinus_communis
Manihot_esculenta
Bischofia_polycarpa
Uapaca_guineensis
Bridelia_tomentosa
Wielandia_leandriana
Leptopus_cordifolius
Poranthera_huegelii
Casearia_velutina
Scolopia_chinensis
Banara_guianensis
Chosenia_arbutifolia
Salix_dunnii
Salix_suchowensis
Salix_brachista
Populus_tremula
Populus_simonii
Populus_trichocarpa
Populus_deltoides
Chamaecrista_fasciculata
Faidherbia_albida
Vachellia_nilotica
Acacia_melanoxylon
Acacia_scalena
Acacia_jibberdingensis
Lupinus_albus
Zornia_diphylla
Hymenolobium_janeirense
Andira_humilis
Aldina_latifolia
Medicago_truncatula
Goodia_macrocarpa
Platylobium_obtusangulum
Mirbelia_oxylobioides
Cercis_canadensis
Schotia_afra
Saraca_indica
Amherstia_nobilis
Maniltoa_lenticellata
Cynometra_ramiflora
Macrolobium_huberianum
Dicymbe_uaiparuensis
Crudia_harmsiana
Paramacrolobium_coeruleum
Cryptosepalum_pellegrinianum
Intsia_bijuga
Afzelia_quanzensis
Julbernardia_globiflora
Gilbertiodendron_dewevrei
Brachystegia_eurycoma
Cercocarpus_montanus
Dryas_octopetala
Fragaria_vesca
Prunus_persica
Adenostoma_fasciculatum
Parasponia_andersonii
Rhamnus_leptacantha
Ziziphus_jujuba
Phylica_pubescens
Discaria_nitida
Cryptandra_amara
Spyridium_polycephalum
Cucumis_sativus
Datisca_glomerata
Begonia_dipetala
Nothofagus_solandri
Fagus_sylvatica
Castanopsis_tibetana
Quercus_suber
Quercus_robur
Quercus_mongolica
Comptonia_peregrina
Morella_rubra
Myrica_gale
Juglans_regia
Carya_kweichowensis
Oreomunnea_mexicana
Engelhardia_fenzelii
Allocasuarina_littoralis
Casuarina_equisetifolia
Casuarina_glauca
Casuarina_cunninghamiana
Ticodendron_incognitum
Betula_pendula
Alnus_glutinosa
Carpinus_fangiana
Corylus_heterophylla
Corylus_avellana

Fig. S24

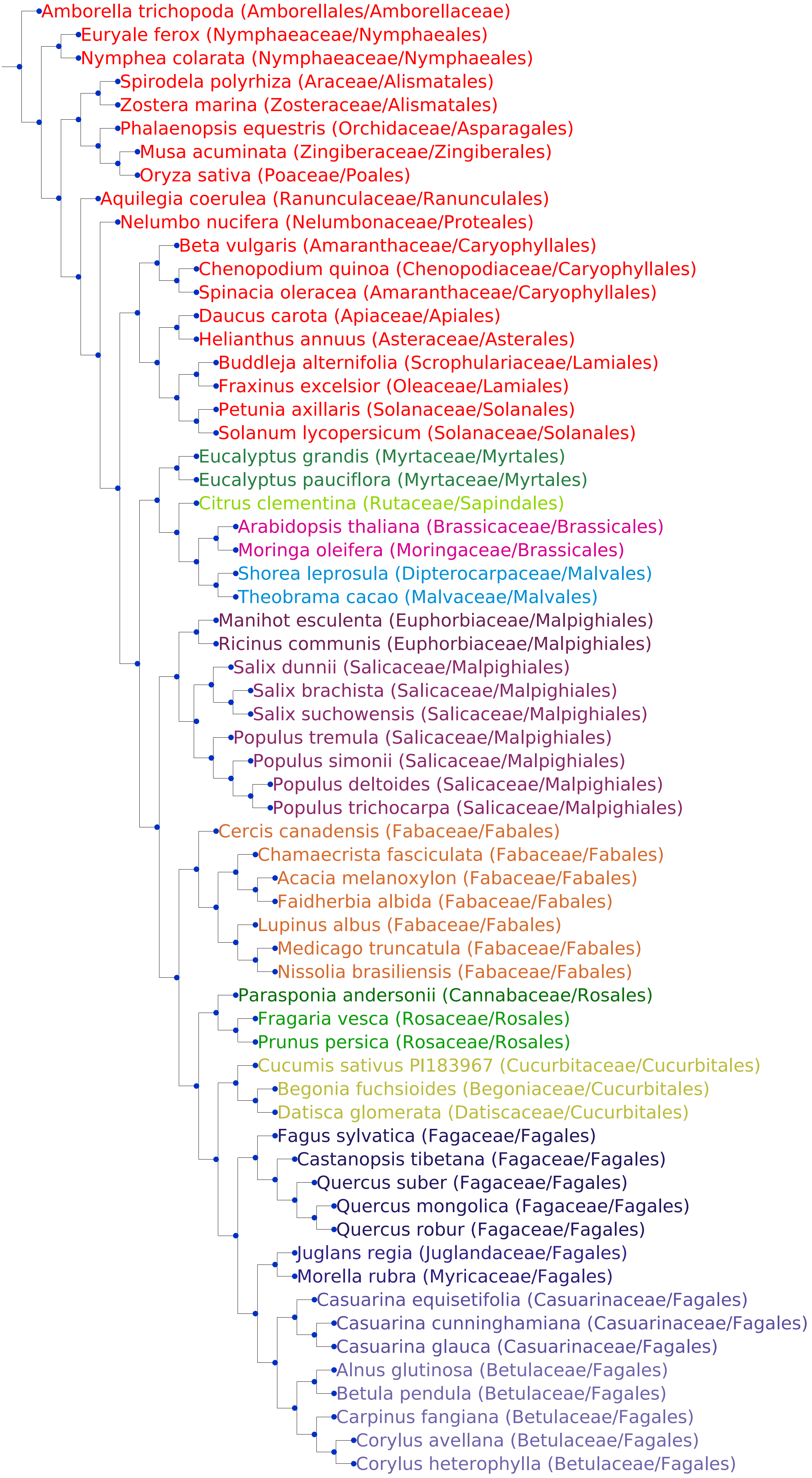
Amborella trichopoda (Amborellales/Amborellaceae)
Euryale ferox (Nymphaeaceae/Nymphaeales)
Nymphea colarata (Nymphaeaceae/Nymphaeales)
Spirodela polyrhiza (Araceae/Alismatales)
Zostera marina (Zosteraceae/Alismatales)
Phalaenopsis equestris (Orchidaceae/Asparagales)
Musa acuminata (Zingiberaceae/Zingiberales)
Oryza sativa (Poaceae/Poales)
Aquilegia coerulea (Ranunculaceae/Ranunculales)
Nelumbo nucifera (Nelumbonaceae/Proteales)
Beta vulgaris (Amaranthaceae/Caryophyllales)
Chenopodium quinoa (Chenopodiaceae/Caryophyllales)
Spinacia oleracea (Amaranthaceae/Caryophyllales)
Daucus carota (Apiaceae/Apiales)
Helianthus annuus (Asteraceae/Asterales)
Buddleja alternifolia (Scrophulariaceae/Lamiales)
Fraxinus excelsior (Oleaceae/Lamiales)
Petunia axillaris (Solanaceae/Solanales)
Solanum lycopersicum (Solanaceae/Solanales)
Eucalyptus grandis (Myrtaceae/Myrtales)
Eucalyptus pauciflora (Myrtaceae/Myrtales)
Citrus clementina (Rutaceae/Sapindales)
Arabidopsis thaliana (Brassicaceae/Brassicales)
Moringa oleifera (Moringaceae/Brassicales)
Shorea leprosula (Dipterocarpaceae/Malvales)
Theobrama cacao (Malvaceae/Malvales)
Manihot esculenta (Euphorbiaceae/Malpighiales)
Ricinus communis (Euphorbiaceae/Malpighiales)
Salix dunnii (Salicaceae/Malpighiales)
Salix brachista (Salicaceae/Malpighiales)
Salix suchowensis (Salicaceae/Malpighiales)
Populus tremula (Salicaceae/Malpighiales)
Populus simonii (Salicaceae/Malpighiales)
Populus deltoides (Salicaceae/Malpighiales)
Populus trichocarpa (Salicaceae/Malpighiales)
Cercis canadensis (Fabaceae/Fabales)
Chamaecrista fasciculata (Fabaceae/Fabales)
Acacia melanoxylon (Fabaceae/Fabales)
Faidherbia albida (Fabaceae/Fabales)
Lupinus albus (Fabaceae/Fabales)
Medicago truncatula (Fabaceae/Fabales)
Nissolia brasiliensis (Fabaceae/Fabales)
Parasponia andersonii (Cannabaceae/Rosales)
Fragaria vesca (Rosaceae/Rosales)
Prunus persica (Rosaceae/Rosales)
Cucumis sativus PI183967 (Cucurbitaceae/Cucurbitales)
Begonia fuchsioides (Begoniaceae/Cucurbitales)
Datisca glomerata (Datiscaceae/Cucurbitales)
Fagus sylvatica (Fagaceae/Fagales)
Castanopsis tibetana (Fagaceae/Fagales)
Quercus suber (Fagaceae/Fagales)
Quercus mongolica (Fagaceae/Fagales)
Quercus robur (Fagaceae/Fagales)
Juglans regia (Juglandaceae/Fagales)
Morella rubra (Myricaceae/Fagales)
Casuarina equisetifolia (Casuarinaceae/Fagales)
Casuarina cunninghamiana (Casuarinaceae/Fagales)
Casuarina glauca (Casuarinaceae/Fagales)
Alnus glutinosa (Betulaceae/Fagales)
Betula pendula (Betulaceae/Fagales)
Carpinus fangiana (Betulaceae/Fagales)
Corylus avellana (Betulaceae/Fagales)
Corylus heterophylla (Betulaceae/Fagales)

Fig. S25

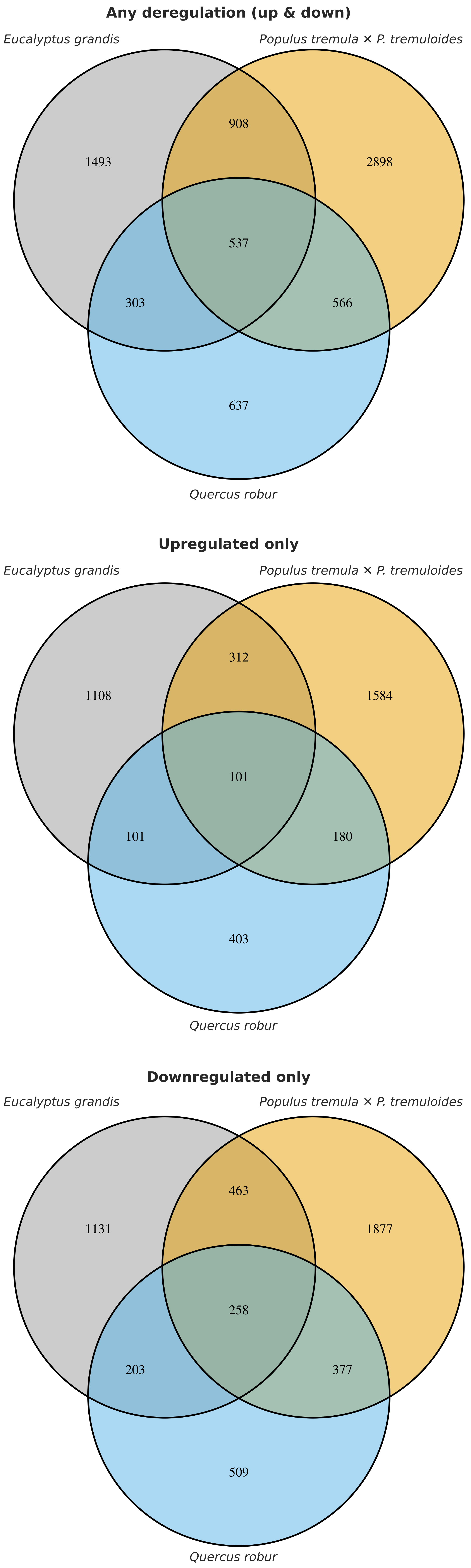
Any deregulation (up & down)
Eucalyptus grandis
Populus tremula × P. tremuloides
908
1493
2898
537
303
566
637
Quercus robur
Upregulated only
Eucalyptus grandis
Populus tremula × P. tremuloides
312
1108
1584
101
101
180
403
Quercus robur
Downregulated only
Eucalyptus grandis
Populus tremula × P. tremuloides
463
1131
1877
258
203
377
509
Quercus robur

Fig. S26

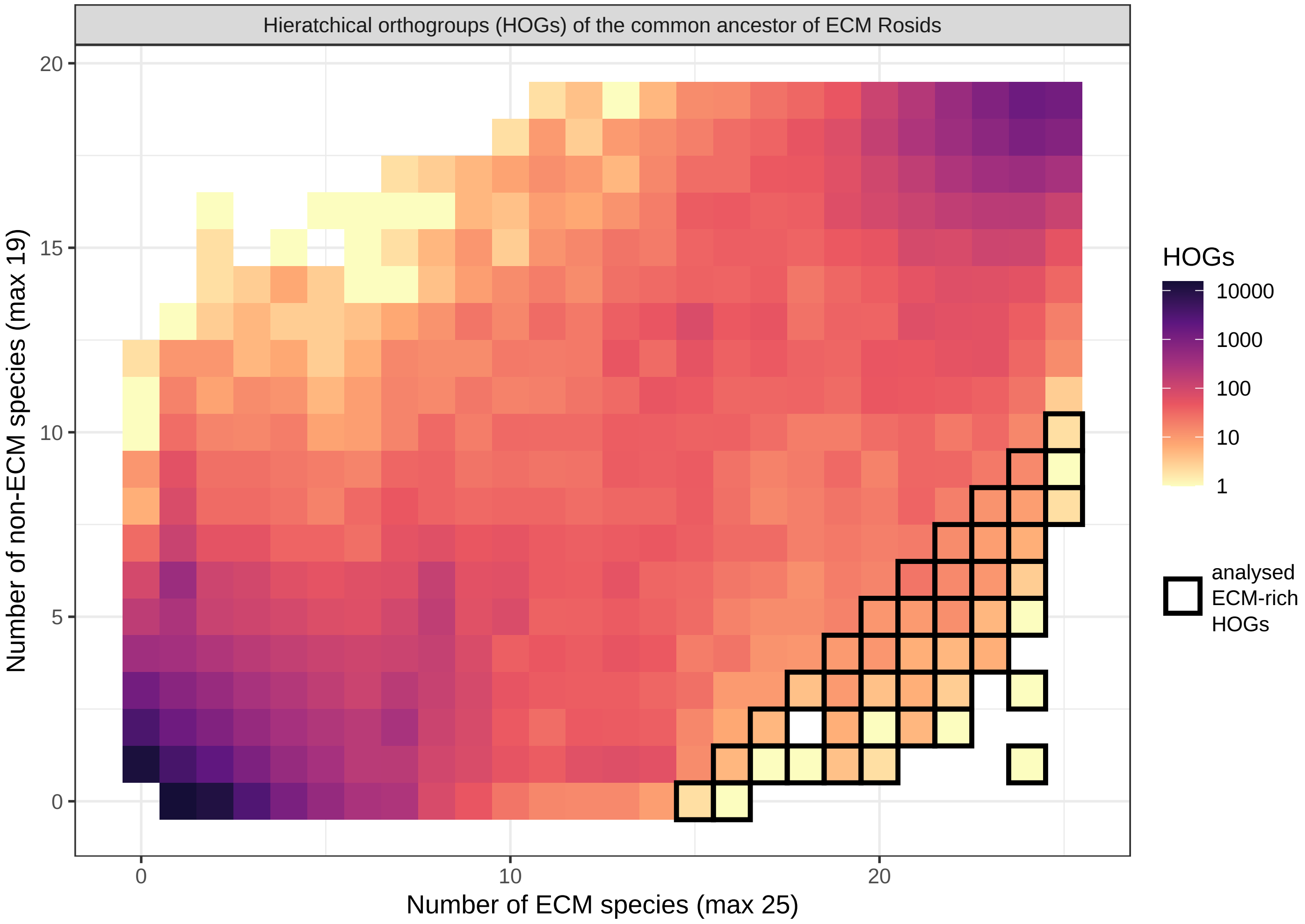

Hieratchical orthogroups (HOGs) of the common ancestor of ECM Rosids
Number of non-ECM species (max 19)
Number of ECM species (max 25)
0
5
10
15
20
0
10
20
HOGs
10000
1000
100
10
1
analysed
ECM-rich
HOGs
